# Water stress adaptive responses in plants require movement of ABA and AB-aldehyde from vascular to target tissues

**DOI:** 10.64898/2026.07.22.739638

**Authors:** Moran Anfang, Kristian B. Kiradjiev, Shir Ben Yaakov, Dan Gothilf, Christa Kanstrup, Barak Bitman, Jacob M Jepson, Liat Fellus-Alyagor, Dana Hirsch, Vadim E. Galperin, Shunsuke Watanabe, Masanori Okamoto, Roshan Kumar, Christoph Crocoll, Sara Morghan, Thorsten Hamann, Yariv Brotman, Mitsunori Seo, Hussam Hassan Nour-Eldin, Craig J. Sturrock, Poonam Mehra, Malcolm J. Bennett, James H. Rowe, Leah R Band, Eilon Shani

**Affiliations:** School of Plant Sciences and Food Security, Tel Aviv University, Tel Aviv, 69978, Israel; Centre for Mathematical Medicine and Biology, School of Mathematical Sciences, University of Nottingham, Nottingham NG7 2RD, UK; DynaMo Center, Department of Plant and Environmental Sciences, Faculty of Science, University of Copenhagen, 1871 Frederiksberg C, Denmark; Department of Veterinary Resources, Weizmann Institute of Science, Rehovot, Israel; Blavatnik Center for Drug Discovery, Tel Aviv University, Tel Aviv, 69978 Israel; RIKEN Center for Sustainable Resource Science, Yokohama, Kanagawa 230-0045, Japan; Department of Biological Science, Faculty of Science and Engineering, Yasuda Women’s University, Hiroshima, 731-0153, Japan; Tropical Biosphere Research Center, University of the Ryukyus, Nakagami-gun, Okinawa 903-0213, Japan; School of Biosciences, University of Nottingham, Sutton Bonington Campus, Loughborough LE12 5RD, UK; Department of Biology, Norwegian University of Science and Technology, Trondheim 7491, Norway; Plants, Photosynthesis and Soil, School of Biosciences, University of Sheffield, Sheffield, UK

## Abstract

Vascular plants rapidly coordinate root and shoot responses to water stress. Abscisic acid (ABA) mediates these adaptations; however, it remains unclear which cells produce ABA, whether ABA synthesis shifts during stress, and whether ABA movement is required for its adaptive functions. Here, we map ABA biosynthesis at cellular resolution in *Arabidopsis* and report that water-stress adaptive responses in roots and shoots require movement of ABA and its precursor AB-aldehyde from vascular tissues to target cells. We suggest that ABA accumulation arises from two parallel routes: (i) ABA synthesized in the vasculature via ABA2 and AAO3, then moving to guard cells, and (ii) phloem-derived AB-aldehyde being converted to ABA in the epidermis or bundle sheath by AAO1 and AAO2. Finally, we predict that tightly packed cells beneath leaf veins facilitate efficient ABA delivery to guard cells, an anatomical arrangement that has enabled angiosperms to evolve the use of ABA to rapidly close stomata.

## Introduction

Plants have evolved complex physiological and molecular strategies to cope with water scarcity. Among these strategies are the actions of abscisic acid (ABA), a key plant hormone (phytohormone) that regulates plant growth and development and orchestrates plant responses to environmental stresses ^1–6^. During water stress, ABA triggers a wide range of molecular responses across different tissues. In leaves, ABA acts on guard cells to control stomatal aperture, reducing water loss through transpiration. In the root, ABA promotes water foraging responses such as xerobranching ^6–12^, an adaptive response in which plants suppress lateral root formation in dry soil zones and prioritize root growth in moisture-rich areas ^6,13,14^.

Despite the central importance of ABA in plant water stress responses, the source of this signal under normal and stress conditions remains unclear. Early studies suggested that upon drought, the root senses the lack of water in the soil, and root-derived ABA is transported via xylem sap to the shoot for stomatal regulation ^15,16^. It was later found that ABA biosynthesis within the shoot can initiate stomatal closure ^9–12,17–22^. For example, *ABA2* and *AAO3* genes, encoding enzymes catalyzing the last two steps of the ABA pathway, were reported to be expressed in the shoot vasculature ^10^. *ABA2* encodes a short-chain dehydrogenase/reductase1 (SDR1) enzyme that makes abscisic aldehyde (AB-ald) from xanthoxin ^23^, while AAO3 (abscisic aldehyde oxidase 3) converts AB-ald to bioactive ABA ^24,25^. One of the fundamental and open questions in the field is whether ABA can be produced directly in guard cells in response to drought stress to regulate stomatal conductance, as studies have reported conflicting or context-dependent findings ^10,21^. For instance, Bauer et al. proposed that guard cell-autonomous ABA biosynthesis is essential for the stomatal response to reduced humidity, supporting the notion of local synthesis ^21^. In contrast, McAdam et al. suggested that mesophyll cells are the predominant site of water-deficit-triggered ABA biosynthesis ^22^. Furthermore, Kurmori et al. concluded that ABA is produced in phloem companion cells and can be transferred to guard cells ^10^. Regardless of the precise sites of ABA biosynthesis, ABA may need to move within and between tissues to reach its target cells. How this movement occurs remains poorly understood.

Studies have shown that during water stress, ABA accumulates at higher concentrations in leaves than in roots, and that its accumulation in roots can depend on basipetal (shoot-to-root) ABA transport from aerial organs ^9,26^. In addition, reciprocal grafting experiments between ABA-deficient mutants and wild-type plants have demonstrated that leaf-synthesized ABA affects stomatal closure^9,27^. If ABA is produced in the vasculature (phloem or xylem), it remains unclear whether it then moves through the apoplast, via plasmodesmata, by cell-to-cell transport, or through a combination of these mechanisms. Furthermore, it is not yet known whether the transported form is bioactive ABA, the conjugated form ABA-GE (abscisic acid glucose ester, an inactive ABA storage form), or ABA precursors such as AB-ald.

In roots, the epidermis is the ultimate site of action during xerobranching. When roots are exposed to water stress, ABA moves radially outwards with water from the vasculature to maintain organ growth and trigger closure of plasmodesmata (PD). PD closure prevents water loss plus inward movement of auxin, a hormone necessary for lateral root formation, effectively suppressing branching when external water is unavailable ^6,13,14^. This suggests that in roots, ABA, or ABA precursors, or ABA downstream signals, are transmitted between inner and outer tissues during water-stress responses; whether similar transport processes occur in leaves remains unresolved.

Several classes of ABA transporters have been identified in recent years ^26,28^. For example, ATP-binding cassette (ABC) family members function as ABA transporters in *Arabidopsis thaliana*: ABCG25 is an ABA exporter from the vasculature, while ABCG40 is an ABA importer to guard cells ^29,30^; AtABCG17 and AtABCG18 are plasma membrane ABA transporters that redundantly mediate ABA import in leaf mesophyll cells, forming inactive ABA sinks that limit stomatal closure and the pool of ABA that can travel to the root to regulate lateral root emergence. These transporters maintain ABA homeostasis under normal conditions, but their expression is repressed under abiotic stress, allowing free ABA to be released and promote rapid responses ^31,32^.

In this study, we provide comprehensive spatial and functional analyses of ABA biosynthesis and movement in roots and shoot tissues, uncovering key mechanisms through which ABA and its precursor, AB-ald, coordinate plant responses under water-stress conditions.

## Results

### Spatial transcriptomic and functional analysis of ABA biosynthesis during plant water stress responses

To reveal the spatial and temporal regulation of ABA biosynthesis during abiotic stress, we performed spatial transcriptomics on *Arabidopsis* leaves under watered and water-stress conditions, focusing on genes encoding the final 3 steps of ABA biosynthesis (**Fig. 1A**). In leaves, *NCED* genes (9-cis-epoxycarotenoid dioxygenase, encoding ABA biosynthesis enzymes localized to plastids ^2,33^), are expressed throughout the leaf; with *NCED5* predominantly epidermal; *NCED1,2* in palisade mesophyll (adaxial part); and *NCED3,9* in the vasculature (**Fig. 1B, Sup. Fig. S1**). The final two steps of ABA biosynthesis are catalyzed in the cytoplasm by enzymes encoded by the *ABA2* and *AAO* genes ^2,23–25^. *ABA2* expression was predominantly detected in the vasculature, alongside *AAO2* and *AAO3*, while *AAO1* and *AAO2* were also expressed in epidermis (**Fig. 1B, Sup. Fig. S2-3**). In addition, using *APL* ^34^ and *MYB46* ^35^ as markers for phloem companion cells and differentiating xylem cells, respectively, we assessed *ABA2* and *AAO3* expression within the vasculature. The spatial transcriptomics results suggest that, in addition to the phloem companion cells expression, *ABA2* and *AAO3* are detected in other phloem or cambium vascular tissues (**Sup. Fig. S3**). *ABA2* or *AAOs* expression was not detected in the cells associated with guard cells markers genes (**Sup. Fig. S4**), and we did not observe a change in *ABA2* and *AAOs* spatial expression under drought conditions (**Sup. Fig. S5-7**).

**Fig. 1.**
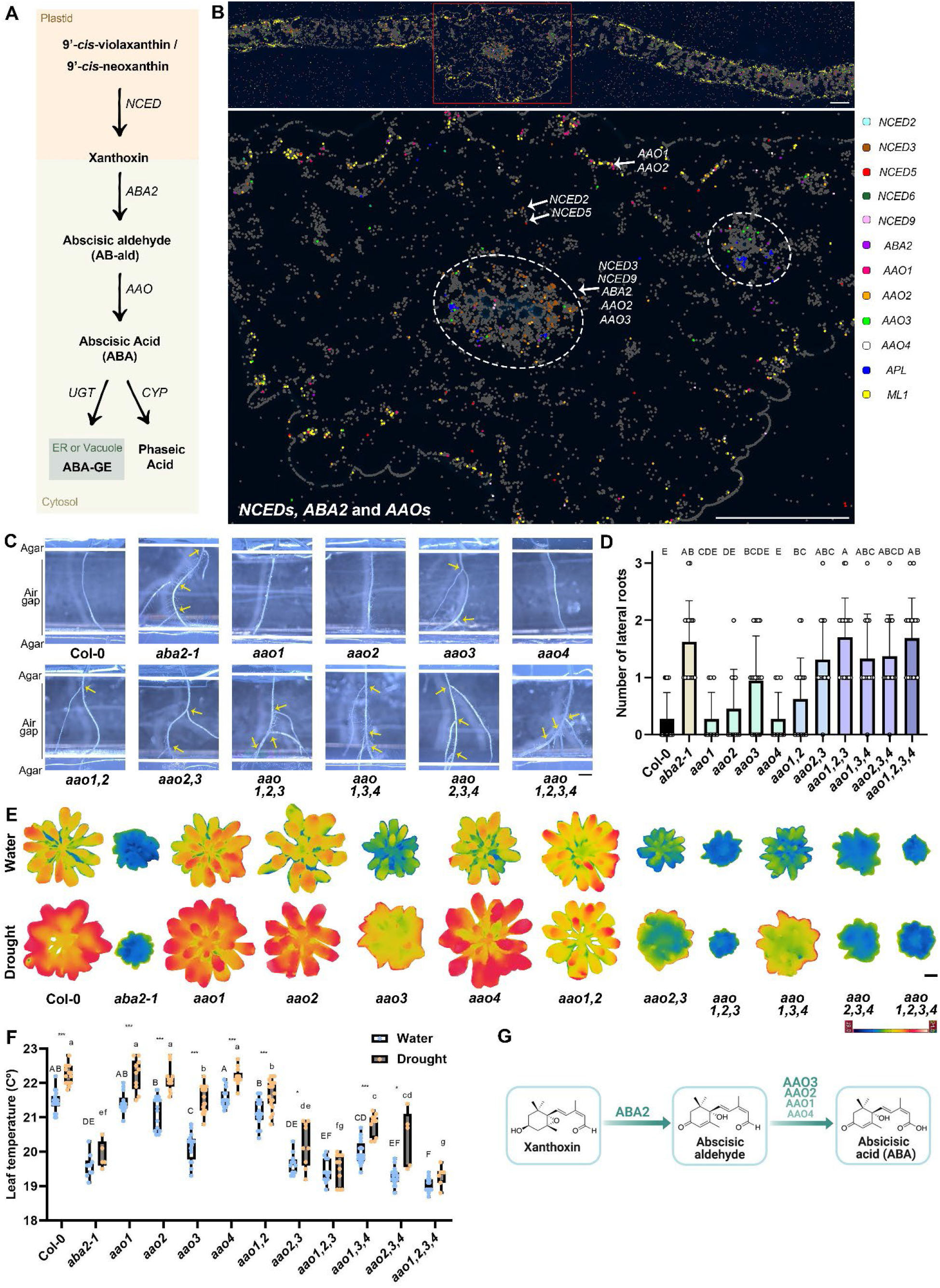
Redundant and robust activities of ABA biosynthesis genes in root and shoot water stress responses. **(A)** Illustration of the last steps in the ABA biosynthesis pathway. **(B)** Representative leaf spatial transcriptomics image showing the expression of ABA biosynthesis genes (top). The red rectangle indicates the region enlarged below, highlighting the spatial distribution of *NCED* genes, *ABA2*, and *AAO* genes. White dashed circles indicate vascular tissue. Scale bars = 50 µm. Genes written in white indicate the tissue in which they are expressed. Marker genes: *ML1* epidermis, *APL* phloem companion cells. **(C-D)** Shown are images **(C)** and quantification **(D)** of 13-day-old *aba2-1* and *aao* single, double, triple, and quadruple mutant combinations, with respective controls, in an agar-based air-gap assay system (Air gap = ∼5 mm). Mutant alleles: *aao1-2, aao2-1, aao3-4, aao4-2*. Yellow arrows indicate lateral roots. Scale bar = 1 mm. Significance was determined using Tukey’s ad-hoc statistical test (treatments marked with different letters are significantly different). n ≥ 11. **(E-F)** Shown are thermal images **(E)** and quantification **(F)** of *aba2-1* and *aao* mutant combinations with respective controls with and without drought stress. Scale bar = 1 cm. Significance was determined using Tukey’s ad-hoc statistical test (treatments marked with different letters are significantly different, capital letters represent significance between genotypes under water treatment, small letters represent significance between genotypes under drought treatment). Significance between watered and droughted plants from each genotype was determined using Student’s t-test, * = *p value < 0.05,* ** = *0.005 < p value < 0.01,* and *** = *p value < 0.005*. The *aao* quadruple mutant is a stable CRISPR line of *pRPS5:2sg-4AAOs* (Sup. Table 1). n ≥ 6. **(G)** Illustration of the last two steps in the ABA biosynthesis pathway. The gene names encoding the enzymes catalyzing each step are shown in green. Genes written in bold uppercase letters were found to play a more important role, while genes shown in lighter green and smaller font were considered less important for xerobranching and stomatal closure.

We next tested the functional importance of the final steps of ABA biosynthesis, ABA2, and AAOs activity mediating model root and shoot water stresses - root xerobranching ^36^ and shoot stomatal conductance ^17^. *ABA2* does not have homologs with the same biochemical activity in *Arabidopsis* ^37,38^, while the *AAO* family comprises four genes; among single mutants, only *aao3* shows a mild loss-of-function phenotype, and, to our knowledge, triple or quadruple AAO mutants have not been reported ^39^. To address this knowledge gap, we generated *aao* mutant combinations using CRISPR-Cas9 gene editing (**Sup. Table S1**) and T-DNA insertion lines.

We first performed a xerobranching bioassay ^6^ using the *aba2-1* mutant and single, double, triple, and quadruple mutants of *AAO* genes. The *aao3* single mutant exhibited a mild but significant increase in lateral root formation when growing across an air-gap compared to no lateral roots formed in wild-type, while other *aao* single mutants did not show a significant difference (**Fig. 1C-D**). Notably, double and higher-order combinations involving *aao3* (*aao2,3, aao1,2,3, aao1,3,4, aao2,3,4,* and the quadruple mutant *aao1,2,3,4*) displayed a stronger lateral root phenotype than *aao3* alone, similar to *aba2-1*, suggesting that *AAO* genes act redundantly to regulate ABA biosynthesis (**Fig. 1C-D**). Among them, *aao2* showed the most significant synergistic effect with *aao3*, while *aao4* combinations showed the weakest effect.

To assess the role of *AAO* genes mediating stomatal closure following drought stress, we performed a drought assay on single and triple *aao* mutant plants (**Fig. 1E-F**). While *aao1, aao2,* and *aao4* single mutants showed no discernible difference compared to wild type, *aao3* exhibited lower leaf temperatures under both watered and drought conditions, reflecting impaired stomatal closure and increased water loss. In both conditions, higher-order *aao* mutants containing *aao3* exhibited significantly lower leaf temperatures than in *aao3* alone, similar to those of the *aba2-1* mutant, confirming their redundant and important activity in the drought stress response (**Fig. 1E-F**). Our results reveal a complex spatial ABA biosynthetic map, where ABA2 likely controls this hormone pathway from the vasculature, and the final step, AAOs diversified over space, with AAO3 as the key regulator of xerobranching and stomatal closure, and AAO1 and AAO2 redundantly enhancing stomatal regulation and lateral root formation in water-limiting conditions (**Fig. 1G**).

### ABA biosynthesis bottleneck occurs in the phloem and not in guard cells

The spatial transcriptomics data reveal that *ABA2* and *AAO3* are not expressed throughout the leaf and are not transcriptionally induced by stress. The promoters of *ABA2* and *AAO3* were previously shown to be expressed in the shoot vasculature ^10^; however, the cloned promoters had not been functionally validated through complementation. To validate the functional activity of *ABA2* and *AAO3* promoters, we generated complementation lines for *ABA2* and *AAO3*. The selected promoters fully rescued *aba2-1* or *aao3-4* mutants when driving *ABA2* or *AAO3* translational fusions, respectively (**Fig. 2A-D**).

**Fig. 2.**
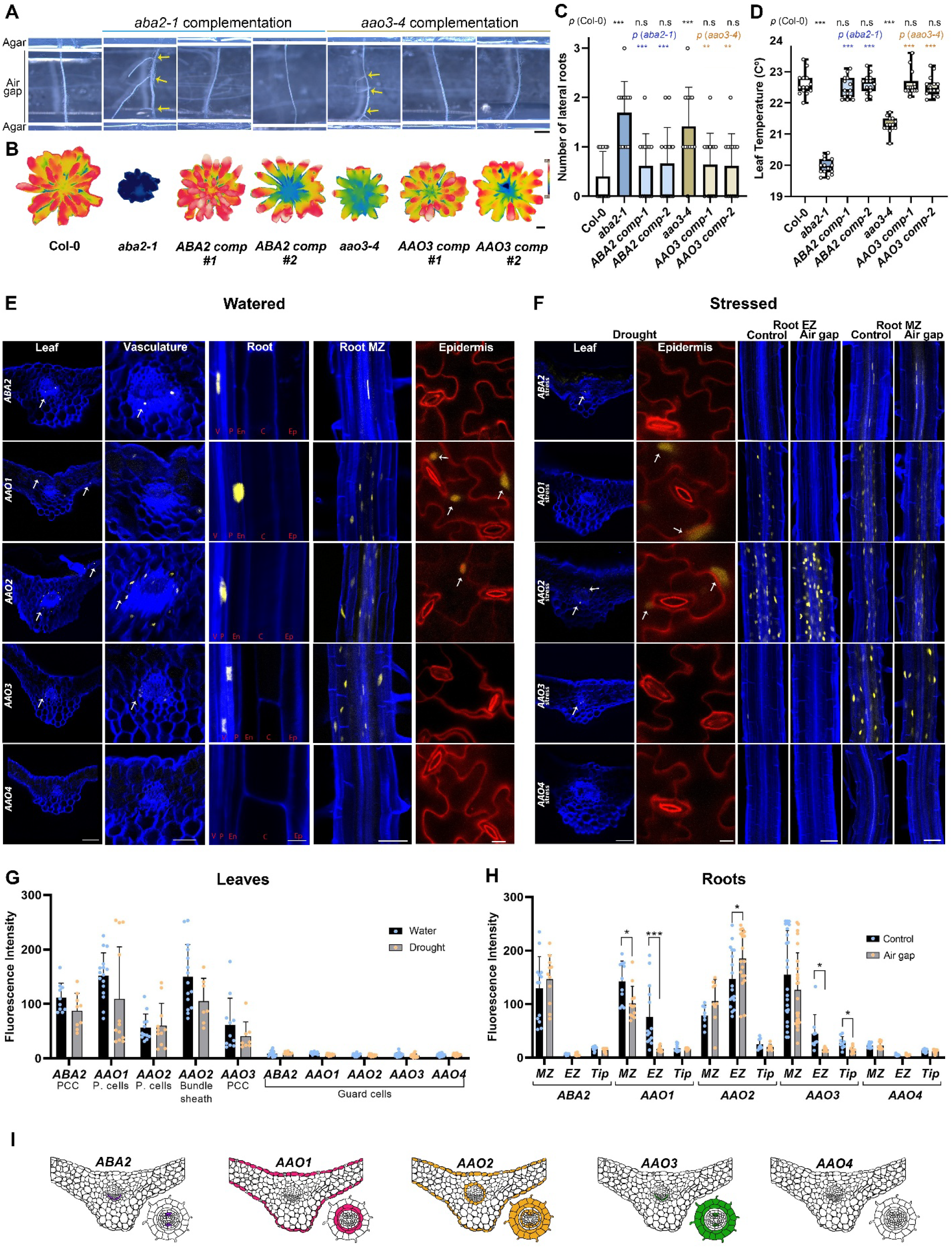
Abscisic-aldehyde biosynthesis is restricted to the phloem companion cells. **(A, C)** Representative root images **(A)** and quantification **(C)** of 13-day-old *aba2-1* and *aao3-4* complementation lines and respective controls in an agar-based air-gap assay system (Air gap = ∼5 mm). Yellow arrows indicate lateral roots. Scale bar = 1 mm. n ≥ 12, significance was determined using student’s t-test. ** = *0.005 < p value < 0.01,* and *** = *p value < 0.005*. **(B, D)** Representative thermal images **(B)** and quantification **(D)** of 45-day-old *aba2-1* and *aao3-4* complementation lines and respective controls grown under short-day conditions. Scale bar = 1 cm. n ≥ 4, Significance was determined using student’s t-test. *** = *p value < 0.005*. **(E)** *pABA2, pAAO1, pAAO2, pAAO3,* and *pAAO4:NLS-YFP* expression patterns under watered conditions in different tissues. Leaf - confocal images of cross-sections of 26-day-old plants expressing *pABA2* and *AAOs* NLS-YFP reporters (*pAAO2:NLS-YFP* is 25 days old). Scale bar = 100 μM. White arrows indicate phloem (*ABA2, AAO3*), pavement cells (*AAO1, AAO2*), and bundle sheath (*AAO2*) expression. Vasculature - enlargement of the leaf cross-section images. Scale bar = 200 μm. White arrows indicate phloem (*ABA2, AAO3*) and bundle sheath (*AAO2*) expression. Root - confocal images of 7-day-old roots expressing *pABA2* and *AAOs* NLS-YFP reporters. The images were taken in point 4, as marked in Sup. Figure S2. Scale bar = 10 μm. Root MZ - confocal images of 7-day-old roots expressing *pABA2* and *AAOs* NLS-YFP reporters. The images were taken in point 6, as marked in Sup. Figure S2. Scale bar = 10 μm. Epidermis - guard cells and pavement cells expression of *pABA2* and *AAOs* NLS-YFP reporters. Scale bar = 10 μm. White arrows indicate pavement cells (*AAO1, AAO2*) expression. **(F)** *pABA2, pAAO1, pAAO2, pAAO3,* and *pAAO4:NLS-YFP* expression patterns under stress conditions in different tissues. Leaf - confocal images of cross-sections of 26-day-old plants expressing *pABA2* and *AAOs* NLS-YFP reporters after drought stress. Scale bar = 100 μM. White arrows indicate phloem (*ABA2, AAO3*), and bundle sheath (*AAO2*) expression. Epidermis - guard cells and pavement cells expression of *pABA2* and *AAOs* NLS-YFP reporters after drought stress. Scale bar = 10 μm. White arrows indicate pavement cells (*AAO1, AAO2*) expression. Root EZ and Root MZ - confocal images of 6-day-old roots expressing *pABA2* and *AAOs* NLS-YFP reporters. Control plants were grown on MS plates. Roots were exposed to an air gap for ∼24-30 hours. The images were taken at the elongation zone (EZ) or in the maturation zone (MZ). Scale bar = 50 μm. **(G-H)** Quantification of *pABA2, pAAO1, pAAO2, pAAO3,* and *pAAO4:NLS-YFP* expression patterns in: G) 28-day-old plants under watered or drought conditions, n ≥ 7, and H) after xerobranching. Roots of 6-day-old plants were exposed to an air gap for ∼24-30 hours. Measurements were taken from MZ, maturation zone, EZ, elongation zone, and Tip, root tip. n ≥ 7. **(I)** Expression patterns illustration of *pABA2, pAAO1, pAAO2, pAAO3,* and *pAAO4* in roots and leaves.

Since the promoters of *ABA2* and *AAO3* fully complemented their respective *aba2-1* or *aao3-4* mutants, we reasoned that they are mimicking each gene’s native expression, and we therefore used the same promoters to generate *NLS-YFP* transcriptional reporter lines and characterized their expression pattern in leaf and root tissues. *ABA2* is expressed in phloem companion cells in both the root and shoot (**Fig. 2E, Sup Fig. S8, S9**), consistent with an earlier study ^10^. No *ABA2* expression was detected in guard cells or mesophyll (**Fig. 2E, Sup. Fig. S10, S11**). To further verify this important result, we also cloned *pABA2:GUS* and recorded similar vascular-restricted expression, with no detectable signal in guard cells, in either 7-day-old cotyledons or mature leaves (**Sup. Fig. S12-S14**). Since ABA acts on guard cells, yet *ABA2* expression is largely restricted to the phloem and mostly absent from guard cells, our results suggest that a signal downstream of ABA2 must move from the phloem to the guard cells.

To further validate the spatial transcriptomics results, we cloned transcriptional reporters for all four *AAO* enzymes: *pAAO1:NLS-YFP*, *pAAO2:NLS-YFP, pAAO3:NLS-YFP,* and *pAAO4:NLS-YFP*. Notably, *AAO3,* which was previously shown to have explicit expression in phloem companion cells ^10,40^, was observed to exhibit broader expression in the root, specifically also in the cortex and epidermis (**Fig. 2E, Sup Fig. S8, S9**). *AAO1, AAO2,* and *AAO4,* which have not been previously characterized, were not expressed in the shoot vasculature. In the leaves, *AAO1* showed expression in the epidermis, and *AAO2* was expressed in the shoot bundle sheath and epidermis (**Fig. 2E**). *AAO1* showed low expression in guard cells of 7-day-old cotyledons (**Sup. Fig. S12, S15**); however, there was no guard cell expression in the leaves starting from a young age (**Sup. Fig. S14**). In the root, *AAO1* was expressed in the endodermis and cortex tissues, while *AAO2* was expressed in the phloem companion cells, pericycle, cortex and epidermis (**Fig. 2E, Sup. Fig. S8, S9**). *AAO4* was not expressed in the root or shoot of 7 and 28-day-old plants (**Fig. 2E, Sup. Fig. S9, S13**). Similar to *ABA2*, none of the four *AAOs* showed expression in mesophyll and guard cells of mature leaves (**Fig. 2E, Sup. Fig. S10, S11, S14**). Notably, it is possible that expression takes place in some context at a basal level that is below the detection level of both the *NLS-YFP* lines and spatial transcriptional approaches. While *AAO4* expression could not be detected in vegetative shoots and roots, GUS staining revealed expression in reproductive tissues, specifically in the anthers and septum, together with other ABA biosynthesis genes: *ABA2* was expressed in the pollen grains and pedicel, and *AAO2* was expressed in the anther and pollen grains (**Sup. Fig. S16**). These results align with the observed lack of *AAO4* functional activity during water stress responses in vegetative shoots and roots (**Fig. 1**).

ABA biosynthesis has been reported to be upregulated in response to abiotic stresses ^9^. We therefore expected a rapid and strong induction of the final two steps of ABA biosynthesis over time and across tissues in roots and shoots. However, the expression patterns before and after drought stress in shoots and air-gap stress in roots did not increase or expand into additional cell types or tissues (**Fig. 2F**), consistent with previous reports ^23,25^. Quantification of the signal in leaves revealed no statistically significant difference in the expression of any of the five ABA biosynthesis genes before or after 10 days of drought (no irrigation) (**Fig. 2G, Sup. Fig. S17, S18**). Following air-gap exposure in roots, we observed a mild but significant reduction in the expression of *AAO1* and *AAO3*, and a very slight elevation of *AAO2* expression in the root elongation zone (**Fig. 2H**). We also conducted a salinity stress assay, in which the seedlings were exposed to 100 mM NaCl for varying periods. Imaging experiments in roots indicated that *ABA2* expression after 6 h of salt treatment was slightly higher only in mature roots and remained restricted to phloem companion cells. *AAOs* expression did not change following salt stress, except for *AAO2,* which showed lower fluorescence intensity after 24 h of salt treatment (**Sup. Fig. S19**). To further validate these results, we conducted a xerobranching RNA-seq experiment, with time points collected at different stages. We found little to no change in *ABA2* and *AAO*s expression during root xerobranching. In contrast, *NCED3*, *NCED*5, and *NCED*9 showed a rapid and significant increase in gene expression, which was quickly restored once the root contacted the moist agar (**Sup. Fig. S20**). Overall, *ABA2* and *AAOs* gene expression patterns remained largely stable across tissues before and after stress exposure, with *ABA2* confined largely to the phloem, and phloem surrounding cells, serving as a spatial bottleneck for ABA production in both shoots and roots (**Fig. 2I**).

### Multiscale modelling predicts a gradient in ABA away from vasculature

As *ABA2* expression is largely confined to phloem companion cells and *AAO* genes are not expressed in guard cells, ABA function in mediating stomata closure is likely non-cell-autonomous, requiring movement of an intermediate or ABA itself. In root tissues, we observe a similar phenomenon: *ABA2* expression is confined to phloem companion cells and is absent from the root epidermis, where it is required for xerobranching.

To understand how tissue-specific synthesis affects ABA distribution within the leaf, we created a mathematical model of a leaf cross-section. The model comprises an area of spongy mesophyll, bounded on the underside by a layer of epidermal cells. Within the spongy mesophyll, we specify a vasculature region and an encasing bundle sheath (**Fig. 3A**). Given the cellular topology within the spongy mesophyll, we developed a model that captures this porous tissue, enabling us to analyze the geometrical features that impact ABA distribution. Thus, we consider a grid of cells, in which cell geometries allow for air gaps (**Fig. 3A**). Within the cells, we simulate ABA biosynthesis, specifying ABA2-mediated AB-ald synthesis in the vasculature region and AAO-mediated conversion of AB-ald in the vasculature (AAO3), bundle sheath (AAO2) and pavement cell layer (AAO1 and AAO2) using the localization data (**Fig. 2I**). To determine whether to also include ABA degradation within the modelled tissues, we performed spatial transcriptomics on *Arabidopsis* leaves and roots, examining ABA degradation processes, focusing on *UGT* and *CYP707A* genes. ABA degradation is regulated by two main pathways: conjugation and hydroxylation.

**Fig. 3.**
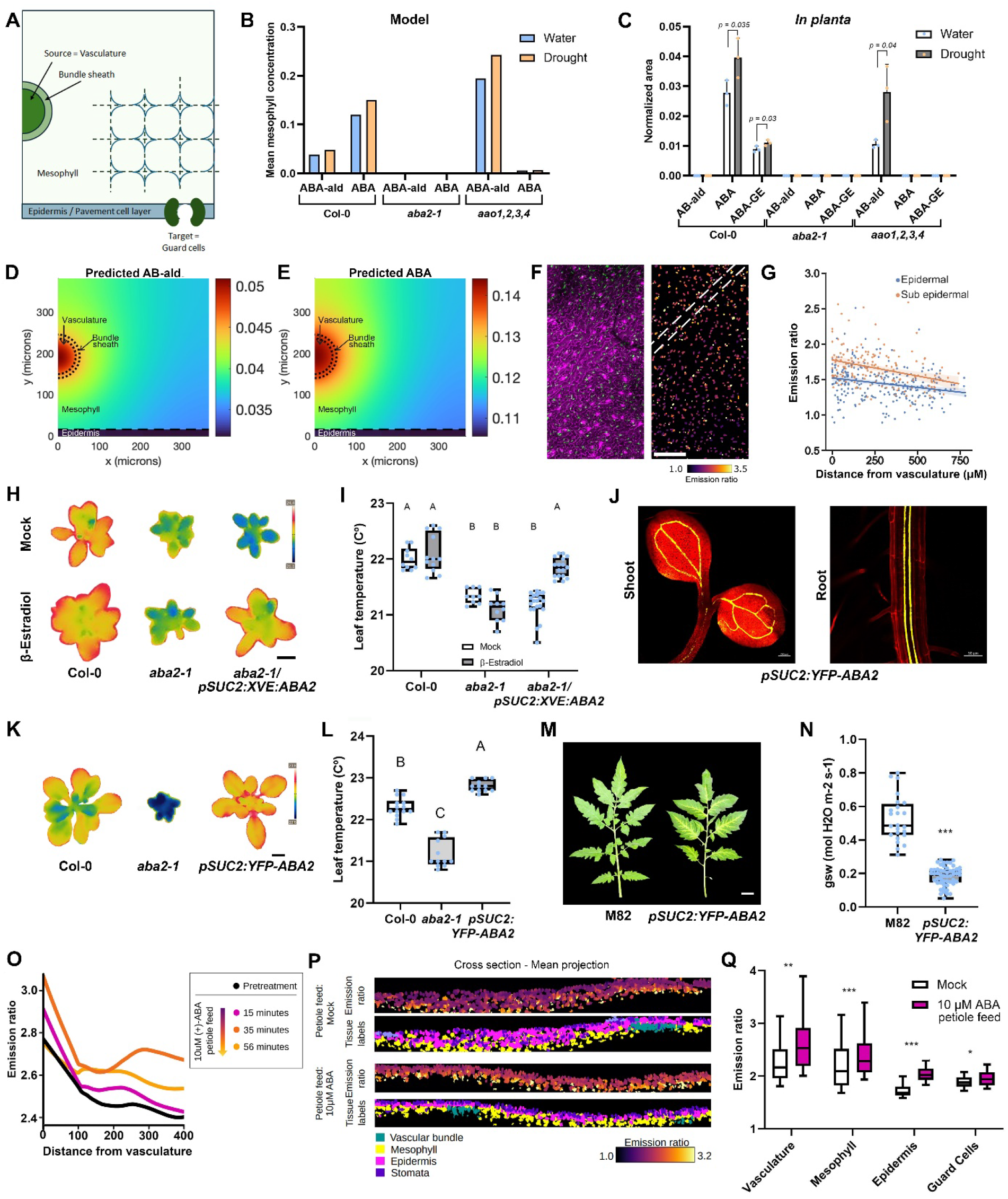
ABA and AB-ald form source to sink gradients in the leaf and move from the phloem to the guard cells to balance stomatal closure. **(A)** Schematic of the mathematical model of the leaf cross-section. **(B)** Model predictions of the mean mesophyll ABA concentrations across different genotypes under well-watered (blue) and drought (orange) conditions. (C) *In planta* AB-ald, ABA and ABA-GE measurements in Col-0, *aba2-1* and *aao1,2,3,4*. Significance was determined using student’s t-test. n ≥ 4. **(D)** A model prediction of AB-ald concentrations in the leaf-cross-section under watered conditions. **(E)** A model prediction of ABA concentrations in the leaf-cross-section under watered conditions. **(F)** ABA sensor (nlsABACUS2–400n - green) and propidium iodide (magenta) emission (left image) and nlsABACUS2–400n emission ratio (right image) in the leaf vasculature and surrounding cells. White dotted line denotes the position of a secondary vein, scale bar = 200 µM. In the emission ratio image, nuclei have been dilated after quantification to make the pattern easier to see. **(G)** Quantifications of ABA sensor emission ratio (nlsABACUS2–400n from F) plotted against distance from the vasculature for leaf epidermal cells and sub-epidermal cells, segmented using EZ-Peeler. n ≥ 148. **(H, I)** Representative thermal images **(H)** and quantification **(I)** of 21-day-old *aba2-1 pSUC2:XVE:ABA2* plants and respective controls, with and without estradiol treatment. 5 μM β-Estradiol was sprayed twice a week for 2 weeks. Shown are averages (±SD), n ≥ 8, different letters represent significant differences. Significance was determined using Tukey’s ad-hoc statistical test (treatments marked with different letters are significantly different). Scale bar = 1 cm. **(J)** Shoot and root images of 7-day-old *Arabidopsis* seedlings expressing *pSUC2:YFP-ABA2*. Scale bar = 20 μm. **(K, L)** Representative thermal images **(K)** and quantification **(L)** of 25-day-old *pSUC2:YFP-ABA2 Arabidopsis* plants and respective controls. Shown are averages (±SD), n ≥ 12. Different letters represent significant differences. Significance was determined using Tukey’s ad-hoc statistical test (treatments marked with different letters are significantly different). Scale bar = 1 cm **(M, N)**. Representative images **(M)** and quantification **(N)** of leaf transpiration of 8-week-old tomato plants expressing *pSUC2:YFP-ABA2*. Scale bar = 1 cm. Significance was determined using Student’s t-test. *** =*p value < 0.005*. **(O)** Emission ratio of nlsABACUS2–400n across distances from the vasculature before treatment (black), 15 (pink), 35 (orange) and 56 (yellow) minutes after 10 µM ABA petiole feed. Leaves were imaged in air, with illumination between imaging acquisitions, without a coverslip to allow transpiration. **(P)** Emission ratio of nlsABACUS2–400n in different leaf tissues after a 35-minute petiole feed of 10 µM ABA or mock solution, illuminated in air. Leaves were mounted in perfluorodecalin after treatment to allow deep imaging. **(Q)** Quantification of Q and R. Significance was determined using Student’s t-test. *** =*p value < 0.005*, ** =*p value < 0.00*1, ** =*p value < 0.0*5.

UGTs reversibly inactivate ABA by conjugating it to glucose (ABA-GE) ^41,42^, while CYP707A cytochrome P450 enzymes irreversibly degrade ABA through hydroxylation to inactive catabolites ^43,44^. *CYP707As* expression was predominantly detected in the vasculature, alongside *UGT71B6, UGT71B8* and *UGT71C5*, while *UGTs* were also expressed in the mesophyll. *UGT71C5* was also expressed in the leaf epidermis (**Sup. Fig. S21, S22**). Notably, we did not observe a change in their gene expression under drought conditions (**Sup. Fig. S21, S23**). Based on these observations, we modelled a constant ABA degradation in all cells.

The model incorporated AB-ald and ABA move between adjacent cells (representing a combination of plasmodesmatal diffusion and membrane transport). Such transport processes depend on the length of the contact regions. Within the spongy mesophyll, the presence of air gaps means only a proportion of each cell length is in contact with adjacent cells. Using idealized cell geometries (**Fig. 3A**), we prescribe contact-region lengths to achieve a specified tissue porosity. In contrast, for the epidermal layer, the wavy pavement-cell boundaries create a contact region longer than the cell width, thereby increasing hormone diffusion within the epidermal layer. We derived a continuum approximation of this cell-based model, which reveals how the effective tissue-scale hormone diffusion rates depend on the cell-scale parameters; (full mathematical details are provided in SI Appendix Text**, Sup. Fig. S24**). Parameterizing the model using geometrical measurements from confocal images (**SI Appendix**), the model predicts the distribution of AB-ald and ABA within the modelled tissues. To validate the model, we sought to test how varying the localized activities of the ABA2 and AAO enzymes affects the distributions of AB-ald and ABA. As expected, the model predicts that reducing ABA2 activity (to 1% to represent *aba2-1*) reduces AB-ald and ABA concentrations throughout, whereas reducing AAO activity (to 1% to represent *aao1,2,3,4*) is predicted to increase AB-ald concentrations and decrease ABA concentrations (**Fig. 3B**), albeit with predicted values depending on the rate constants (**Sup. Fig. S25**). Consistent with these predictions, experimental measurements (LC-MS/MS analysis applied to quantify differential abundances) in *aba2-1* revealed low AB-ald and ABA, whereas in *aao1,2,3,4* the measurements validated higher AB-ald and lower ABA (**Fig. 3C**). The LC-MS/MS measurements also revealed an increase in both AB-ald and ABA under water stress conditions - assuming a 25% increase in AB-ald production, the model predictions are also consistent with these water-stress hormone profile measurements in WT, *aba2-1* and *aao1,2,3,4* (**Fig. 3B-C**).

Model simulations predict a gradient of AB-ald across the mesophyll, reducing away from the ABA2-mediated source in the vasculature region (**Fig. 3D**). AAO-mediated conversion of AB-ald into ABA is predicted to result in high ABA concentrations in the vasculature region and a gradient through the mesophyll (**Fig. 3E**). Notably, the ABA level within the epidermal layer depends on the rate of diffusion between the mesophyll and epidermis, and the rates of AAO conversion and ABA degradation. By choosing low rates of mesophyll-to-epidermis diffusion and AAO conversion and a high rate for ABA degradation, the model predicted low epidermal ABA levels in agreement with ABACUS observations ^45^ To validate the predicted ABA leaf distribution, we exploited the ABACUS2-400n FRET sensor ^45^ to visualize bioactive ABA levels at cell-level resolution. *In planta* ABACUS2-400n biosensor data were consistent with our model predictions, showing ABA gradients, with hormone levels reducing with distance from the major vein, in addition to higher emission ratios of sub-epidermal cells compared to epidermal cells (**Fig. 3F-G**).

### ABA and/or AB-ald move from the phloem to the guard cells to regulate stomatal closure

To test whether ABA synthesized in the vasculature can dynamically travel from leaf vascular tissues to regulate guard cells, we engineered an estradiol-inducible system to control ABA biosynthesis in phloem companion cells (*aba2-1 pSUC2:XVE:ABA2*). We observed that estradiol treatment restored stomatal function in *aba2-1* compared to mock-treated plants (**Fig. 3H-I**). The shoot growth of the plants also showed a similar trend: the leaf surface area in the E2-treated *aba2-1 pSUC2:XVE:ABA2* plants was similar to that of wild-type, whereas non-induced *aba2-1 pSUC:XVE:ABA2* plants exhibited a smaller surface area like *aba2-1* (**Sup. Fig. S26A-B**). To confirm the cell type-specific activity of ABA2 in the phloem, we cloned *pSUC2:YFP-ABA2* and generated transgenic *Arabidopsis* plants. Confocal images revealed that the YFP-ABA2 translational fusion is localized in the phloem (**Fig. 3J**), ruling out possible ABA2 protein movement. Expression of *YFP-ABA2* in the phloem increased leaf temperature compared to controls (**Fig. 3K-L**). Overall, our functional studies confirm that either AB-ald and/or ABA or downstream effectors of ABA can move from the vasculature to guard cells, indicating the effectiveness of leaf ABA and AB-ald movement.

Next, we investigated whether ABA or AB-ald movement from the vasculature to the guard cells is conserved in other plant species. To test this, we generated tomato *pSUC2:YFP-ABA2* transgenic plants and observed that enhancing ABA biosynthesis in phloem companion cells promoted stomatal closure in tomato, as indicated by stomatal conductance measurements (**Fig. 3M-N**, **Sup. Fig. S26C**).

To determine whether ABA (rather than a downstream signal) is able to move from leaf vascular tissues to guard cells in response to environmental stress, we monitored the ABACUS2-400n sensor after elevating ABA levels in the leaf vasculature (via petiole feeding - injecting 10 µM ABA into the leaf petiole). Strikingly, feeding studies led to a rapid increase in ABA levels spreading throughout the mesophyll within 35 minutes (**Fig. 3O, Sup. Fig. S27**). Importantly, these data demonstrate the direct accumulation of vasculature-derived bioactive ABA in guard cells (**Fig. 3P, Q**). Taken together, our functional and reporter results reveal that ABA can move from phloem companion cells to guard cells.

### Tissue-restricted ABA biosynthesis knockouts demonstrate that mobile ABA and AB-ald govern stomatal aperture and xerobranching

Our results in leaves reveal that ABA gradients are formed and that ABA can move from phloem to guard cells to regulate water-stress responses (**Fig. 3**). To evaluate whether ABA is the main mobile signal or whether AB-ald also moves, we performed knockouts in *ABA2* and *AAO3* ABA biosynthesis genes in a tissue-specific manner using CRISPR/Cas9 ^46^, and quantified stomatal conductance and xerobranching. To test the efficiency of the sgRNAs, we first overexpressed Cas9 using the *RPS5* promoter ^47^ to generate stable heritable mutations and compared them with known knockout lines. As expected, *sg-ABA2* and *sg-AAO3* expressing plants exhibited *aba2-1* and *aao3-4* leaf phenotypes, respectively (12/12 lines for *ABA2*, 14/15 lines for *AAO3*) (**Fig. 4A-B**). Since the efficiency was high, we cloned several different tissue-specific promoters upstream to Cas9: *Sucrose-Proton Symporter 2* (*SUC2*) for phloem companion cells expression ^48^; *Coronatine Induced 3 (CORI3)* for spongy mesophyll expression ^49^; *Voltage-dependent K+ channel (KST1)* for guard-cell specific expression ^50^; *Plastid Transcription Factor AtSIG6* (*SIG6*) for shoot-specific expression ^51^; *Arabidopsis Root Specific Kinase 1* (*ARSK1*) for root-specific expression ^52,53^; SCARECROW (*SCR*) for endodermis and bundle sheath expression ^54^; and meristem layer 1 (*ML1*) for shoot epidermis expression ^55^ (**Fig. 4C**).

**Fig. 4.**
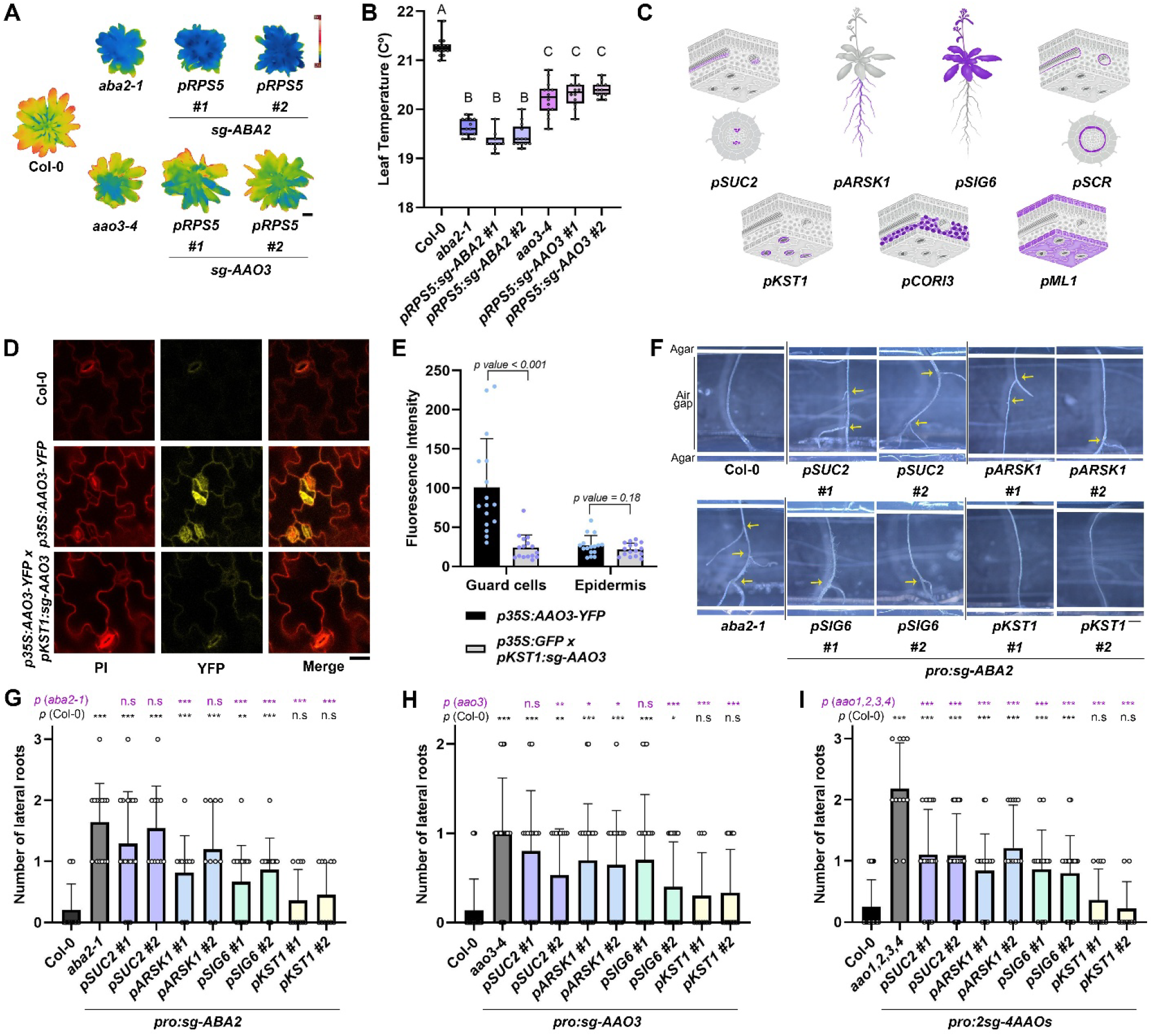
Tissue-specific ABA synthesis knockout suggests that long-distance ABA and AB-ald regulate stomatal aperture and xerobranching. **(A, B)** Thermal images **(A)** and quantification **(B)** of 45-day-old *pRPS5:sgRNA-ABA2/AAO3* plants with respective controls. Plants were grown under short-day conditions. Shown are averages (±SD), n ≥ 10; Significance was determined using Tukey’s ad-hoc statistical test (treatments marked with different letters are significantly different). Scale bar = 1 cm. **(C)** Illustration of tissue-specific promoters cloned for targeting *ABA2 AAO3* and *4AAOs*. **(D, E)** Images **(D)** and quantification **(E)** of 7-day-old *p35S:AAO3-YFP x pKST1:sg-GFP* expression and respective controls in the guard cells and pavement cells. Scale bar = 50 uM. n ≥ 16; Significance was determined using Student’s t-test. **(F-I)** Shown are images **(F)** and quantification **(G-I)** of 13-day-old roots of cell type-CRISPR lines targeting *ABA2* (G), *AAO3* (H), and *4AAOs* (I) with respective controls in an agar-based air-gap assay system (Air gap = ∼5 mm). Mutant alleles: *aao1-2, aao2-1, aao3-4, aao4-2*. Yellow arrows indicate lateral roots. Scale bar = 1 mm. n ≥ 9; Significance was determined using Student’s t-test. * = *p value < 0.05,* ** = *0.005 < p value < 0.01,* and *** = *p value < 0.005*.

The promoters selected span a range of leaf and root tissues, from the vasculature to the epidermis, enabling spatial resolution of ABA biosynthesis activity. To validate Cas9 activity in a defined tissue, we generated a transgenic *pKST1:sg-AAO3* line and crossed it with *p35S:AAO3-YFP*. We found a significant reduction in fluorescent intensity taking place only in the guard cells, and not in the pavement cells surrounding them, indicating a robust and tissue-specific editing strategy (**Fig. 4D-E, Sup. Fig. S28**). Next, *sgRNA-ABA2, sgRNA-AAO3*, and *2sg-4AAOs* (**Sup. Table 1**) were cloned downstream of Cas9 and transformed into the background of wild-type plants. *pML1* and *pSCR* were ruled out as they produced stable inherited mutations, likely due to embryonic or meristematic expression (**Sup. Fig. S29**).

We initially assessed the location of ABA biosynthesis required for the root xerobranching response. After performing the air gap-based bioassay on these tissue-specific lines, we observed *pSUC2:sg-ABA2* (phloem companion cells) lines showed an increased number of lateral roots compared to the WT, closely resembling the *aba2-1* mutant phenotype (**Fig. 4F-G**), confirming ABA2 activity in the phloem companion cells is required for xerobranching. Additionally, *pARSK1* (root-specific) and *pSIG6* (shoot-specific) lines targeting *ABA2* displayed an intermediate phenotype, indicating root and shoot ABA2 activity contribute to xerobranching (**Fig. 4F-G**). This is the first indication that shoot-derived ABA and/or AB-ald is needed for the root xerobranching response. On the other hand, *pKST1:sg-ABA2* (guard cell-specific) lines did not show a significant reduction compared to Col-0 (**Fig. 4F-G**). Similar trends were observed for both the *AAO3* and quadruple *aao* (*4AAO*) mutants (**Fig. 4H, I, Sup. Fig. S30**). Targeting *AAO* genes in the phloem companion cells resulted in a partial xerobranching phenotype, with increased lateral root formation, which did not fully reproduce the strong response observed in the corresponding mutant controls. This suggests that either the Cas9-mediated cell type-specific DNA editing efficiency is not high enough, or that AAO activity in the phloem contributes to the xerobranching response but is not sufficient on its own. Likewise, root and shoot-specific knockouts also displayed partial phenotypes, indicating that AAO functions in both organs are required for regulating xerobranching (**Fig. 4H-I**).

To assess the contribution of local ABA biosynthesis during stomatal closure, we analyzed leaf temperatures under well-watered and drought conditions across CRISPR lines targeting *ABA2*, *AAO3*, and *4AAOs* (**Sup. Fig. S31A-F**). In all three lines, expressing the sgRNAs under the phloem-specific (*pSUC2*), root-specific (*pARSK1*), or shoot-specific (*pSIG6*) promoters showed lower leaf temperatures compared to Col-0, suggesting reduced stomatal closure activity. However, these lines displayed only partial phenotypes relative to their corresponding full mutants, indicating that the targeted genes also act in additional tissues, or, in the case of *AAO3* that other members of the AAO family mediate partial ABA synthesis. The expression of *ABA2* and *AAO3* (**Sup. Fig. S3**) detected in the leaf phloem companion cells but also the surrounding cells (cambium and phloem) (**Fig. 1**), could explain why phloem companion cell-specific knockout of *ABA2* or *AAO3* using the *SUC2* promoter resulted in only a partial phenotype (**Sup. Fig. S31A-D**). Following 10 days of drought stress, the *pSUC2:sg-ABA2* (phloem companion cell) lines showed a significant compromise in leaf temperature compared to WT, indicating that the phloem contributes specifically to stomatal regulation under drought. None of the other promoters expressing *sg-ABA2* showed consistently significant results across all independent lines (**Sup. Fig. S31A-F**). Notably, knocking out the different *ABA* biosynthesis genes specifically in the guard cells or spongy mesophyll did not result in a reduction in leaf temperature, indicating that ABA synthesis in these cell types does regulate stomatal conductance, in line with the lack of *ABA2* or *AAO* expression in these tissues (**Sup. Fig. S31G-L**). More importantly, *pSUC2:sg-AAO3* and *pSUC2:2sgRNA-4AAO* were also consistently significant, underscoring their requirement in the phloem companion cells and for ABA movement from the phloem to the guard cells during drought response. Collectively, the findings suggest a significant role of phloem-localized ABA biosynthesis as a contributor to both xerobranching and stomatal closure processes.

### AAOs distribution is functionally important, leading to a dual-transport pathway of ABA to guard cells

ABA2 and AAO3 co-expression in the phloem suggests companion cells are a key cell type for ABA synthesis; however, the roles of AAOs in other root and shoot tissues remain unclear. We propose two pathways. (1) Bioactive ABA is synthesized directly within the phloem by co-expression of *ABA2* and *AAO3* and then transported to guard cells (**Fig. 5A**). (2) ABA2 in the phloem produces AB-ald, which moves to outer tissues where AAOs convert it to bioactive ABA (such as the epidermis pavement cells and bundle sheath in the shoot, or the cortex and endodermis in the root); the resulting ABA then moves to the target sites (**Fig. 5B**).

**Fig. 5.**
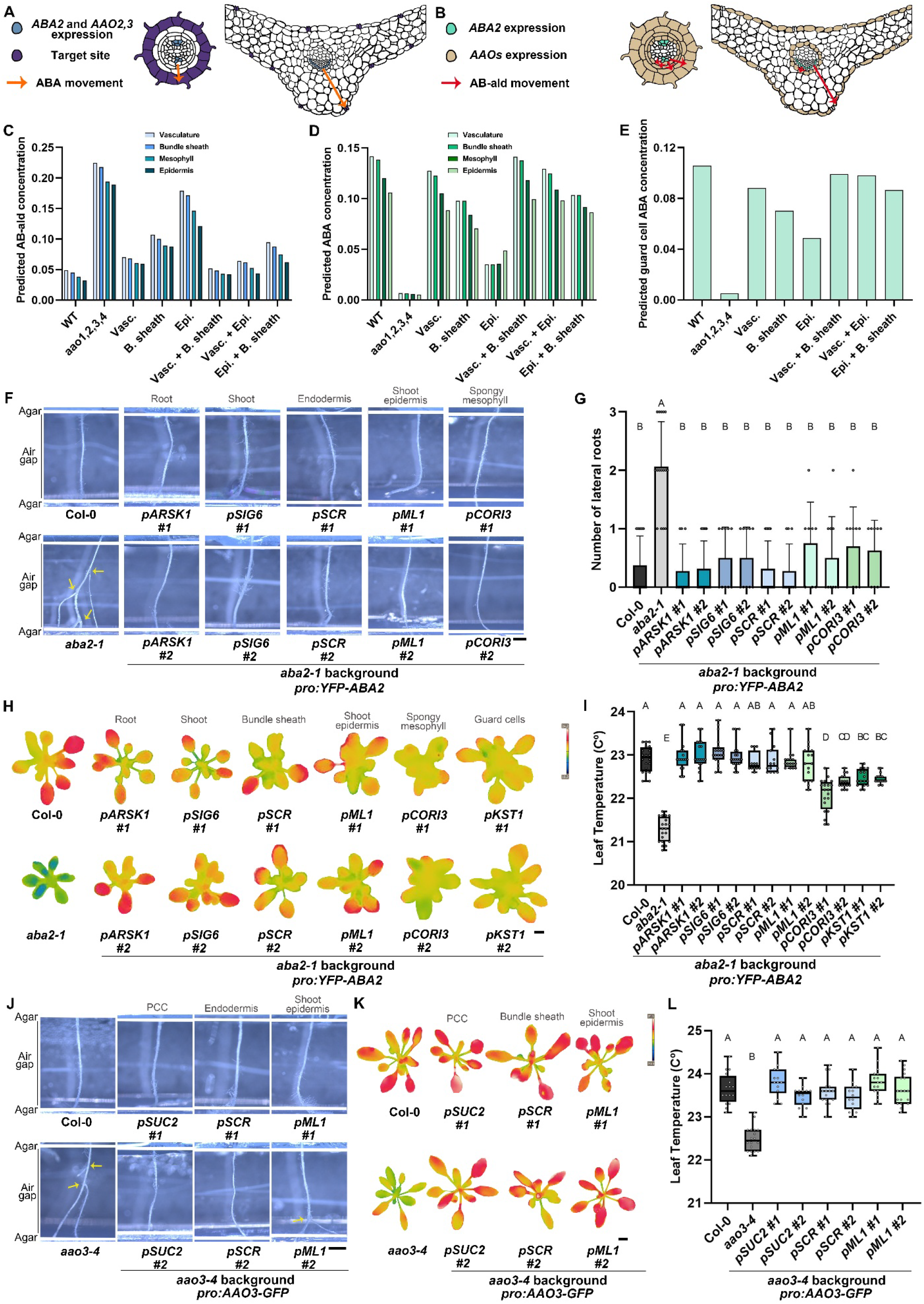
Both ABA and AB-ald movement are required for xerobranching and stomatal closure responses. **(A)** Model of ABA movement from the phloem (where *ABA2*, *AAO3* and *AAOs* can be expressed, marked in blue) to the target site (epidermis in the root, guard cells in the shoot, marked in purple). Arrow represents ABA movement in orange. **(B)** Model of abscisic aldehyde movement from the phloem (where ABA2 is expressed, marked in green) to the sites of AAOs expression (pericycle, endodermis, cortex and epidermis in the root, bundle sheath and pavement cells in the shoot, marked in light brown). Arrow that represents abscisic aldehyde movement in red. **(C-E)** Predicted AB-ald **(C)**, ABA **(D)** and guard cell-ABA **(E)** concentrations when AAOs are localized in specific tissues. Vas. = Vasculature, B.sheath = Bundle sheath, Ep. = Epidermis. **(F, G)** Representative images **(F)** and quantification **(G)** *pARKS1, pSIG6, pML1, pCORI3,* and *pSCR:YFP-ABA2* on the background of *aba2-1* with respective controls in an agar-based air-gap assay system (Air gap = ∼ 5 mm). Shown are averages (±SD), n ≥ 8, and different letters represent significant differences. Significance was determined using Tukey’s ad-hoc statistical test (treatments marked with different letters are significantly different). Yellow arrows indicate lateral roots. Scale bar = 1 mm. **(H, I)** Representative thermal images **(H)** and quantification **(I)** of 26-day-old *pARKS1, pSIG6, pML1, pCORI3, pKST1,* and *pSCR:YFP-ABA2* on the background of *aba2-1* with respective controls. Shown are averages (±SD), n ≥ 12, and different letters represent significant differences. Significance was determined using Tukey’s ad-hoc statistical test (treatments marked with different letters are significantly different). Scale bar = 1 mm. **(J)** Representative images of 13-day-old *pSUC2, pSCR, and pML1:AAO3-GFP* on the background of *aao3-4* roots with respective controls in an agar-based air-gap assay system (Air gap = ∼5 mm). Yellow arrows indicate lateral roots. Scale bar = 1 mm. **(K, L)** Representative thermal images **(K)** and quantification **(L)** of 26-day-old *pSUC2, pSCR, and pML1:AAO3-GFP* on the background of *aao3-4* with respective controls. Shown are averages (±SD), n ≥ 14, and different letters represent significant differences. Significance was determined using Tukey’s ad-hoc statistical test (treatments marked with different letters are significantly different). Scale bar = 1 mm.

To test these different possibilities and understand how the localization of ABA2 and AAOs contributes to ABA distribution, we used the mathematical model to predict how tissue-specific AAO localizations affect the distributions of AB-ald and ABA across the leaf tissues and the ABA level at a guard cell (i.e., within the pavement cells, taken to be at a horizontal distance of 330 μm from the vein). We first removed AAOs either from the vasculature, bundle sheath, or epidermal cells (**Fig. 5C-E**). The model predicts that the presence of AAOs in the vasculature, bundle sheath and pavement cells all contribute to the ABA level at the guard cell. Removing AAOs from any of these three tissues is predicted to reduce guard cell ABA levels (**Fig. 5E**), albeit with relative contributions from each tissue depending on the unknown rate constants governing AAO-mediated conversion in different tissues (**Sup. Fig. S25**). These model predictions are consistent with the observation that phloem-specific *AAO3* and *4AAO* mutants have a partial leaf-temperature phenotype (**Sup. Fig. S31A-F**); the predictions suggest that with no AAO-mediated AB-ald conversion in the vasculature, more AB-ald diffuses into the bundle sheath and epidermis, where the remaining AAOs convert it to ABA, leading to a predicted guard cell ABA concentration much higher than that of the quadruple *aao* mutant (albeit less than that predicted for wild-type) (**Fig. 5E**). Thus, the modelling suggests that in wild-type, the guard cell ABA level is the combination of both ABA synthesized in the vasculature (as in **Fig. 5A**) and ABA synthesized in the epidermis from the AB-ald that has diffused/transport from the vasculature (as in **Fig. 5B**).

Next, we set up an experimental system to determine whether bioactive ABA or AB-ald can move between distal tissues to regulate stomatal closure and xerobranching. We hypothesized that if AB-ald and/or bioactive ABA could move from phloem companion cells to guard cells, expressing an ABA2 or AAO3 enzyme in distinct cells would rescue the respective *aba2-1* or *aao3-4* mutant phenotype. To test this hypothesis and also elucidate the xanothoxin distribution, we cloned *ABA2* downstream of several additional tissue-specific promoters: *pKST1* ^50^*, pSCR* ^54^*, pML1* ^55^*, pCORI3* ^49^*, pARSK1* ^52^ and *pSIG6* ^51^ (**Fig. 4C**) and transformed the constructs into the *aba2-1* mutant background. Comparison with their respective controls revealed that expressing *ABA2* under all tested tissue-specific promoters restored the *aba2-1* root phenotype in the agar air gap (**Fig. 5F-G**). In the shoot, expression of *ABA2* under root-specific (*pARSK1*), shoot-specific (*pSIG6*), endodermis and bundle sheath-specific (*pSCR*), and shoot epidermis-specific (*pML1*) promoters fully rescued the *aba2-1* mutant phenotype (**Fig. 5H-I**). Expression under the spongy mesophyll-specific (*pCORI3*) and guard cell-specific (*pKST1*) promoters also led to rescue, although not to the same extent as wild-type or the other cell type-specific constructs (**Fig. 5H-I**). These findings support the conclusion that the ABA precursor, xanthoxin, is present across multiple tissues and that AB-ald and/or bioactive ABA can move from these cells to their target sites, thereby promoting ABA-mediated responses.

To further explore the mobility of bioactive ABA, we generated lines in which ABA is synthesized only in specific tissues. First, using the mathematical model, we predicted that having AAO in only one tissue reduces the predicted guard cell concentration (**Fig. 5E)**. Interestingly, the model predicted that the predicted guard cell ABA concentration in wild-type is much lower than the sum of the three predicted values with AAOs in the individual tissues due to the availability of AB-ald. For example, with AAO only in the epidermis, the lack of AAO-mediated AB-ald depletion in the vasculature and bundle sheath enables more AB-ald to move to the epidermis, providing more precursor for the epidermal AAO to convert to ABA (**Fig. 5E**).

To test these model predictions, we generated plants expressing *AAO3* in phloem companion cells (*pSUC2*), endodermis and bundle sheath (*pSCR*), and shoot epidermis (*pML1*) in the *aao3-4* mutant background. Expressing *AAO3* in all these tissues rescued the *aao3-4* deficient mutant phenotype in roots and shoots (**Fig. 5J-L, Sup. Fig. S32)**. Since AB-ald is produced largely in the phloem, these results suggest that AB-ald moves out of the phloem and is present in physiological levels across various tissues, including the endodermis, bundle sheath and epidermis. In addition, the data suggest that bioactive ABA can move from any of these sites to regulate xerobranching and stomatal aperture. Similarly, the data indicate that AB-ald is present in the leaf epidermis at physiological levels, and can be converted into bioactive ABA by *AAO1* and *AAO2*, which can then move to the guard cells to regulate their conductance. This result is in agreement with the *aao1,aao2* double mutant, which are both predominantly expressed outside the leaf vasculature and exhibit a significant thermal leaf temperature phenotype under stress conditions, indicating AB-ald movement out of the phloem (**Fig. 1G-H**). Collectively, the model prediction and *AAOs* misexpression data results provide further evidence that AB-ald can move out of the phloem and that *AAO*s can also work from cell types other than the phloem to execute local stress responses.

We conclude that the ABA accumulation in the guard cell ABA is due to a combination of both hypothesized mechanisms: ABA is synthesized within the vasculature via ABA2 and AAO3 and diffuses/transported to the guard cell, and in parallel, AB-ald synthesized within phloem diffuses/transport across the mesophyll and is converted to ABA in the epidermis via the AAO1 and AAO2 (**Fig. 5A-B**).

### Exploring the mechanisms of ABA movement across membranes

In leaves, stomatal closure progressively reduces water movement and thereby limits bulk hydraulic flow. Yet our physiological experiments and ABACUS2 sensor data (**Fig. 3**) suggest that after phloem ABA induction, the signal accumulates in the epidermis and guard cells within 20–30 minutes and is sustained there during stomatal closure. The persistence of ABA accumulation despite progressively restricted water flow suggests that mechanisms beyond bulk hydraulic transport contribute to ABA distribution within leaves.

Our results reveal that ABA biosynthesis initiates predominantly in phloem companion cells and that both ABA and its precursor, AB-ald, move from their sites of synthesis to distal target tissues to mediate root (xerobranching) and leaf (stomatal closure) water stress responses. While movement of AB-ald and ABA from the vasculature plays a central role in coordinating root and shoot water stress responses, the exact route and dynamics are unclear. To assess the role of ABA and AB-ald movement, we used our mathematical model to predict how the rate of ABA and AB-ald diffusion between adjacent cells affects spatiotemporal ABA redistribution in response to water stress. Based on our measurements showing increased levels of AB-ald and ABA under water stress (**Fig. 3C**), we predict an increase in AB-ald and ABA levels in the vasculature; these higher levels spread across the leaf, while maintaining the spatial gradients away from the vasculature source (**Sup. Fig. S33**). Simulations predict that a cell-to-cell permeability of 0.5 µm/s for both ABA and AB-ald would be required to enable a substantial increase in guard cell ABA levels within 30 minutes (**Fig. 6A**).

**Fig. 6.**
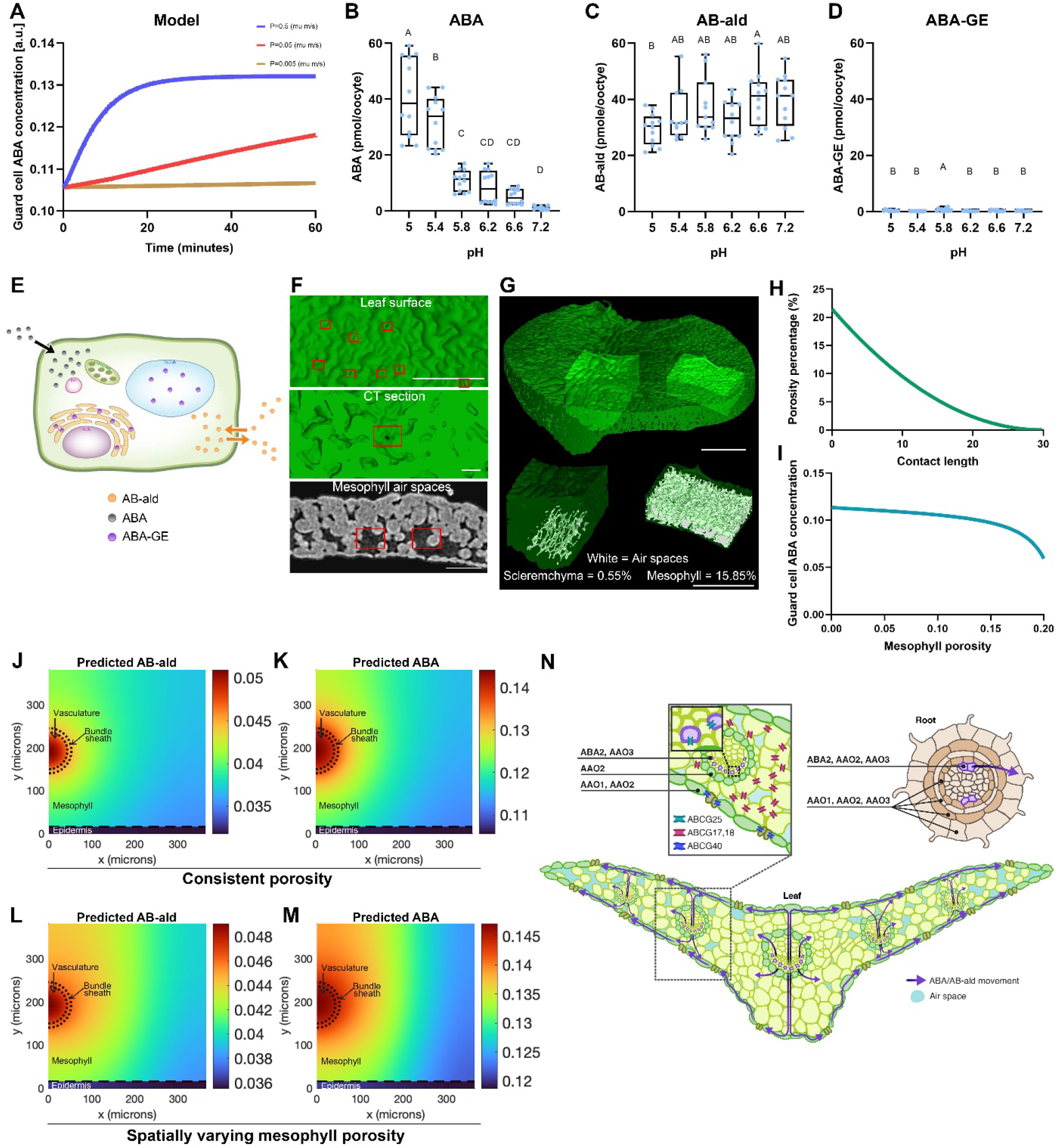
Membrane permeabilities and mesophyll porosity affect ABA and AB-ald routes and concentrations towards the guard cells. **(A)** Predicted guard cell ABA concentration after a stress-induced increase in AB-ald production in the vasculature. The predictions show how the cell-to-cell permeability, P (mu m/s), affects the guard cell ABA dynamics. **(B-D)** Diffusion rates of ABA **(B)**, AB-ald **(C)** and ABA-GE **(D)** in oocytes at different external pH. Shown are averages (±SD), n ≥ 6, and different letters represent significant differences. Significance was determined using Tukey’s ad-hoc statistical test (treatments marked with different letters are significantly different). **(E)** Illustration showing membrane permeability of ABA (grey), ABA-GE (purple), and abscisic aldehyde (orange). A one-way black arrow indicates ABA entry into the cell, while a two-way orange arrow represents bidirectional diffusion of abscisic aldehyde across the plasma membrane. ABA-GE is captured within the ER and the vacuole. **(F)** *Arabidopsis* 3D render leaf disk CT image of abaxial leaf surface, main leaf body. Scale bar = 0.1 mm; *Arabidopsis* 3D render leaf disk CT image of adaxial leaf surface, main leaf body, stomata highlighted, scale bar = 0.1 mm; 2D CT image of region used for mesophyll porosity measurement. Stomata position aligned to 3D image. Scale bar = 0.1 mm. Guard cells and stomata are marked in red rectangles. **(G)** *Arabidopsis* 3D render leaf disk CT image and location of regions of porosity analysis. Scale bar = 0.4 mm. Regions of interest in *Arabidopsis* leaf CT image for porosity analysis (left: mid vein/sclerenchyma; right: Leaf body/ mesophyll). Scale bar = 0.4 mm. Regions of interest in *Arabidopsis* leaf CT image showing total intercellular air space in white. Porosity is expressed as the % of intercellular air to the total volume of the region of interest. Left: Mid vein = 6.68%. Right: Leaf body = 18.09%. Scale bar = 0.4 mm. Regions of interest in *Arabidopsis* leaf CT image showing total intercellular air space in white. Porosity is expressed as the % of intercellular air to the total volume of the region of interest. Left. Sclerenchyma = 0.55%, Right. Mesophyll = 15.85%. Scale bar = 0.4 mm. **(H)** Model predicting contact region length depends on porosity. **(I)** Model predicting reduced porosity (which corresponds to larger contact lengths) results in higher guard cell concentrations (at equilibrium). **(J)** Predicted AB-ald concentrations in the shoot under watered conditions and consistent porosity. **(K)** Predicted ABA concentrations in the shoot under watered conditions and consistent porosity. **(L)** Predicted AB-ald concentrations in the leaf-cross-section under watered conditions, with a spatially varying mesophyll porosity. **(M)** Predicted ABA concentrations in the leaf-cross-section under watered conditions with a spatially varying mesophyll porosity. **(N)** Illustration of ABA/AB-ald movement in the leaf from the phloem to the guard cells target sites, moving through reduced porosity tissue.

We first questioned whether the required AB-ald and ABA movement could be achieved via passive diffusion across the membrane. We quantified the membrane permeability of ABA, AB-ald and ABA-GE in *Xenopus* oocytes across a range of external pH conditions, simulating the pH gradient between the apoplast and cytosol. ABA accumulation showed a strong pH dependence, with significantly higher levels able to cross the membrane at acidic pH and sharply reduced levels at neutral pH (**Fig. 6B**). This pattern suggests that ABA enters cells efficiently at apoplastic pH but exits poorly at cytosolic pH (i.e., acid trapping). In contrast, AB-ald accumulated to relatively stable levels across all pH conditions tested (**Fig. 6C**), implying it can move more freely across cell membranes and is not influenced by pH-dependent trapping. ABA-glucose ester (ABA-GE) levels remained consistently low across all pH values (**Fig. 6D**), indicating poor membrane permeability and that it is likely confined to internal compartments, such as the endoplasmic reticulum (ER) or vacuole, where the BG enzymes are expressed ^17,43^. These results suggest that ABA membrane permeability is strongly influenced by pH, facilitating cellular uptake at acidic apoplastic conditions but limiting exit at neutral cytosolic pH. In contrast, AB-ald crosses membranes more readily, whereas ABA-GE remains largely membrane-impermeable, likely confined to internal compartments, and overall affects ABA accumulation in guard cells (**Fig. 6A, E**). The results may suggest that under water-stress conditions, when apoplastic pH increases and more ABA is trapped in the apoplast as the deprotonated form, AB-ald can still enter the cell, where it can be converted to ABA by AAOs.

To assess whether oocyte uptake measurements equate to the required permeability identified via the mathematical model (**Fig. 6A**), the oocyte measurements were incorporated into a mathematical model of oocyte uptake ^56^, enabling estimation of the values for the AB-ald and ABA membrane permeabilities (Supp Appendix Text, section 7). The estimated permeabilities were substantially lower than those suggested by the model (**Sup. Fig. S34A-B**). Predictions indicated that using these estimated values led to very slow hormone diffusion from the vasculature, yielding lower concentrations of AB-ald and ABA in the epidermis (**Sup. Fig. S34C-D**). The results indicate that the passive membrane transport observed in oocyte experiments would not be sufficient to cause substantial hormone movement to the guard cells within an appropriate time.

One option that could explain the movement of AB-ald or ABA is specialized transporters that would allow hormone movement across cell membranes. The model predicted that if only ABA moves between adjacent cells, ABA synthesized in the vasculature via AAO3 is sufficient to maintain an ABA gradient across the mesophyll and to reach the guard cells (**Sup. Fig. S34E-F**). In contrast, if only AB-ald moves between adjacent cells, the model predicted that ABA synthesized in the vasculature cannot diffuse across the leaf, resulting in very low ABA levels throughout the mesophyll, in contrast to the ABACUS2 observations (**Sup. Fig. S34G-H**). These modelling results therefore suggest that specialized transporters mediating ABA movement, rather than AB-ald movement alone, could account for the observed ABA distribution pattern. Because several ABA transporters have been identified, whereas no transporters are currently known for AB-ald, we focused subsequent experiments on testing transporter-mediated ABA movement.

Based on model predictions and oocyte assays, we next designed experiments to test potential routes of ABA movement in planta, with a focus on membrane transporters. Previous studies have identified several ABA transporters, including ABCG25, which exports ABA from leaf companion cells ^30^, and ABCG40, which facilitates ABA import into guard cells ^29^. To assess whether these transporters are essential for ABA movement from the phloem to target sites, we performed air-gap assays using *abcg25* and *abcg40* mutants, evaluated lateral root formation, and employed thermal imaging to assess stomatal closure in shoots. Additionally, we crossed these mutants with the *aba2-1/pSUC2:XVE:ABA2* line, which restores ABA2 function specifically in phloem companion cells and rescues the *aba2-1* phenotype (**Fig. 3H, I**) ^6^. In all cases, the transporter mutants displayed phenotypes similar to both wild-type and the inducible *ABA2* line, with no observable defects in lateral root suppression or stomatal closure (**Sup. Fig. S35**). Notably, *Yang et al., 2024* reported that the *abcg25* mutant shows a stomata aperture phenotype under non-drought conditions ^57^, which was not observed here, and is likely dependent on growth conditions and experimental setup. Together, our results suggest that ABCG25 and ABCG40 are not the primary transporters responsible for ABA export from the phloem or import into guard cells. Instead, yet-to-be-identified transporters are required to facilitate ABA movement from its phloem source to the epidermal stomata.

### Mesophyll air spaces dictate the pathway for ABA movement in the leaf

Leaf lamina tissues feature extensive intercellular air spaces that surround the mesophyll, accelerating CO_2_ and O_2_ exchange to support photosynthesis ^58^. Because ABA and AB-ald are water-soluble and do not volatilize at physiological temperatures, we hypothesized that these air spaces could strongly influence the movement of ABA and AB-ald from the phloem to their target, the guard cells. To test this, we imaged mature *Arabidopsis* leaves using computed microtomography (micro-CT). Micro-CT reconstructions reveal that guard cells are physically separated from the mesophyll and in direct contact with only epidermal pavement cells; beneath each guard cell lies a prominent sub-stomatal air cavity (**Fig. 6F**), which is thought to facilitate efficient gas exchange ^58,59^. In this context, transpiration drives a large water flux through the leaf: water is drawn along a water potential gradient into the mesophyll, where it evaporates from mesophyll surfaces into the intercellular air space and exits through the stomatal pore, a process that has been extensively modeled in anatomically explicit frameworks ^60^. We reasoned that these cavities could modulate hormone transport, potentially influencing the movement of ABA and AB-ald between tissues.

To understand how anatomical organization impacts ABA movement across the leaf, we quantified tissue porosity, defined as the percentage of intercellular air volume within a region of interest relative to the total volume. Mesophyll porosity away from the vein was 15.85% (**Fig. 6G**), indicating extensive air space that would limit the continuity of aqueous pathways for freely dissolved metabolites such as ABA. We therefore hypothesized an alternative route: that ABA and AB-ald predominantly move vertically through the sclerenchyma located beneath the vascular bundles to reach the epidermal pavement cells. Consistent with this possibility, the sclerenchyma surrounding the vasculature exhibited only 0.55% porosity, approximately 3% of the mesophyll’s porosity, suggesting a denser, more continuous tissue that could facilitate solute transport (**Fig. 6G**).

To further investigate the impact of mesophyll airspaces, we varied porosity values in our mathematical model. Increasing porosity reduces the length of the contact region between neighboring cells - with the idealized cell geometries (**Fig. 3A**) and measured mesophyll cell size, a porosity over ∼22% would result in the contact regions reducing to zero and no cell-to-cell hormone movement (**Fig. 6H**). Considering a spatially uniform mesophyll porosity, the model predicts that high levels of porosity substantially reduce hormone diffusion across the mesophyll and the predicted guard cell ABA concentration (**Fig. 6I**). Introducing a spatially varying porosity using the micro-CT measurements, the model predicts that the mesophyll airspaces have a major impact on the path taken by ABA, with the hormone signal diffusing predominantly towards the epidermis, leading to a higher predicted ABA guard cell concentration (**Fig. 6J-M**). Thus, the modelling suggests that the tightly packed cells beneath the vein provide an efficient pathway for hormone transport, facilitating rapid ABA delivery to the guard cells to mediate stress responses (**Fig. 6N**).

## Discussion

ABA biosynthesis is critical for root and shoot water-stress responses ^9^. Despite ABA’s long-known importance for plant abiotic stress responses, major questions remained unanswered until this study. For example, which specific cell types produce ABA? Does the localization or rate of ABA cellular-level production change in response to stress over time and space? Is ABA movement necessary for its functional activity during stress responses? If so, what mechanisms enable its transport, which chemical forms of ABA are mobile, and are these mechanisms conserved between the root and the shoot?

Our study provides compelling evidence that ABA and its precursor, AB-ald, move within leaf and root tissues to coordinate water stress adaptive responses. The phloem acts as a biosynthetic hub for ABA production, with ABA synthesized predominantly in vascular tissues under water stress and subsequently transported to guard cells. Additionally, we provide evidence supporting the importance of the movement of the ABA precursor, AB-ald. Targeted knockout of *ABA2* specifically in the phloem companion cells resulted in a xerobranching phenotype similar to that of the *aba2-1* mutant (**Fig. 4**), highlighting the functional importance of this tissue in root water-stress responses. However, phloem-specific knockout of *AAO3* or all *AAOs* did not fully reproduce their respective mutant phenotypes, suggesting that a proportion of AB-ald moves out of the phloem and is converted by the AAOs to bioactive ABA in the ground tissues.

Tissue-specific complementation experiments further demonstrated that restoring *ABA2* or *AAO3* expression exclusively in phloem companion cells was sufficient to rescue both stomatal closure and lateral root suppression in ABA-deficient backgrounds (**Fig. 3, 5**), supporting a model in which ABA and/or AB-ald move from the phloem to outer target tissues. Intriguingly, expressing *ABA2* in guard cells or spongy mesophyll cells partially rescued the *aba2-1* stomatal aperture phenotype, but not to the level of wild-type plants (**Fig. 5**). This partial rescue could be partly due to insufficient levels of the precursor xanthoxin in these tissues or reflect the absence of *AAO* expression in these tissues (**Fig. 2**), which would prevent the local synthesis of bioactive ABA. These findings raise the possibility that AB-ald moves to other sites, such as the bundle sheath, epidermis, or vasculature, where *AAOs* are expressed and active. Bioactive ABA could then be synthesized and transported to guard cells to mediate stomatal closure. Similarly, we observed that the *aao3-4* mutant phenotype could be rescued by expressing *AAO3* in the bundle sheath or epidermis, tissues in which *ABA2* is not expressed, supporting the hypothesis that AB-ald is mobile. In this case, the precursor likely moves from its site of production in the phloem to distal sites where *AAO3* is present, allowing local ABA synthesis and downstream signaling.

Our findings are further supported by assays in *Xenopus oocytes*, which suggest that while ABA readily enters cells under acidic apoplastic conditions, it cannot exit once in the neutral cytosolic environment (**Fig. 6B**). ABA is known to be a weak acid that is mostly uncharged when present in the relatively acidic apoplastic compartment of plants ^2^, but rapidly becomes ionized when entering the more neutral cytoplasm. Thus, the oocyte assays provide support for ABA being subject to an ion-trapping mechanism, indicating that ABA export is unlikely to rely solely on passive diffusion across membranes. Consequently, transporter-mediated ABA export is likely to contribute to intercellular ABA movement ^61,62^. In contrast, AB-ald displays unrestricted movement across a range of pH values, suggesting it may serve as a mobile intermediate (**Fig. 6C**). Furthermore, during water stress, a rising apoplastic pH ^63,65^ causes more ABA to remain in its deprotonated form, increasing its retention in the apoplast; however, AB-ald can still enter target cells to be converted to ABA by AAOs. Together, these data argue that intercellular and possibly long-distance movement of ABA and its precursor is essential for coordinating root and shoot responses during drought. ABA surrounding the mesophyll cells is rapidly imported by ABCG17 and ABCG18, which work redundantly to keep ABA away from its target sites (e.g., guard cells), thus serving as a storage mechanism to prevent activity under non-stress conditions ^31,32^. Our efforts to identify the routes of ABA export out of the phloem and import into the guard cells revealed that known transporters such as ABCG25 and ABCG40 are not solely responsible for delivering ABA to guard cells. Instead, multiple pathways, including apoplastic diffusion, symplastic movement, and active transport (unknown transporters), likely cooperate to ensure the effective redistribution of ABA from source to target tissues.

Are ABA synthesis and movement mechanisms conserved between roots and shoots? At the level of synthesis, the spatial map appears broadly conserved between both organs in *Arabidopsis*. *ABA2* functions as a bottleneck and is largely restricted to phloem companion cells, implying that ABA and/or AB-ald must move non-cell autonomously to reach their targets (guard cells in leaves and the epidermis/cortex in roots). In contrast, the mechanisms that deliver ABA are likely to differ at the functional transport level. In water-stressed roots (e.g., xerobranching), ABA can be co-mobilized with water as it redistributes toward outer root tissues ^6^. However, in leaves, water flux is strongly curtailed as stomata close, reducing bulk-flow transport. Despite this reduction in water movement, our modeling, physiological assays, and ABACUS2 sensor measurements demonstrate that the signal accumulates in epidermis and guard cells within 20–30 minutes of phloem ABA induction. This rapid buildup under limited water flow suggests that ABA movement in leaves is strongly dependent on active or facilitated transport mechanisms yet to be identified.

Our study also presents a computational model of ABA transport within the leaf. While hormone transport has been modeled extensively to understand patterning in the root, shoot apical meristem and leaf venation ^56,64,66,67^, equivalent models in other plant tissues have received far less attention. Existing leaf-focused models have instead largely emphasized carbon and water transport ^68,69^. By deriving tissue-scale transport parameters from an idealized cell-based framework, we put forward a mechanistic approach for representing transport processes within the complex cellular topology of the spongy mesophyll. Our leaf model predicts that mesophyll airspaces play a major role in determining the path ABA takes. Simulations suggest that the tightly packed cells beneath leaf veins facilitate rapid ABA delivery to guard cells, paralleling the role these structures play in rapid hydraulic responses ^70^. Intriguingly, angiosperm leaves exhibit up to fourfold higher vein and stomata densities than gymnosperm leaves ^71^. We conjecture that this distinct anatomical arrangement, combining efficient ABA transport pathways with high vein density, could provide an explanation of why angiosperms evolved the use of elevated ABA biosynthesis to rapidly close stomata, in contrast to the multiple hours required for a stomatal response in gymnosperm leaves.

ABA levels rise rapidly in *Arabidopsis* leaves upon stress (**Fig. 3C, O**) ^72^. One would expect a corresponding induction of ABA biosynthetic genes. However, we did not detect significant transcriptional changes in the final steps of ABA biosynthesis (**Fig. 2**). One possibility is that regulation occurs upstream of these steps. For example, *NCED3* is stress-induced ^72^, and increased xanthoxin production could elevate pathway flux without altering expression of the downstream enzymes. Indeed, our xerobranching RNA-seq data showing a rapid increase in *NCED3*, *NCED*5, and *NCED*9 expression support this hypothesis. A second possibility is that elevated ABA derives, at least in part, from ABA-GE hydrolysis mediated by the two BG enzymes ^43,74^. A third possibility is that key control points in ABA biosynthesis are regulated post-translationally, enabling rapid increases in ABA production without detectable changes in transcript abundance.

In summary, our findings demonstrate that ABA biosynthesis is highly tissue-specific and that both ABA and its precursor are mobilized to coordinate water stress-induced root and shoot responses. By combining spatial genetics, models and physiological assays, we reveal that ABA movement is not restricted to a single route but likely occurs through multiple, tissue-dependent mechanisms. These insights refine current models of ABA action and highlight the importance of spatial hormone dynamics and tissue anatomy in regulating environmental responses in plants.

## Materials and methods

### Plant material and growth conditions

All *Arabidopsis thaliana* lines used in this work are in Colombia background (Col-0 ecotype, Salk Institute La Jolla, CA, USA). Sterilized seeds were plated on Murashige & Skoog (MS) x 0.5 (Duchefa Biochemic) medium containing 1% sucrose and 0.8% plant agar (Duchefa Biochemic), pH was adjusted to 5.6-5.8 with 1 M KOH. For transgenic plant selection, antibiotics were added to a final concentration of 50 μg/ml kanamycin (Duchefa Biochemic), 100 μg/ml spectinomycin, 150 μg/ml gentamycin (Duchefa Biochemic), and 30 μg/ml hygromycin (Bio-Gold). Plates with seeds were stratified for 48 hours at 4 °C, then transferred to growth chambers (Percival CU41L5) at 21 °C, 100-120 µEm^-2^S^-1^ light intensity under long-day conditions (16 h light/8 h dark). For seed production, plant transformation, crossing, and soil pot assays, seeds were sown on wet soil. Plants were grown in growth rooms under long-day conditions at 21 °C. The following constructs were previously described: *pML1:H2B-GFP* ^73^, *pKST1:GFP* ^50^, *pSUC2:YFP*, *pCO2:YFP*, *pSCR:YFP*, *pUBQ10:YFP* ^75^.

### Seed sterilization

Seeds were sterilized by vapor-phase sterilization (chlorine fumes) for 2.5 hours in Eppendorf tubes in the presence of 100 ml of 11% sodium hypochlorite and 5 ml of 32% hydrochloric acid in a sealed desiccator.

### Bacterial material and growth condition

All bacteria were grown on LB agar media: 20 g of LB and 15 g bacteriological agar (DIFCO) was added to 1 L doubly distilled water and autoclaved for 20 minutes at 120 °C. Antibiotics were added according to the specific resistances of bacteria at final concentrations of 50 µg/ml kanamycin, 100 µg/ml ampicillin, 30 µg/ml hygromycin, 100 µg/ml spectinomycin, 25 µg/ml gentamycin, 10 µg/ml tetracycline, and 25 µg/ml rifampicin. Plasmids were multiplied in chemically competent *E. coli* strain DH5α and extracted with a GenElute plasmid mini extraction kit (Sigma-Aldrich) following the manufacturer’s protocol.

### *Agrobacterium* transformation

Electro-competent *Agrobacterium tumefaciens* strain GV3101 was incubated on ice with 100 ng plasmids for 2 minutes, then electroporated in a MicroPulser (BIO-RAD) (2.2 Kv, 5.8 ms). Bacteria were transferred immediately to 1 ml liquid LB and shaken for 2 hours at 28 °C. Subsequently, bacteria were plated on LB agar plates containing the relevant antibiotics for 2 days at 28 °C.

### Plant DNA extraction and PCR

A “crude” DNA extraction method was used to extract plant genomic DNA to be used as a template for the PCR for sequencing and genotyping purposes. A few young leaves from *Arabidopsis thaliana* (about 100 mg) were placed in a 2 ml round-tip Eppendorf tube and frozen in liquid nitrogen. The leaves were crushed using a tissue-lyser to a thin powder and were homogenized with 400 μl DNA extraction buffer (200 mM Tris-HCL, pH 7.5-8.0, 25 mM EDTA, 250 mM NaCl, 0.5% SDS). The tubes were vortexed for 5 seconds and centrifuged for 1 minute at 13,000 rpm in an Eppendorf mini centrifuge. The supernatant was transferred to a new tube and DNA was precipitated with 300 μl isopropanol and incubated for 5 minutes at room temperature, followed by centrifugation at 13,000 rpm for 5 minutes at room temperature. The pellet was washed with 400 µl 70% EtOH and centrifuged for 1 minute at 13,000 rpm at room temperature, and the DNA pellet was dried and resuspended in 100 µl doubly distilled water. DNA amplification for sequencing and cloning was done by PCR in a Sensoquest labcycler using the Taq Ready Mix (HyLabs) following the manufacturer’s protocol.

### Histochemical GUS staining

For histochemical detection of GUS activity, plant tissues were incubated for approximately 16 hours at 37 °C in 100 mM sodium phosphate buffer (pH 7.0) containing 0.1% Triton X-100, 1 mM 5-bromo-4-chloro-3-indolyl-β-D-glucuronic acid cyclohexylammonium salt (Sigma-Aldrich), 2 mM potassium ferricyanide, and 2 mM potassium ferrocyanide. Tissues were immersed in 70% ethanol until transparent. GUS-stained tissues were imaged using a Zeiss Stemi 2000-C stereomicroscope ^76^. Images were captured using ZEN software (Zeiss).

### Cross-sections

Leaf cross-sections were performed as previously described ^77^. Leaves (4-week-old) were fixed in 4% PFA for an hour, rinsed twice in 1× PBS embedded in 8% agarose and sectioned to 100 µM slices using a Leica VT1000S vibratome. Sections were counterstained with 0.1% calcofluor white in 1x PBS solution for 30 minutes. Next, the seedlings were washed in 1x PBS for 30 min with gentle shaking and then imaged using a Zeiss LSM 780 inverted microscope.

### Confocal imaging

#### PI cell wall staining

Seedlings were stained in 10 mg/L propidium iodide for 1 minute, rinsed, and mounted in water. Seedlings were imaged on a Zeiss LSM 780 laser scanning confocal microscope with the laser set at 514 nm for YFP and propidium iodide excitation.

#### Calcofluor white cell wall staining

Seedlings/ leaves were stained in 0.1% calcofluor white as previously described ^78^. Seedlings were imaged on a Zeiss LSM 780 laser scanning confocal microscope with the laser set at 514 nm for YFP and 405 nm excitation for calcofluor white. Emission filters used were 517–570 nm for YFP, 425-475 nm for calcofluor white. Image analysis and signal quantification were done using ZEN lite 2012 software.

#### ABACUS2-400n imaging

Fully expanded leaves were excised from mature plants and mounted by their petioles on a 3D printed treatment chamber. Vacuum grease was used to keep the petiole separate from the leaf blade. The petiole was submerged in deionized water and cut again to prevent air bubbles from entering the vasculature ^21^. For time-course imaging, leaves were placed on an inverted Zeiss Airyscan 880 Multiphoton stage, in a chamber, and illuminated with a white lamp between imaging acquisitions to allow transpiration. For endpoint imaging, plants were mounted under a white lamp, a leaf disk was taken at 35 minutes of treatment, and imaged after 2 minutes of perfluorodecalin submergence ^79^. Plants were excited with a multiphoton 880nm laser to excite the CFP donor, and emission was captured at 465-500nm and 525-560 nm for donor and acceptor emission, respectively.

### Image analysis

ABACUS2-400n data was analyzed with the FRETENATOR2 beta (https://github.com/JimageJ/FRETENATOR2) using the *Segment and Ratio*, and *ROI labeller* plugins ^80,81^. Leaf epidermal cells and sub-epidermal cells, segmented using EZ-Peeler ^83^, https://github.com/JimageJ/EZ-Peeler.

### Thermal imaging

Thermal images were obtained using an FLIR T6xx series instrument. The camera was placed vertically ∼50 cm above the plants. Leaf temperature was quantified using FLIR TOOLS+ software, using a customized region-of-interest tool implemented according to the manufacturer’s instructions. Thermal imaging of tissue-specific CRISPR lines was performed on the T2 generation. Following imaging, plants were sprayed with Basta herbicide. Only plants that survived the Basta treatment, indicating that they contain the desired plasmid, were included in the quantitative analysis.

### Stomatal aperture measurements

Stomatal aperture was determined using a rapid and almost permanent imprinting technique previously described ^31^. Leaves of similar size were cut from 21-day-old plants. The abaxial side of the leaves was attached to 0.5-ml silicone impression material (elite HD+, Zhermack Clinical). The leaves were removed after the silicone impression material dried. Subsequently, transparent nail varnish was placed on the epidermal side. The samples were placed on slides with coverslips and imaged with a light microscope. The stomatal aperture size was quantified using Fiji software.

### Stomatal conductance measurements

Stomatal conductance was measured for 45-day-old *pRPS5:sgRNA-ABA2/AAO3* plants and respective controls using Leaf Porometer (LI-COR600) under room temperature conditions. 1-2 leaves per plant were measured.

### Drought assays

∼20-day-old plants were subjected to drought, in which the plants were not watered for 7-10 days. After the drought period, the soil moisture was measured using an electrode (ZSC Bluetooth sensor interface) to validate that the water content in the soil is at least 50% lower than the watered condition.

### β-estradiol treatment

β-estradiol (E2758, Sigma-Aldrich) was dissolved in ethanol to 10 mM and stored at −20 °C. For treatments, 5 µM β-estradiol with 0.01–0.02% Tween 20 was used as the final working solution. Seven-day-old seedlings were transplanted to soil and sprayed with 5 µM β-estradiol. β-estradiol was then applied twice weekly by spraying.

### Mutant lines

T-DNA insertion lines for single mutants were ordered from *The Arabidopsis Information Resource* (www.arabidopsis.org/): *aao1-2* (SALK_069221), *aao2-1* (SALK_104895), *aao3-4* (SALK_072361), *aao4-2* (SALK_057531) (**Sup. Table S3**).

### Constructs Cloning

#### Cell type-CRISPR cloning

The cell type-specific CRISPR vectors were generated as described previously ^82^. Next, the sgRNA/multiplex of interest was cloned using BsaI sites on the edges, followed by a Golden Gate reaction.

##### pSUC2:XVE:ABA2

As described in Mehra et al. ^6^, ABA2 was amplified from cDNA (using primer Fw-<u>CACC</u>ATGTCAACGAACACTGAATCTTCT and primer Rev: TCATCTGAAGACTTTAAAGGAGTG) and was cloned into pENTR/D-TOPO (Invitrogen K2400). MultiSite Gateway technology (Invitrogen) was used to combine the inducible promoter (*p1R4-pAtSUC2:XVE*) ^84^, ABA2, and a terminator (nosT) ^84^ in Gateway entry clones with Gateway-compatible binary destination vector pH7m34GW ^85^ in a MultiSite LR Clonase reaction. The recombination reaction was carried out according to the MultiSite Gateway Three-Fragment Vector Construction Kit manual (Thermo Fisher Scientific; catalog no. 12537-023)

#### Tissue-specific rescue

##### *aba2-1* rescue lines

*ABA2* in pENTR/D-TOPO was cloned into *pH7WGY2* destination vector using LR Gateway reaction (Invitrogen 11791). The YFP-ABA2 fragment was amplified (using primer Fw-<u>ACGGTCTCAATTG</u>ATGGTGAGCAAGGGC, primer Rev: <u>CAGGTCTCTAAAC</u>TCATCTGAAGACTTTAAAGGAGT) and cloned into the following plasmids: *pSUC2* in pBIN; *pSIG6* in pG0229-T; *pARSK1* in pG0229-T; *pML1* in pG0229-T; *pSCR* in pG0229-T; *pKST1* in pG0229-T and *pCORI3* in pG0229-T ^86^.

##### *aao3-4* rescue lines

*AAO3* CDS was synthesized by TWIST directly into the pENTR vector and cloned using the LR Gateway reaction (Invitrogen 11791) into pGWB504. Then, pSUC2, pSCR, and pML1 were amplified from plasmids ^86^, and cloned into the vector using restriction enzymes, generating the following constructs: *pSUC2:AAO3-GFP, pSCR:AAO3-GFP, and pML1:AAO3-GFP*.

#### Cloning of *ABA2, AAOs* reporter lines

*ABA2, AAO1, AAO2*, and *AAO4* promoters were synthesized by TWIST directly into the pENTR vector and cloned using the LR Gateway reaction (Invitrogen 11791) into the *pMDC7* vector for NLS-YFP reporters and the *pGWB3* vector for GUS reporters. *AAO3* promoter was amplified from genomic DNA and inserted using restriction enzymes into pG0229-T empty plasmid. Next, the NLS-YFP and GUS sequences were amplified from *pMDC7* and *pGWB3* vectors using primers containing BsaI sites. Then these sequences were cloned into the destination plasmid using Golden-Gate reaction. All primers are listed in Supplementary Table 2.

### Complementation lines cloning

#### pABA2:YFP-ABA2

*pABA2* was amplified using AscI and BamHI restriction sites and inserted to a *pBIN* empty vector. *YFP-ABA2* sequence was amplified from a plasmid described above using BsaI sites and inserted to the final destination using Golden Gate reaction.

#### pAAO3:AAO3-GFP

*AAO3* CDS was synthesized by TWIST directly into the pENTR vector and cloned using the LR Gateway reaction (Invitrogen 11791) into *pGWB504*. Then, *pAAO3* was amplified from genomic DNA, and cloned into the vector using SbfI and BamHI restriction enzymes.

### Hormone diffusion assays in *Xenopus* oocytes

Ovarian lobes from *Xenopus laevis* were kindly provided by Prof. Stephan Pless at the Department of Drug Design and Pharmacology, University of Copenhagen. The ovarian tissue was manually dissected into smaller pieces using tweezers and washed four times in Kulori buffer (90 mM NaCl, 1 mM KCl, 1 mM MgCl₂, 1 mM CaCl₂, 5 mM HEPES, pH 7.4), followed by four washes in OR2 buffer (82.5 mM NaCl, 2 mM KCl, 1 mM MgCl₂, 5 mM HEPES, pH 7.4). Defolliculation was performed by incubating approximately 5 mL of ovarian tissue with 80–100 mg collagenase (Collagenase NB 4G Proved Grade, Cat. No. S1746503, Nordmark) at a final concentration of 2.5 mg/mL. The tissue was agitated on a roller table (IKA® ROLLER 10 digital) at 50 rpm for 50–70 minutes. After enzymatic treatment, oocytes were washed four times in OR2 buffer and subsequently four times in Kulori buffer. *X. laevis* oocytes stage V-VI were selected from collagenase-treated oocytes for diffusion assays.

Selected oocytes were pre-incubated in Kulori buffer pH 5-7.2 (90 mM NaCl, 1 mM KCl, 1 mM MgCl_2,_ 1 mM CaCl_2,_ 5 mM MES/HEPES. MES for pH 5-5.8, HEPES for pH 6.2-7.2) for 2 min before transferring to compound containing Kulori buffer of same pH as pre-incubation for 1 h. Concentration of compounds was 100 µM. After 1 h, the oocytes were washed three times in Kulori buffer pH 7.4, followed by one wash in Milli-Q water. 1 µL buffer samples were taken after the addition of oocytes to compound containing buffer. Buffer samples were treated as oocyte samples.

Sample preparation: Single oocytes per sample were analysed; each oocyte was homogenized in 62.5 µL 50% methanol containing the internal standard Sinigrin (1.25 µM) and Caffeine (1 µM) and stored minimum 1 h at −20 °C. The homogenized oocyte samples were pelleted at max speed for 10 min at 4 °C. Then, 50 µL of the supernatant was mixed with 75 µL Milli-Q water to a final methanol concentration of 20% before filtration through a 0.22 µm filter plate.

### Metabolite analysis by LC-MS/MS

Samples were subjected to analysis by liquid chromatography coupled to tandem mass spectrometry. Chromatography was performed on a 1290 Infinity II UHPLC system (Agilent Technologies). Separation was achieved on a Kinetex XB-C18 column (100 x 2.1 mm, 1.7 µm, 100 Å, Phenomenex, Torrance, CA, USA). Formic acid (0.05%, v/v) in water and acetonitrile (supplied with 0.05% formic acid, v/v) were employed as mobile phases A and B, respectively. The elution profile for glucosinolates was: 0-0.2 min, 5 % B; 0.2-3.5 min, 5-65 % B; 3.5-4.2 min 65-100 % B, 4.2-4.9 min 100 % B, 4.9-5.0 min, 100-5% B and 5.0-6.0 min 5 % B. The mobile phase flow rate was 400 µL/min. The column temperature was maintained at 40 °C. The liquid chromatography was coupled to an Ultivo Triplequadrupole mass spectrometer (Agilent Technologies) equipped with a Jetstream electrospray ion source (ESI) operated. The instrument parameters were optimized by infusion experiments with pure standards (**Sup. Fig. S36**). The ion spray voltage was set to 4500 V in negative ion mode and 3000 V in positive ion mode, respectively. Dry gas temperature was set to 325 °C and dry gas flow to 12 L/min. Sheath gas temperature was set to 400 °C and sheath gas flow to 12 L/min. Nebulizing gas was set to 50 psi. Nitrogen was used as dry gas, nebulizing gas and collision gas. Multiple reaction monitoring (MRM) was used to monitor precursor ion → fragment ion transitions. MRM transitions were determined by direct infusion experiments of reference standards. Detailed values for mass transitions can be found in **Sup. Table S4**. Both Q1 and Q3 quadrupoles were maintained at unit resolution. Mass Hunter Quantitation Analysis for QQQ software (Version 10, Agilent Technologies) was used for data processing. Linearity in ionization efficiency was verified by analyzing dilution series that were also used for quantification.

### Oocyte data analysis

Oocyte data analysis was performed using RStudio. Outliers were removed automatically using a custom R function that excludes values outside ±1.5 times the interquartile range. The response factor for oocytes was determined by standard curves for the compound investigated and the internal standards (Sinigrin used for ABA and ABA-GE, Caffeine used for AB-Aldehyde): response factor (f) = slope of internal standard / slope of compound investigated. Concentration in samples was calculated as follows: (Area of peak for compound / Area of peak for internal standard) × f × final internal standard concentration × dilution factor. Dilution factor: 156.25; final internal standard concentration: 0.5 µM (Sinigrin) and 0.4 µM (Caffeine).

### Mathematical model

Full details of the mathematical model are provided in the Supplementary Appendix Text. This text details the cell-based model and derives the equations governing the corresponding continuum approximation (sections 1-4). The text details how we integrated the localization of the biosynthesis enzymes (section 5), how we calculated the cell-to-cell contact lengths using data on the porosity of the spongy mesophyll (section 6), how we estimated the membrane permeabilities using the experimental data from the oocyte experiments (section 7) and how these membrane permeabilities and the plasmodesmatal permeabilities contribute to the effective cell-to-cell permeability (section 8). The model is programmed using Matlab; code is provided at https://gitlab.com/leahband/ABA_Arabidopsisleaf_waterstress

### Leaf micro-CT images

Mature Arabidopsis leaves were imaged at a spatial resolution of 2.5 microns using a v|tome|x M X-ray Tomography System (Waygate Technologies (a Baker Hughes Company), Germany) at The Hounsfield Facility, University of Nottingham, following a modified method of ^59^. A 5 mm diameter disk was cut from an Arabidopsis leaf and suspended in polystyrene in a 0.2 ml microcentrifuge tube attached to a plastic rod. The sample was allowed to acclimatize for 20 mins before imaging in the micro-CT system. The CT scan was conducted using the nano-focus transmission X-ray tube and a Dynamic41|100 detector panel. X-ray tube energy and current was 75 kV and 105 µA, respectively. The scan collected 3600 projection images over a 360° rotation of the sample in the x-ray beam in ‘continuous’ acquisition mode with no image averaging. Detector timing was 334 ms per projection. No X-ray tube filters were used.

Projection images were reconstructed to 3D volume using datos|REC (Waygate Technologies (a Baker Hughes Company), Germany). Images and porosity were prepared using VGStudioMAX (Hexagon AB, Sweden).

### ABA and related molecules measurements

In the present work, a targeted liquid chromatography-tandem mass spectrometry (LC-MS/MS) analysis was applied to quantify differential abundances (relatively) of selected phytohormones ABA, AB-ald and ABA-GE. For that purpose, three examples of leaf samples: wild type *(Col-0)* and two genetic mutants *aba2-1* and *4aao* were subjected to the extraction procedure adopted from ^87^. Chromatographic conditions were based on the method described ^88^ with minor modifications.

Watered and drought samples for LC-MS/MS were preliminarily frozen in liquid nitrogen and lyophilized. The obtained material was ground to powder with 5-mm stainless steel beads in a Qiagen Tissue Lyser II for 2 min at 25 Hz. Approximately 100 mg of ground leaves was weighed into 2 mL sample tubes and added with 400 μL of extraction buffer (10% methanol, 1% acetic acid) spiked with 100 ng/mL of isotopically labeled standards ABA-D6 acid and D5-ABA-GE (IS). Tubes were mixed and shaken, then incubated on ice for 30 min and centrifuged for 10 min at 16,000 × g and 4 °C. Supernatants were transferred into fresh tubes and centrifuged two more times to remove all debris prior to liquid chromatography with tandem mass spectrometry (LC-MS/MS) analysis. Aliquots of 200 ul were transferred to HPLC vials with inserts. An extraction control not containing plant material was treated equally to the plant samples

Method development and LC-MS/MS analysis were performed on a Thermo Scientific Vanquish Flex UHPLC System coupled to a TSQ Altis Triple Quadrupole mass spectrometer (Thermo Fisher Scientific, Waltham, MA, USA) with an electrospray EASY-Max NG ionization source (OPTON-30157). Chromatographic separation was carried out on a ACQUITY UPLC HSS T3 Column (100Å, 1.8 µm, 2.1 mm X 100 mm) (Waters, Milford, MA, USA). Tuning and data acquisition were carried out using Xcalibur and quantification using TraceFinder 5.2 version (Thermo Scientific™).

LC-MS grade Methanol was obtained from Supelco (Merck, Darmstadt, Germany). Water (18.2 MΩ·cm) was purified in a Milli-Q device with LC-Pak, Millipore Purification System (conductivity 0.055 μS cm−1 at 25 °C, resistivity equals 18.2 MΩ·cm) (Merck, Darmstadt, Germany). Acetic acid (LC-MS grade) was obtained from Sigma-Aldrich (St. Louis, MO, USA). For the detection and quantification of selected compounds, a triple quadrupole mass spectrometer (MS) was operated using electrospray ionization (ESI) in negative mode. Compound-specific scheduled selective reaction monitoring (t-SRM) transitions were optimized based on built-in optimization procedure injecting authentic standards. All molecules were monitored as [M-H]^−^ ion according to their m/z values. In **Sup. Table S5** summarized final SRMs. MS source parameters were maintained for maximum sensitivity and set up as follows: ESI spray voltage in the negative-ion mode, 3.4 kV; sheath gas flow-rate, 50 arb; auxiliary gas flow-rate, 9 arb;; sweep gas flow-rate, 1 arb; ion transfer tube temp, 325 °C; vaporizer temp, 350 °C.

Chromatographic elution was carried out on isocratic form with constant ratio of methanol:water:acetic acid(45:54:1,v/v/v)as mobile phase at flow rate 0.30 ml/min. The column temperature was maintained at 35 °C. Injection volume was 2 ul. The retention times were about 3.43 min for AB-Ald, 4.87 min for ABA, and 1.92 min for ABA-GE. The analyte and its IS co-eluted in the same or similar time (Sup. Fig. S27).

Individual and combined stocks of phytohormones were prepared in methanol:water (50:50) with 0.1% acetic acid. To verify specificity and reproducibility of analytes a series of dilutions in the range of 1 to 2,000 ng/mL, prepared in extraction solvent containing deuterated internal standards, and were used to build up calibration curves. Limit of quantification (LOQ) was estimated as (signal/noise) >10 and calculated based on the following equations LOQ = SD·10/S (a). Retention time and peak areas were investigated by repeated injection (n = 3) of each dilution. Resulted linearity range, correlation coefficients (R2), LOQ and corresponding relative standard deviations (RSDs) are presented in **Sup. Table S6**.

LC–MS/MS analysis was performed to elucidate differential abundance of ABA-related metabolites between watered and drought conditions. Relative abundances were calculated from integrated peak areas normalized to isotopically labeled internal standards. ABA and ABA-GE were normalized to their matched labeled analogs (ABA-d6 and d5-ABA-GE, respectively), whereas AB-ald, lacking a compound-specific labeled standard, was evaluated using surrogate internal standard normalization (geometric mean of internal standards) ^89,90^. Group comparisons were conducted using statistical analyses on log2-transformed relative abundances using Student’s t-tests, and fold changes were calculated by back-transforming mean log2 differences to a linear scale (n=3). AB-Ald levels were significantly higher in the irrigated condition than in the drought (fold change = 2.56, p < 0.04). Similarly, ABA exhibited a moderate difference between groups (fold change = 1.42, p ≈ 0.05). ABA-GE also demonstrated differential relative abundance, but it did not reach statistical significance (fold change = 1.24). The visualization of results is shown in Fig. 3A. (a) ICH harmonized tripartite guideline: validation of analytical procedures: text and methodology Q2(R1), International Conference of harmonization of technical requirements for registration of pharmaceuticals for human use 2005. A revised/renewed draft version of this guideline: ICH guideline Q2(R2) on validation of analytical procedures 2022.

### Bioinformatics

DNA sequence alignment was performed using the SnapGene alignment software (http://www.snapgene.com/) and the NCBI BLAST tool (https://blast.ncbi.nlm.nih.gov/Blast.cgi/). Gene sequences and relevant information were obtained from the *Arabidopsis* Information Resource (https://www.arabidopsis.org/).

### Spatial transcriptomics

Leaf samples were collected from 22-day-old plants and processed in accordance with Merscope guidelines for FFPE sections. To facilitate post-sectioning processing and obtain more data from a single section, leaves from the same plant were stacked, creating a layered arrangement of leaves in proximity. 4 μm sections from FFPE blocks were mounted on MERSCOPE glass slides (Vizgen, USA). The procedures of de-paraffinization, de-cross-linking, anchoring, gel-embedding, tissue clearing, and probe hybridization were performed according to the MERSCOPE V2.0 FFPE sample preparation user guide. Briefly, after gel embedding, tissues were cleared for a total time of 24 h at 47 degrees, with a 4h step in 5mg/ml proteinase K (New England Biolabs, USA) and then 20 h in 0.2 mg/ml proteinase K. Clearing continued for another 48h at 37 degrees, including a photobleaching step of 12 h during that time and digestion time of 2h. Next, sections were incubated with MERFISH probe mix for a minimum of 18 h at 47 degrees and enhancer probe hybridization for a minimum of 16 h at 37 degrees. A custom-made gene panel of 140 genes was used. After probe hybridization, the samples were labeled with DAPI and poly T and placed in the MERSCOPE flow chamber for imaging. The raw image files obtained were processed via the MERlin image analysis pipeline ^91^. The Cellpose2 algorithm was utilized to segment cells from the nuclear DAPI signal and poly T. The decoded RNA molecules were then partitioned into individual cells to generate single-cell count matrices. The sample data were segmented into areas of interest (ROIs), representing different tissue types, using the MERSCOPE Vizualizer software. For water stress comparison - mRNA reads for genes within each ROI were normalized to the relative prevalence in that ROI to mitigate differences in mRNA quality between spatial transcriptomics slides and samples.

### Expression profiling of ABA metabolism genes

Five-day-old seedlings were subjected to air-gap conditions (xerobranching), followed by recovery (reconnection with agar). Roots were allowed to grow to different lengths under air-gap (xerobranching) and recovery conditions. Samples were designated as Control, XB_1, XB_2, XB_3, XB_5, Rec_1, and Rec_2. Here, XB_1, XB_2, XB_3, and XB_5 correspond to root tips that grew 1, 2, 3, or 5 mm, respectively, within the air gap, while Rec_1 and Rec_2 represent roots that grew 3 mm and 5 mm, respectively, after reconnection to agar. For transcriptomic analysis, the apical 2 mm of root tips were harvested. Each biological replicate contained root tips from approximately 100 seedlings. Four biological replicates were prepared for all samples, except for the Control, which had three replicates.

Total RNA was extracted using the RNeasy Kit (Qiagen) and subjected to bulk RNA sequencing as previously described ^92^, with a sequencing depth exceeding 60 million paired-end reads per sample. Reads were aligned to the Arabidopsis thaliana reference genome (TAIR10). Differential gene expression (log₂ fold-change) was calculated relative to control, with a significance threshold of p ≤ 0.05.

## Acknowledgments

We thank Yakir Divald for graphical assistance. This work was supported by the Israel Science Foundation (1462/24, 2712/24 and 1346/25 to E.S.), the European Research Council (101118769-HYDROSENSING to E.S, M.J.B and T.H), the Hounsfield Facility, University of Nottingham received funding for the v|tome|x CT System from Natural and Environment Research Council (NE/X005801/1 to C.J.S.), the Royal Society (URF/R1/241367 to J.R.), BBSRC Discovery fellowship (BB/Z514482/1 to P.M.), Horizon Europe ERC Starting grant (101161820-WATER-BLIND to P.M.), Biotechnology, the Novo Nordisk Foundation (NNF23OC0082218 to C.K.) and Biological Sciences Research Council (BBSRC) (BB/V003534/1, BB/W015080/1, BB/W008874/1 to M.J.B.).

