## Supplementary material for "Water stress adaptive responses in plants require movement of ABA and AB-aldehyde from vascular to target tissues": Sup Data

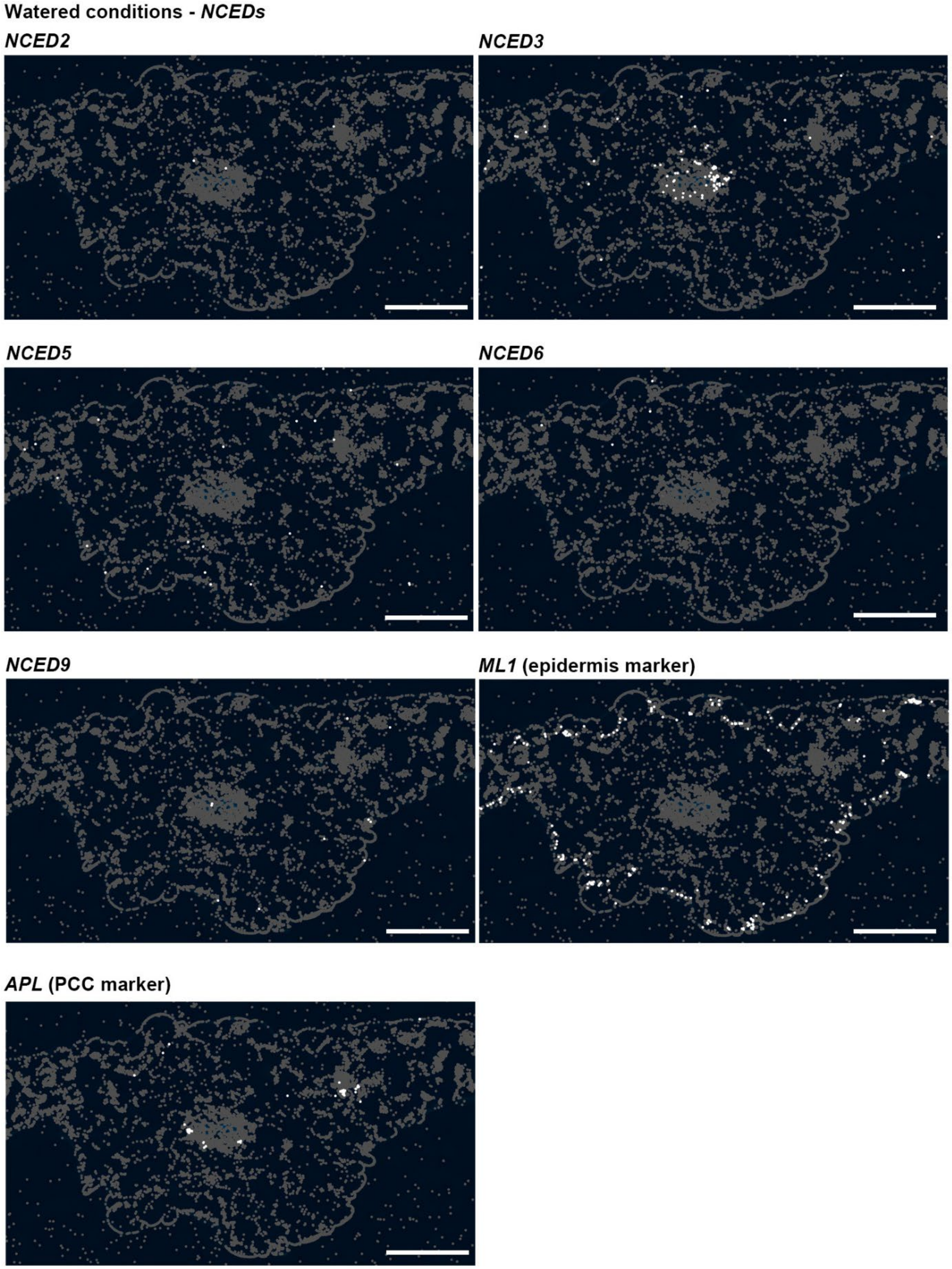


**Sup. Fig. S1. *NCEDs* are mainly expressed in the leaf vasculature and mesophyll.** Leaf spatial transcriptomics for *NCED* genes. Marker genes: *ML1* epidermis, *APL* phloem companion cells. Scale bar = 250 µm.

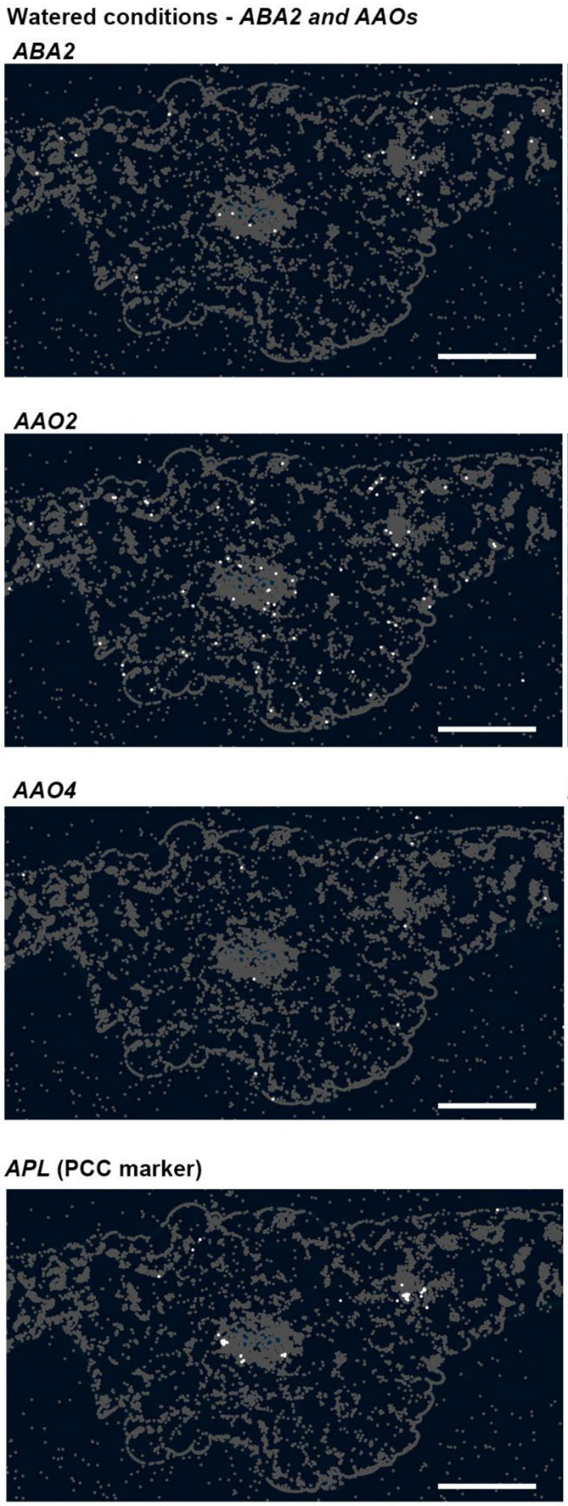


**Sup. Fig. S2. *ABA2* and *AAOs* are mainly expressed in the leaf vasculature and epidermis.** Leaf spatial transcriptomics for *ABA2* and *AAO* genes. Marker genes: *ML1* epidermis, *APL* phloem companion cells. Scale bar = 250 µm.

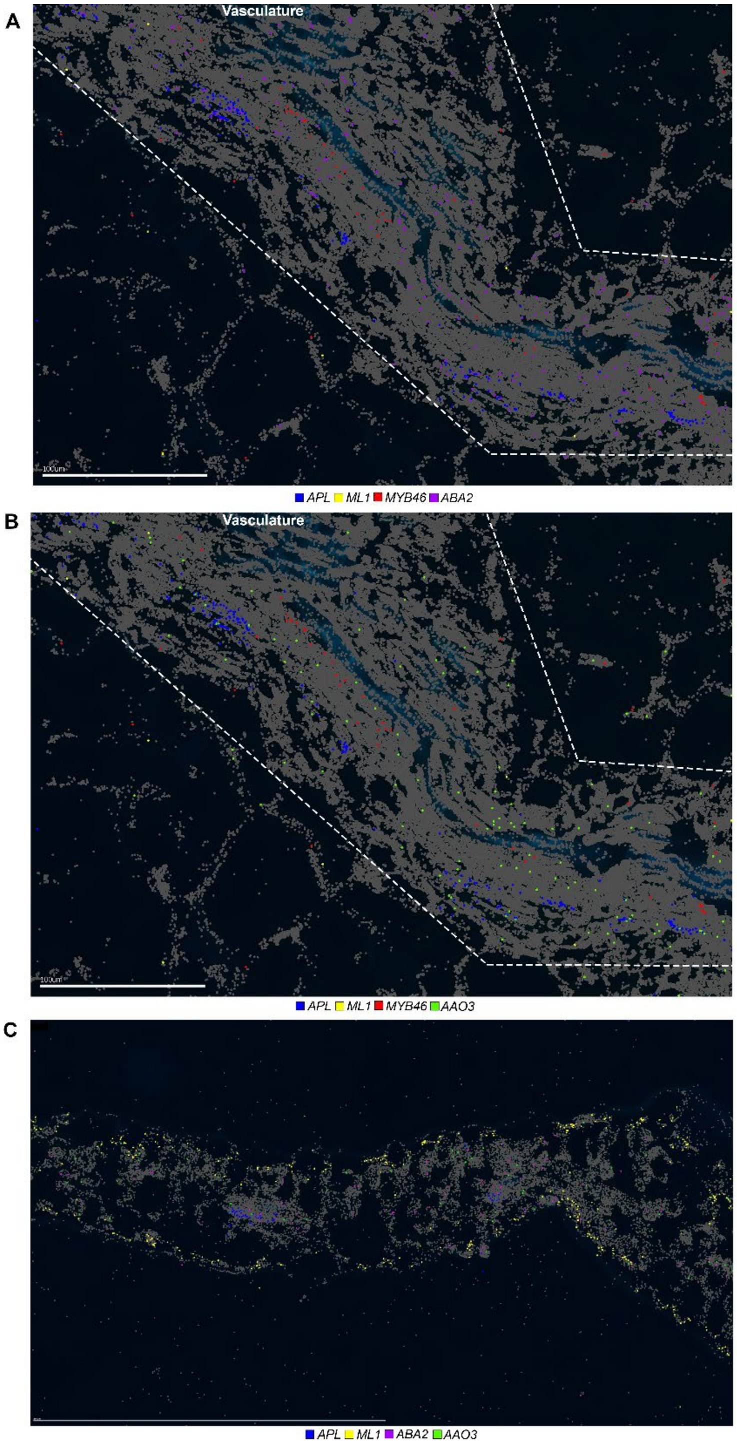


**Sup. Fig. S3. *ABA2* and *AAO3* expression beyond the phloem companion cells. (A-B)** Leaf spatial transcriptomics for *ABA2* **(A)**, *AAO3* **(B)** in a developing leaf sectioned near the vegetative shoot apical meristem. The vasculature is highlighted. Marker genes: *ML1* epidermis, *APL* phloem companion cells, *MYB46* differentiating xylem cells. Blue autofluorescence marks the xylem. *ABA2* and *AAO3* expression is not detected outside the vasculature. Scale bar = 100 µm. **(C)** Leaf spatial transcriptomics for *ABA2* in a mature leaf that is significantly different than the other leaves, showing a stronger signal throughout all transcripts. Marker genes: *ML1* epidermis, *APL* phloem companion cells. Scale bar = 250 µm.

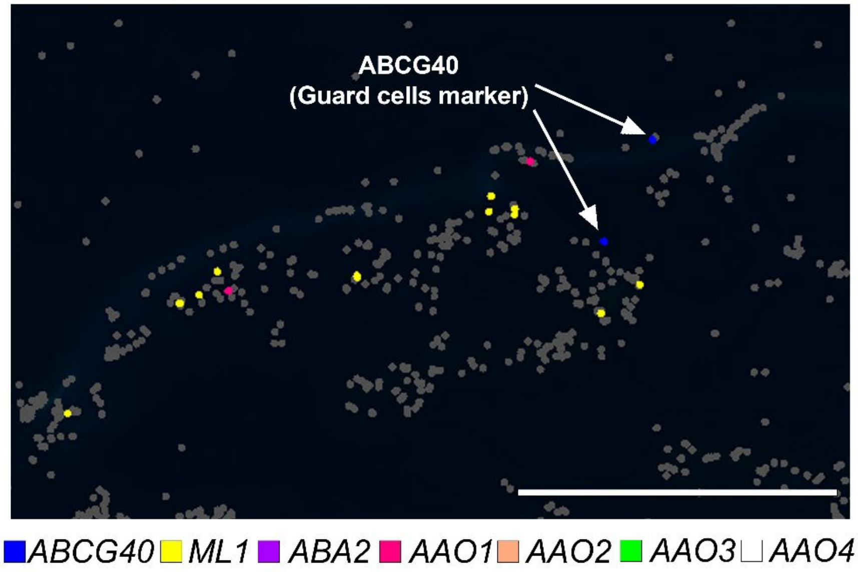


**Sup. Fig. S4. *ABA2* and *AAOs* are not detected in proximity to the guard cells ABCG40 marker.** Leaf spatial transcriptomics for *ABA2* and *AAOs*. Marker genes: *ML1* epidermis; *ABCG40* guard cells. Scale bar = 50 µm.

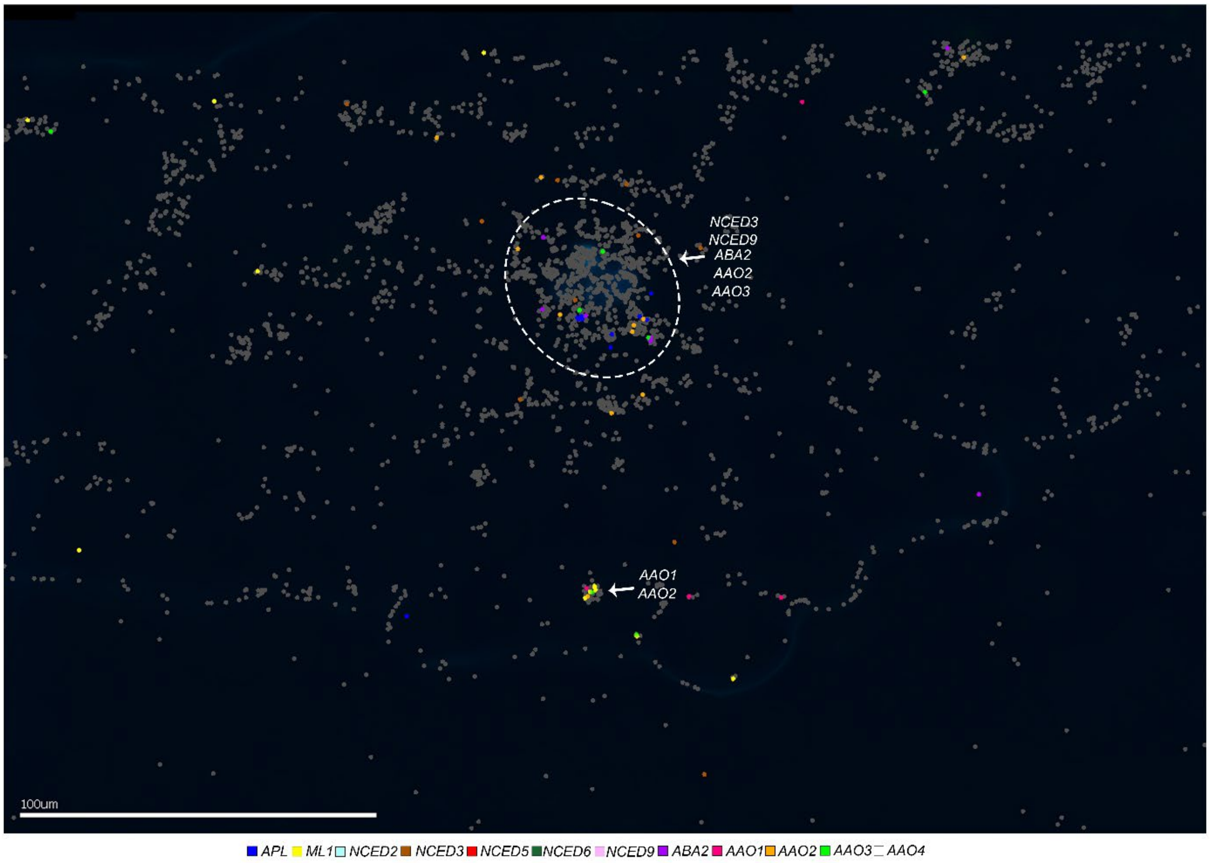


**Sup. Fig. S5. *NCEDs, ABA2,* and *AAOs* expression under water stress.** Leaf spatial transcriptomics for *NCED*, *ABA2,* and *AAO* genes after 10 days of drought stress (no irrigation). Marker genes: *ML1* epidermis, *APL* phloem companion cells. Scale bar = 100 µm. White dashed circles represent the vasculature. Genes written in white indicate the tissue in which they are expressed.

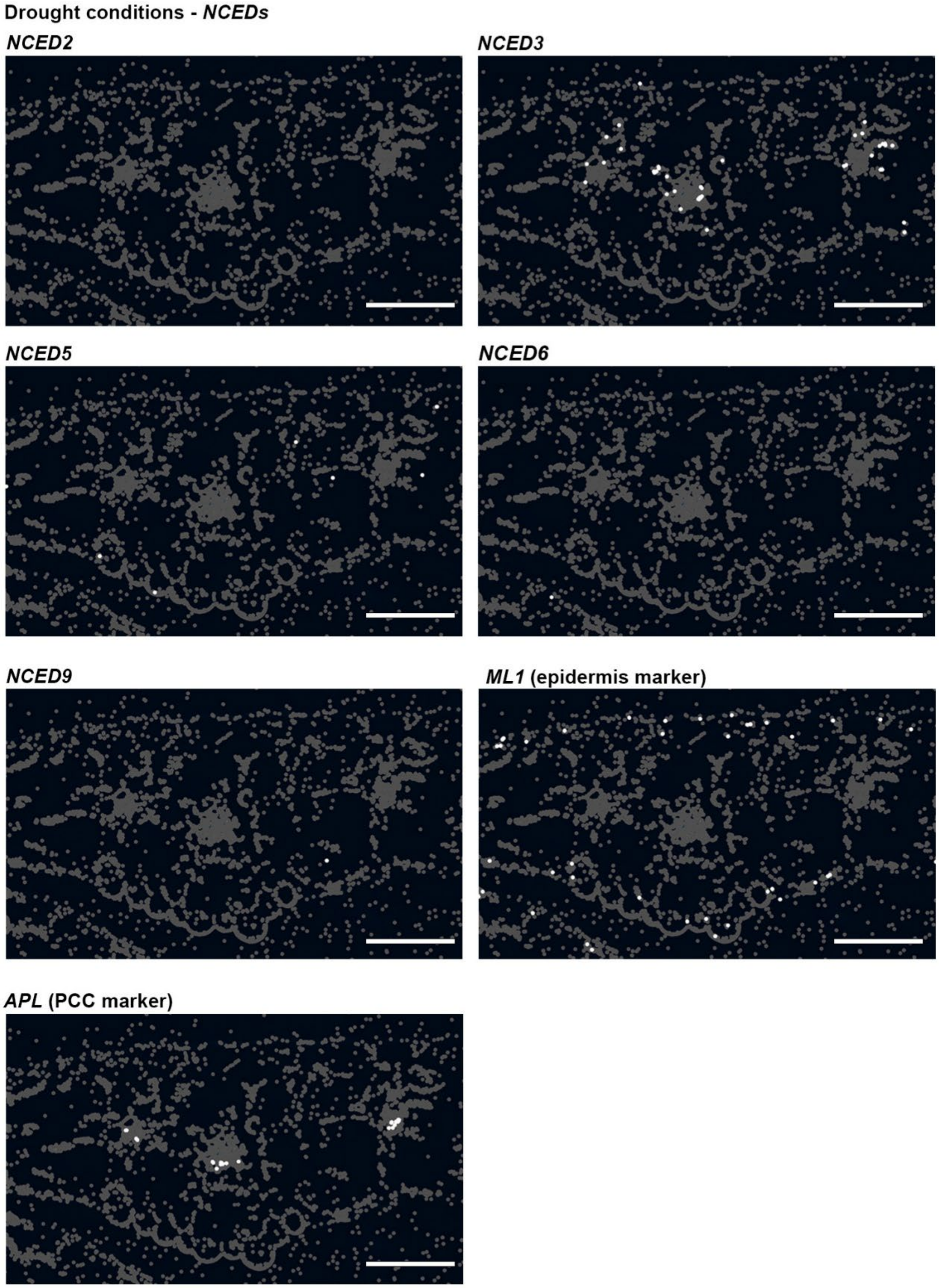


**Sup. Fig. S6. *NCEDs* are mainly expressed in the leaf vasculature and mesophyll after drought stress.** Leaf spatial transcriptomics for *NCED* genes after 10 days of drought (no irrigation). Marker genes: *ML1* epidermis, *APL* phloem companion cells. Scale bar = 100 µm.

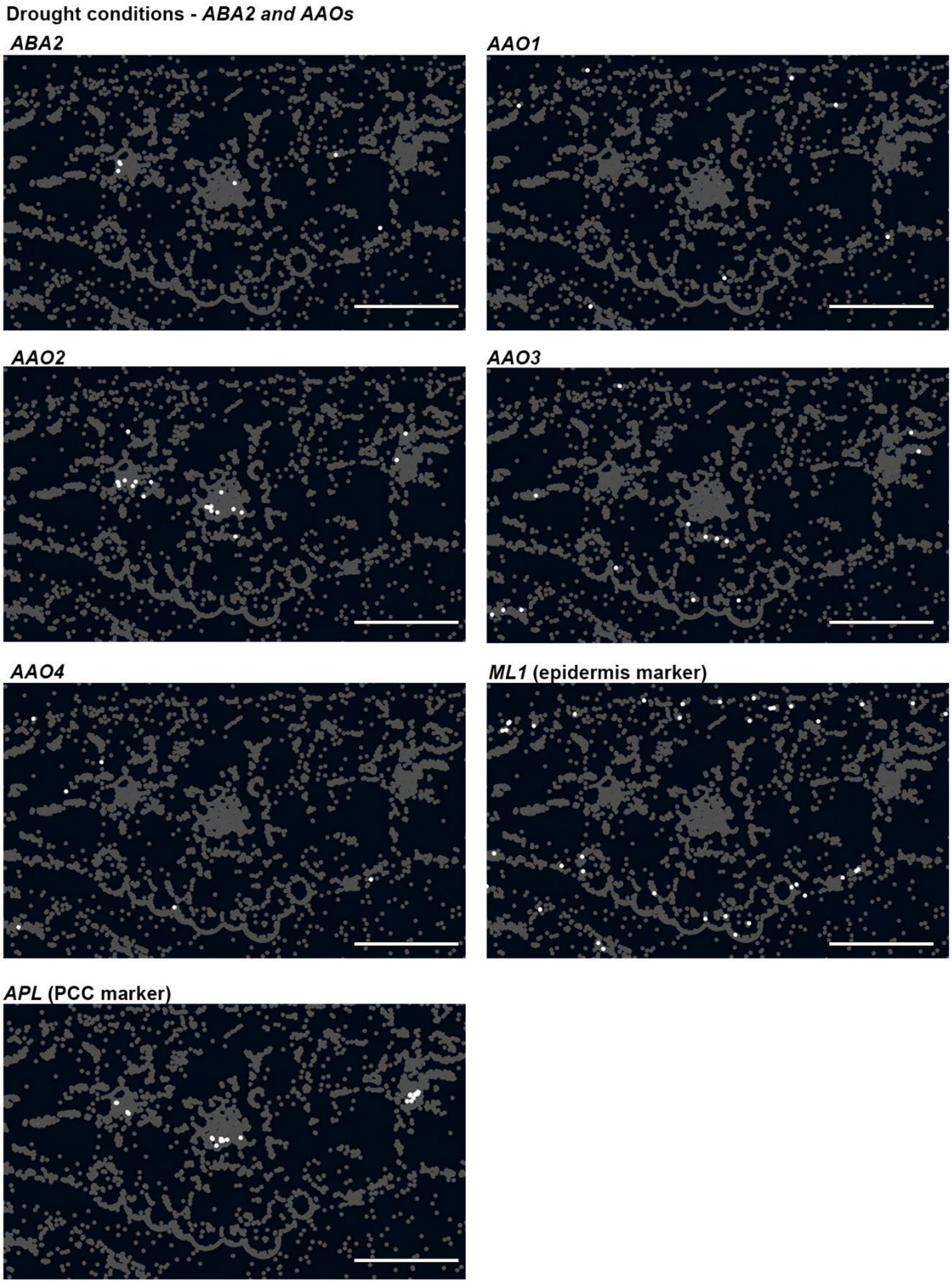


**Sup. Fig. S7. *ABA2* and *AAOs* are mainly expressed in the leaf vasculature and epidermis after drought stress.** Leaf spatial transcriptomics for *ABA2* and *AAO* genes after 10 days of drought (no irrigation). Marker genes: *ML1* epidermis, *APL* phloem companion cells. Scale bar = 100 µm.

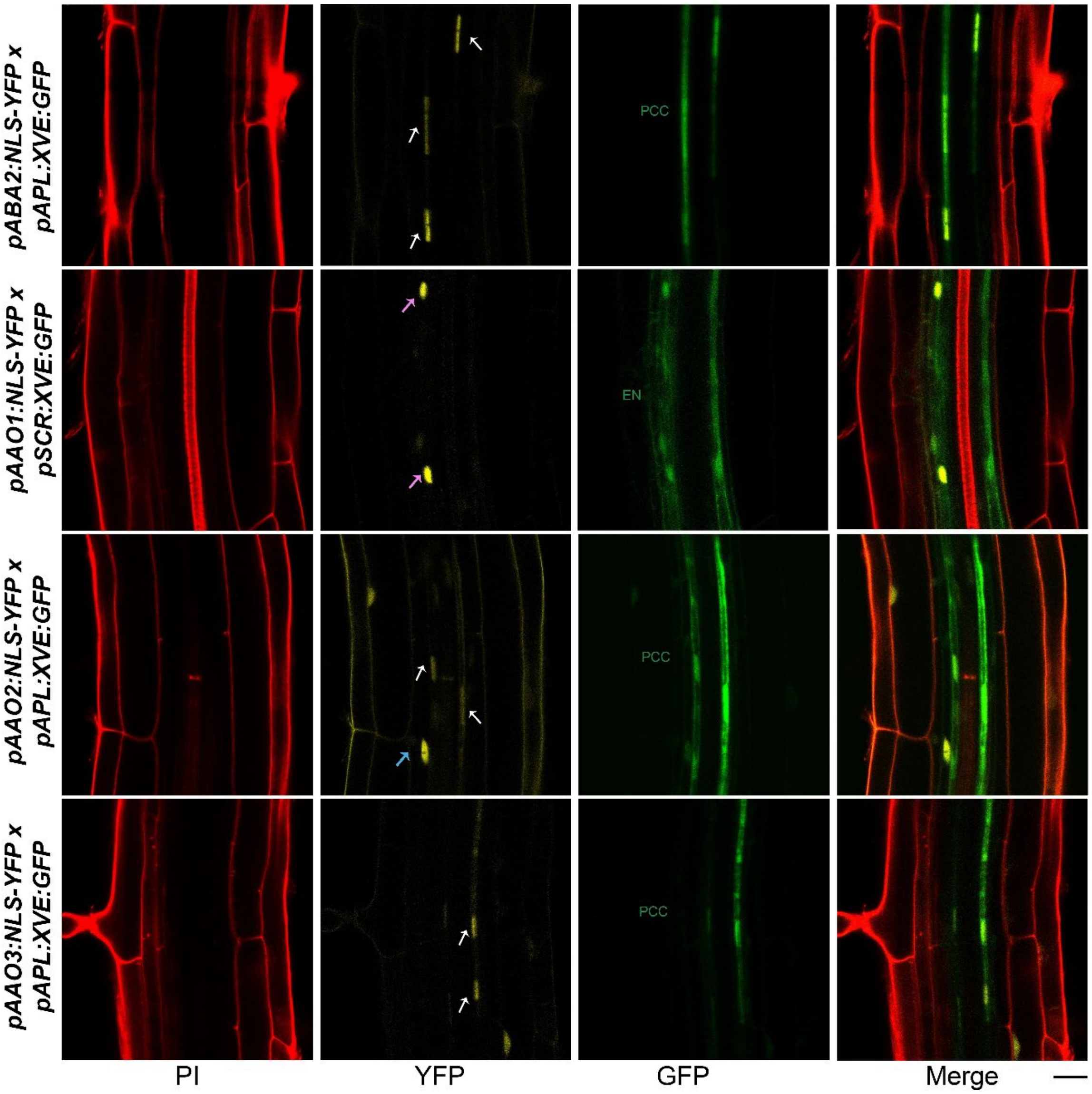


**Sup. Fig. S8. Characterizing *pABA2, AAO1, AAO2, and AAO3* expression patterns using genetic markers for the phloem and endodermis.** Representative confocal image of 7-day-old roots of *pABA2, AAO2,* and *AAO3:NLS-YFP* x *pAPL:XVE:GFP (APL;* phloem companion cells marker), and *AAO1:NLS-YFP* x *pSCR:XVE:GFP (SCR;* endodermis marker), stained with PI (red). White arrows indicate phloem companion cell expression, pink arrows indicate endodermis expression, and blue arrows indicate pericycle expression. Scale bar = 20 µM.

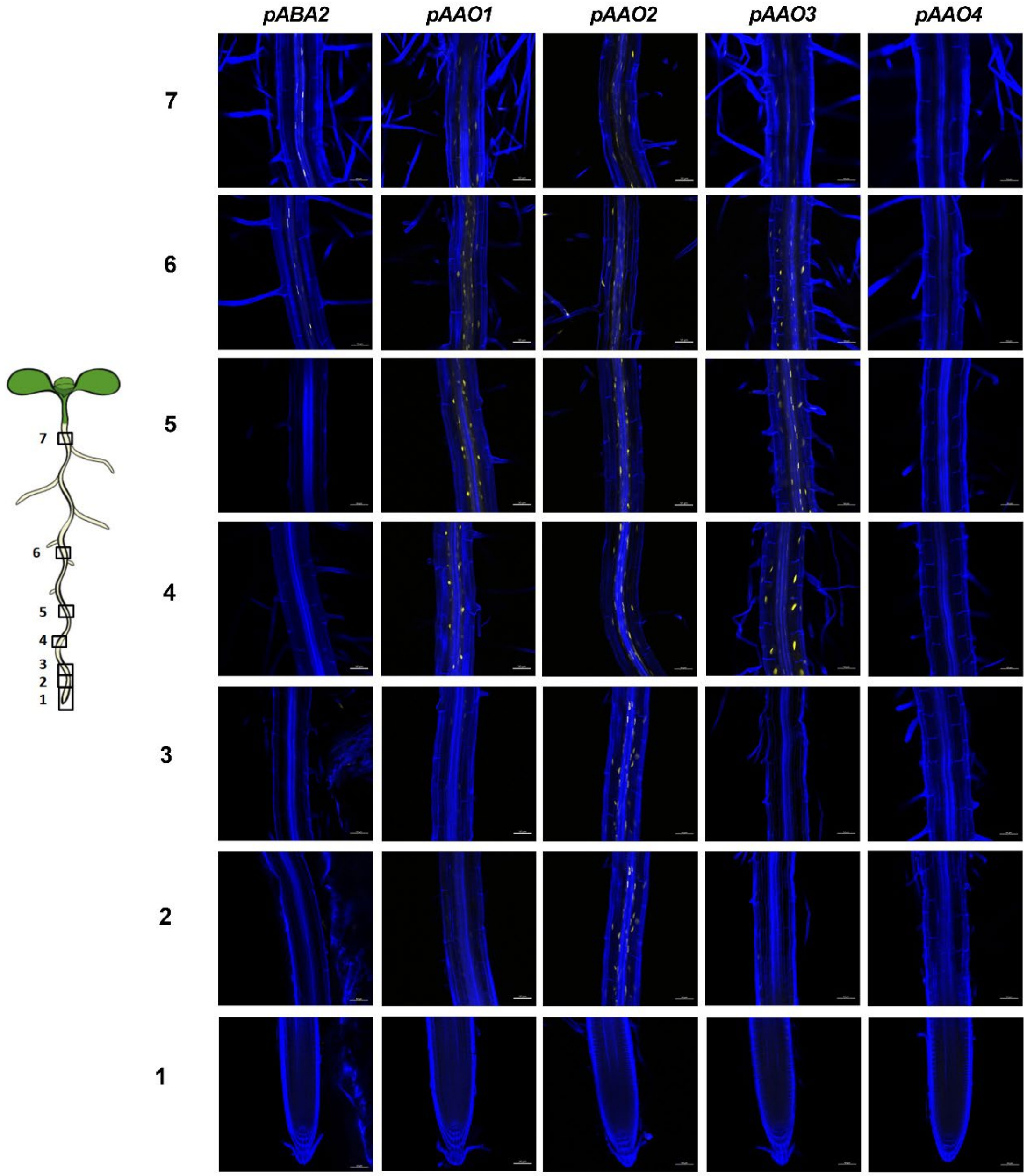


**Sup. Fig. S9. *ABA2* and *AAO*s expression in the root.** Shown are root images of 7-day-old *pABA2, pAAO1, pAAO2, pAAO3* and *pAAO4:NLS-YFP* plants. Scale bars = 50 µm.

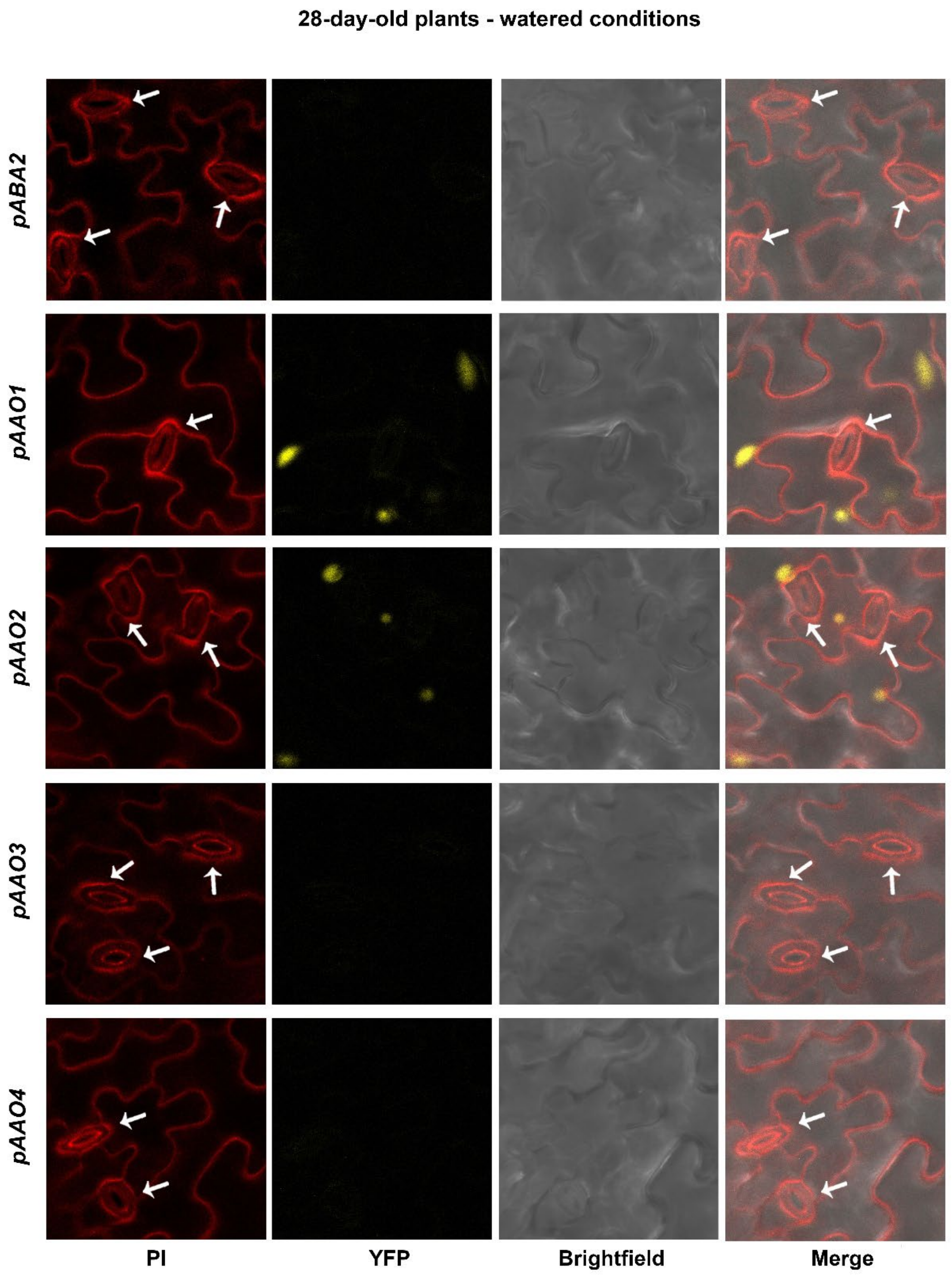


**Sup. Fig. S10. *ABA2* and *AAO*s expression is not detected in guard cells of 28-day-old leaves.** Shown are 28-day-old leaf images of *pABA2, pAAO1, pAAO2, pAAO3* and *pAAO4:NLS-YFP* plants. Scale bars = 20 µm. White arrows point at guard cells.

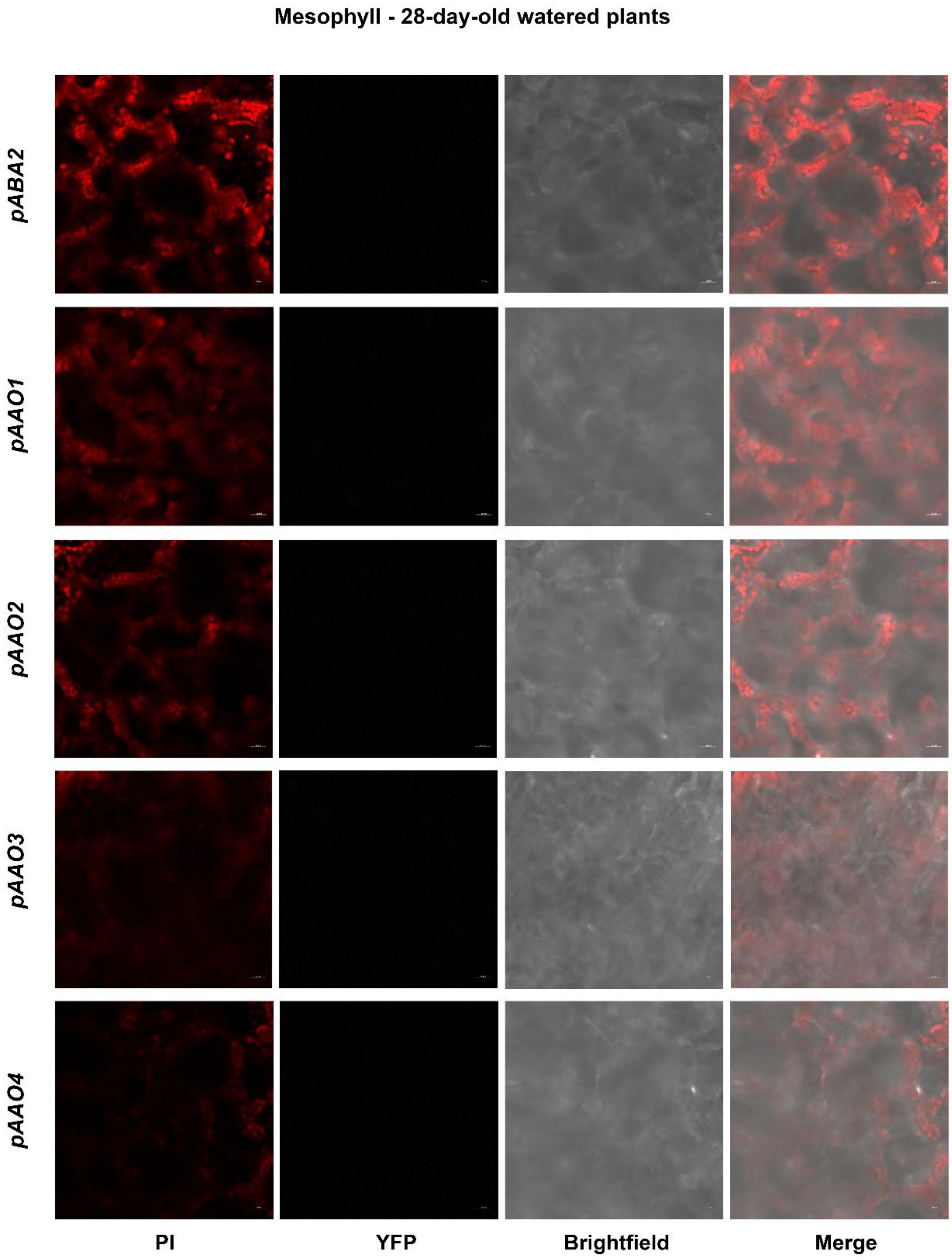


**Sup. Fig. S11. *ABA2* and *AAO*s expression is not detected in mesophyll cells of 28-day-old leaves.** Shown are 28-day-old leaf images of *pABA2, pAAO1, pAAO2, pAAO3* and *pAAO4:NLS-YFP* plants. Scale bars = 20 µm.

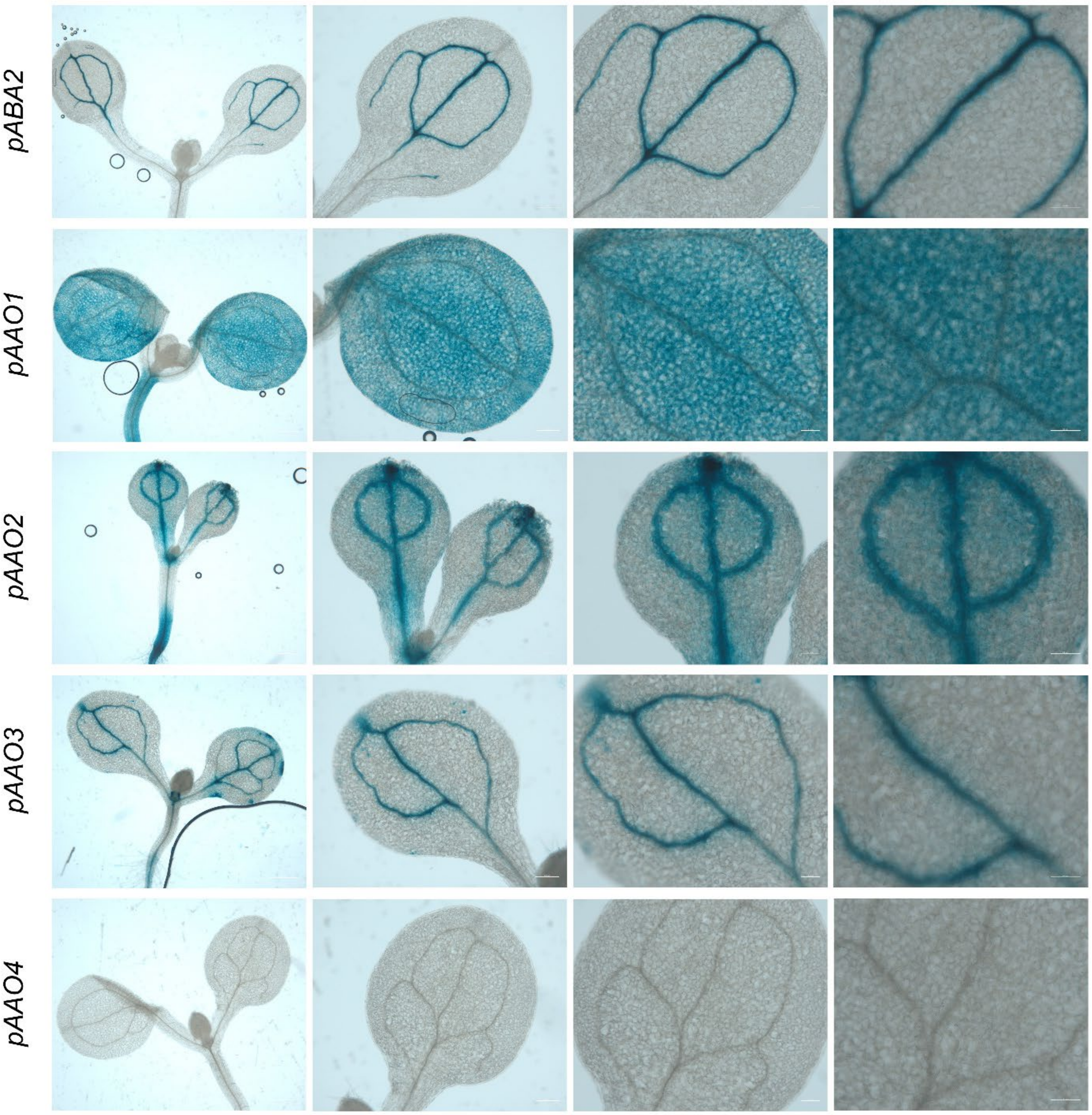


**Sup. Fig. S12.** ***ABA2* and *AAO*s expression in cotyledons.** Shown are cotyledon images of 7-day-old *pABA2, pAAO1, pAAO2, pAAO3* and *pAAO4:GUS* plants. Scale bars (left to right column) = 0.5 mm, 0.2 mm, 0.1 mm, and 0.1 mm.

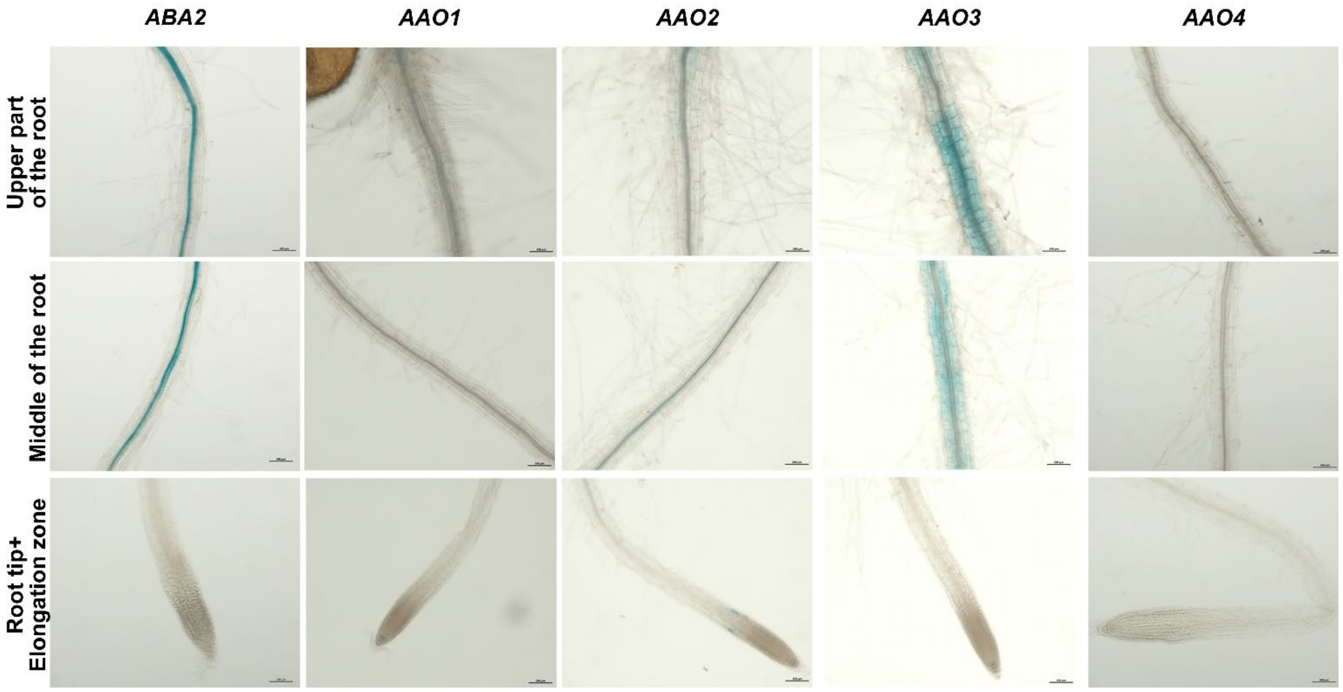


**Sup. Fig. S13. *ABA2* and *AAO*s expression in roots.** Shown are root images of 7-day-old *pABA2, pAAO1, pAAO2, pAAO3* and *pAAO4:GUS* plants. Scale bars = 0.1 mm.

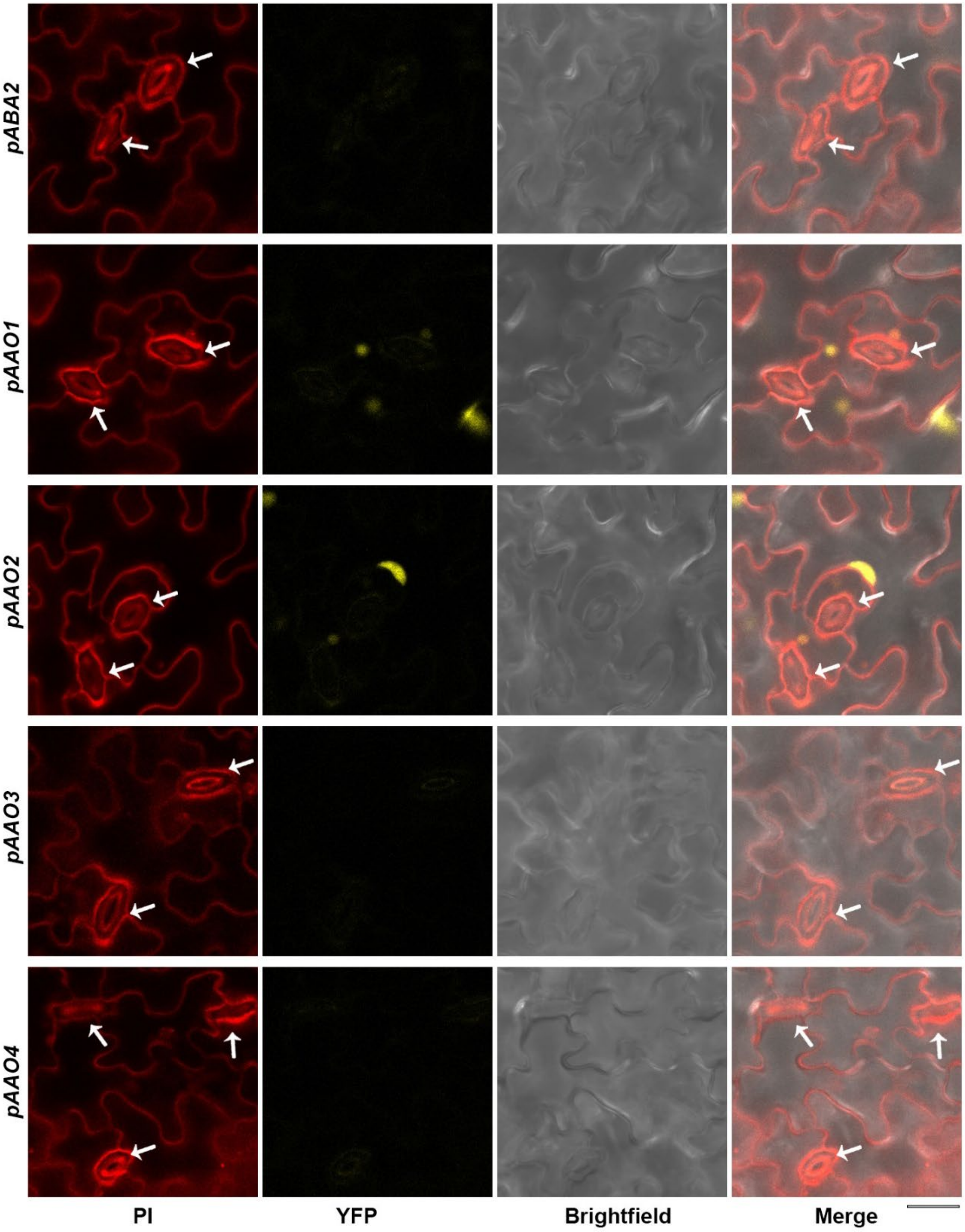


`

**Sup. Fig. S14. *ABA2* and *AAO*s are not expressed in guard cells of 14-day-old leaves.** Shown are 14-day-old leaf images of *pABA2, pAAO1, pAAO2, pAAO3* and *pAAO4:NLS-YFP* plants. Scale bars = 20 µm. White arrows point at guard cells.

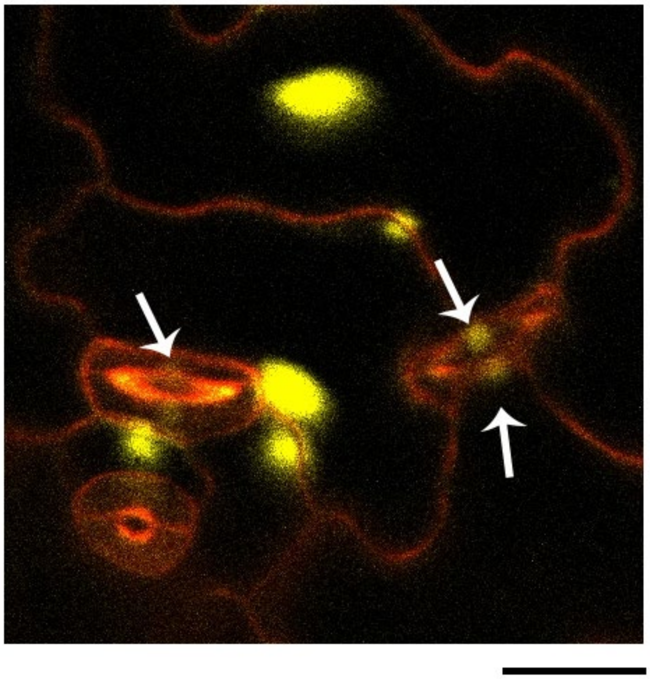

**Sup. Fig. S15. *pAAO1* shows low expression in the guard cells of 7-day-old cotyledons.** Pavement cells and guard cells image of 7-day-old cotyledons of *pAAO1:NLS-YFP*. Scale bar = 20 µm.

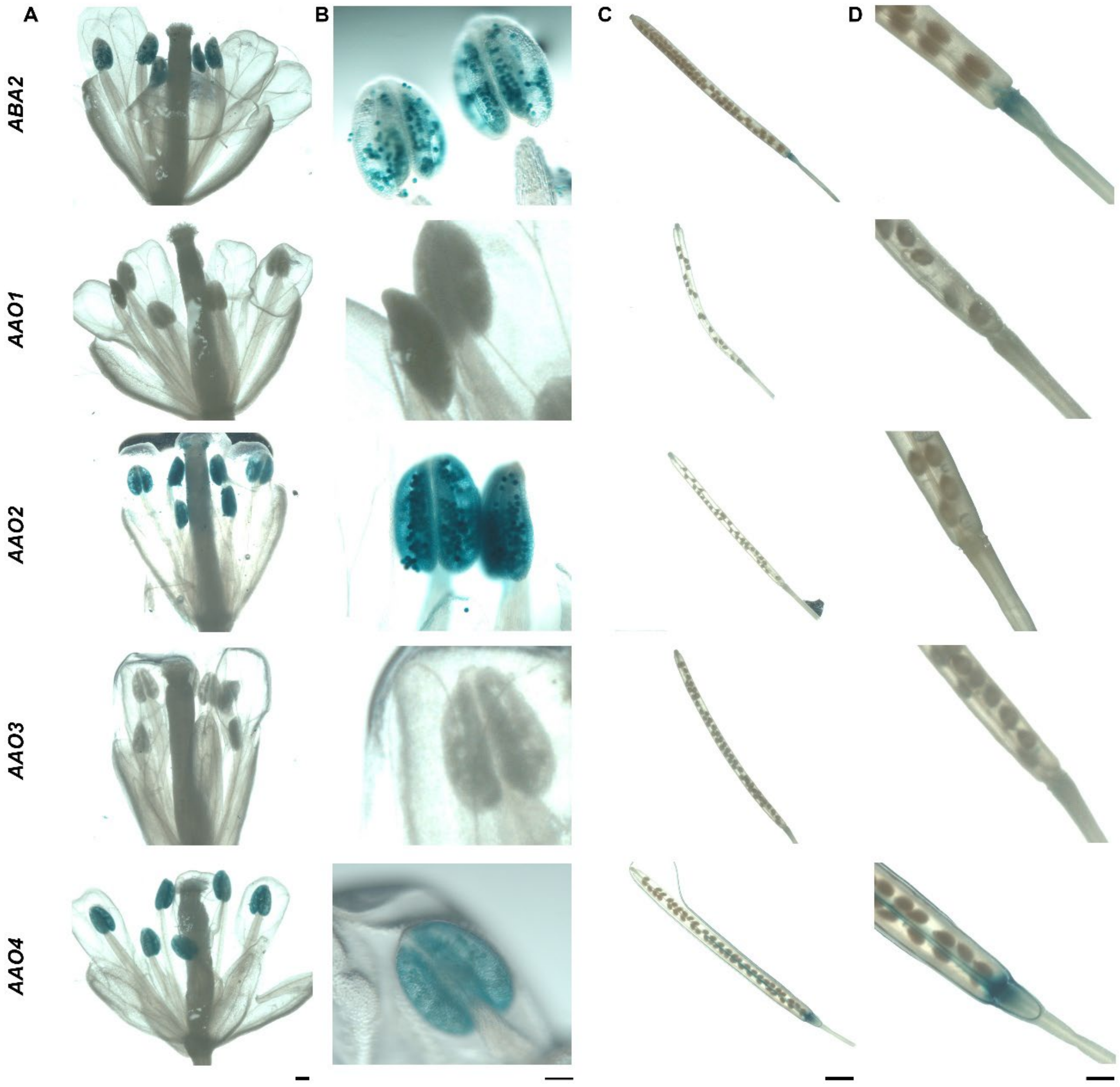


**Sup. Fig. S16. Expression pattern of *ABA2, AAO1, AAO2, AAO3, AAO4* in the flowers and siliques. (A)** Images of flowers from 42-day-old plants expressing *pABA2, AAO1, AAO2, AAO3,* and *AAO4:GUS*. Scale bar = 2 mm. **(B)** Enlargement of the flower images. Scale bar = 200 µm. **(C)** Images of siliques from 42-day-old plants expressing *pABA2, AAO1, AAO2, AAO3,* and *AAO4:GUS*. Scale bar = 500 µm. **(D)** Enlargement of the silique images. Scale bar = 100 µM.

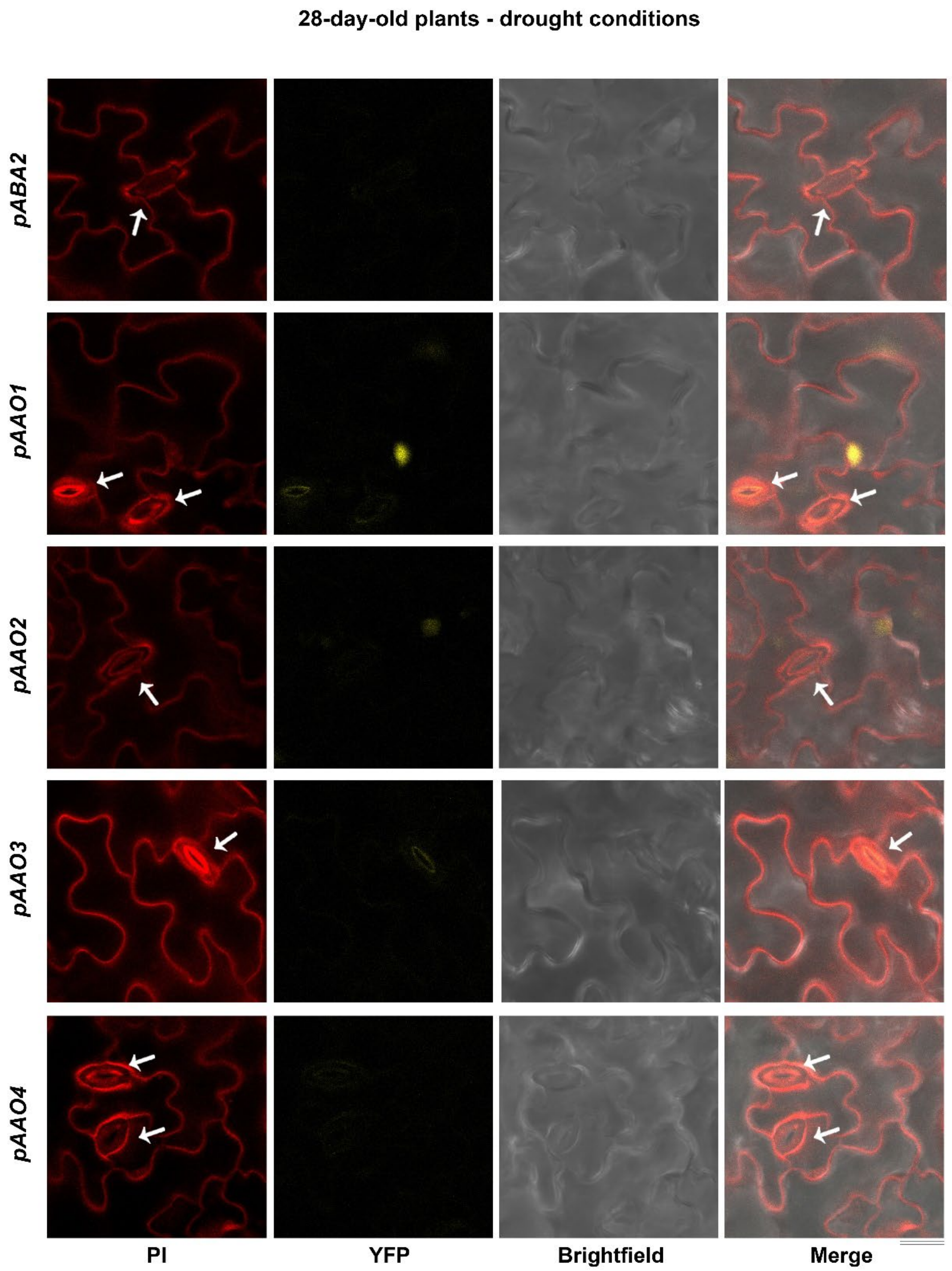


**Sup. Fig. S17. *ABA2* and *AAO*s expression is not detected in guard cells of 28-day-old leaves following water stress.** Shown are 28-day-old leaf images of *pABA2, pAAO1, pAAO2, pAAO3* and *pAAO4:NLS-YFP* plants, subjected to drought (no irrigation) for 10 days. Scale bars = 20 µm. White arrows point to guard cells.

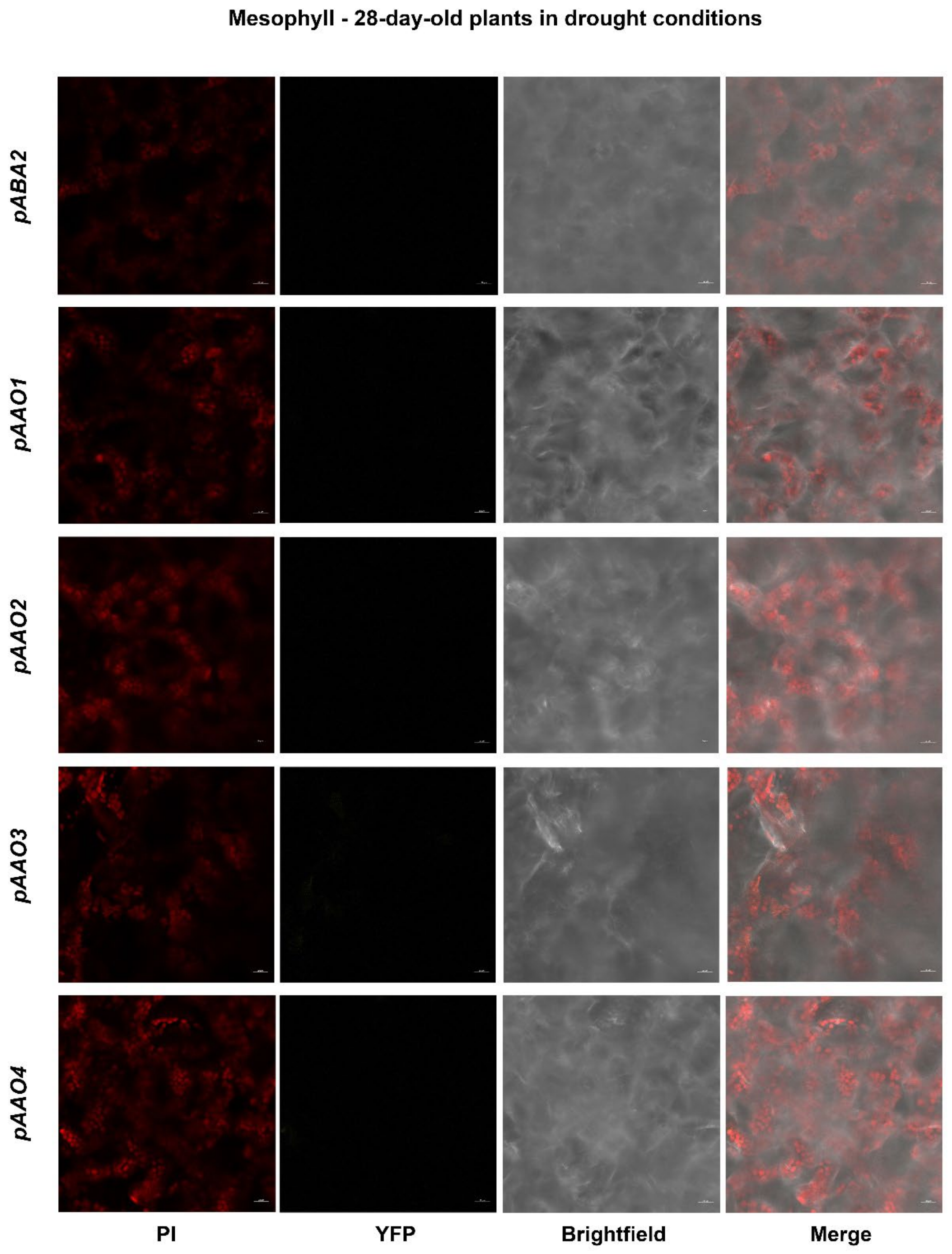


**Sup. Fig. S18.** ***ABA2*** **and** ***AAO*****s expression is not detected in mesophyll cells of 28-day-old leaves during water stress.** Shown are 28-day-old leaf images of *pABA2, pAAO1, pAAO2, pAAO3* and *pAAO4:NLS-YFP* plants, subjected to drought (no irrigation) for 10 days. Scale bars = 20 µm.

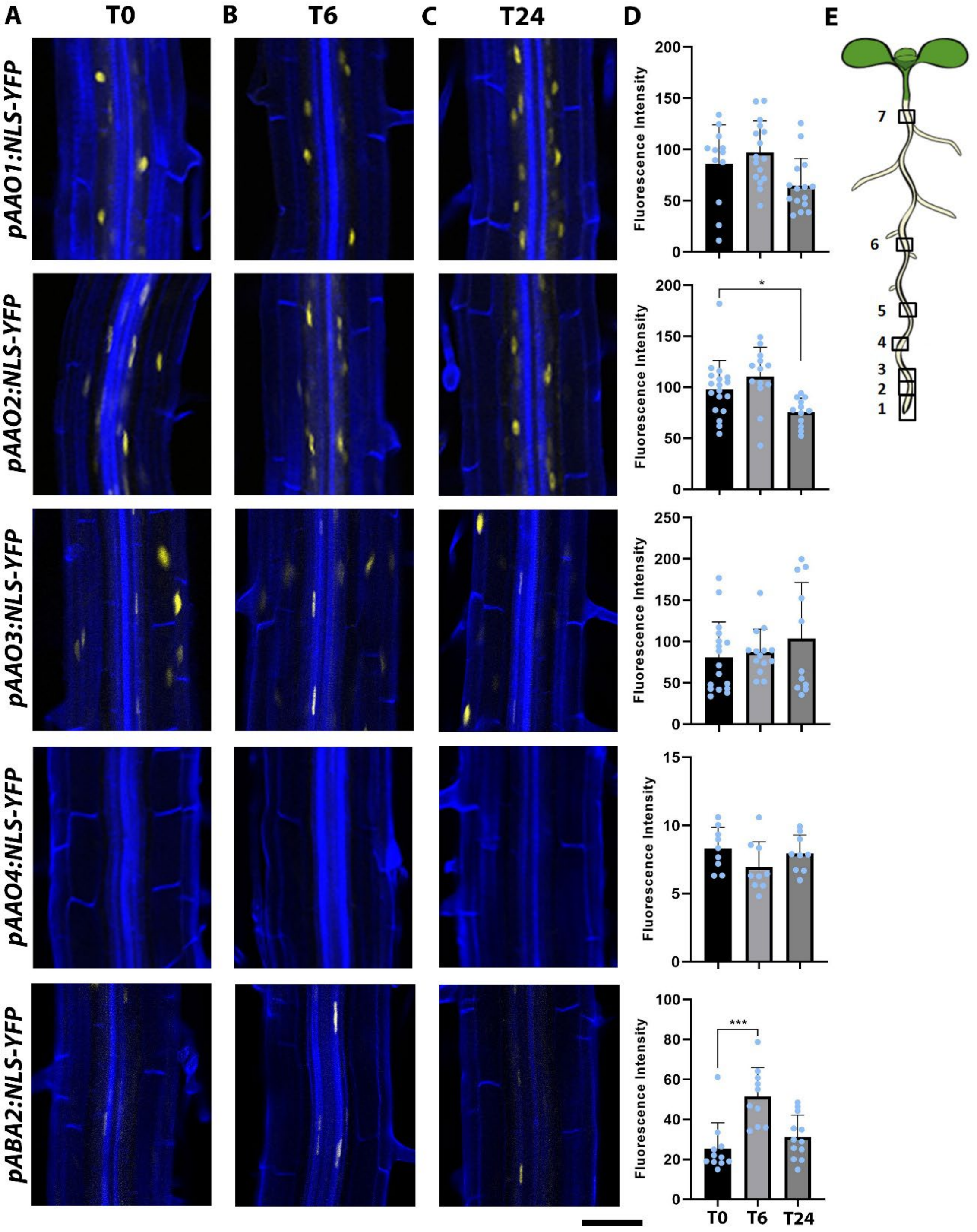


**Sup. Fig. S19. Expression of *ABA2, AAO1, AAO2, AAO3,* and *AAO4* in roots does not change in response to 24-hour salt treatment.** Shown are root images **(A-C)** and quantification **(D)** of 7-day-old *pABA2 and pAAOs:NLS-YFP* plants, before (T0; time point 0) **(A)**, after 6 hours (T6) **(B)**, and after 24 hours (T24) **(C)** of salt stress treatments (100 mM NaCl). Scale bar = 50 µm. Fluorescence intensity was measured in all fluorescent tissues, shown are averages (±SD), (3-5 cells sampled from each root, n ≥ 9), and two-tailed t-test ($P = 0.05$). **(E)** An illustration of Arabidopsis root, images, and measurements were taken at point 4.

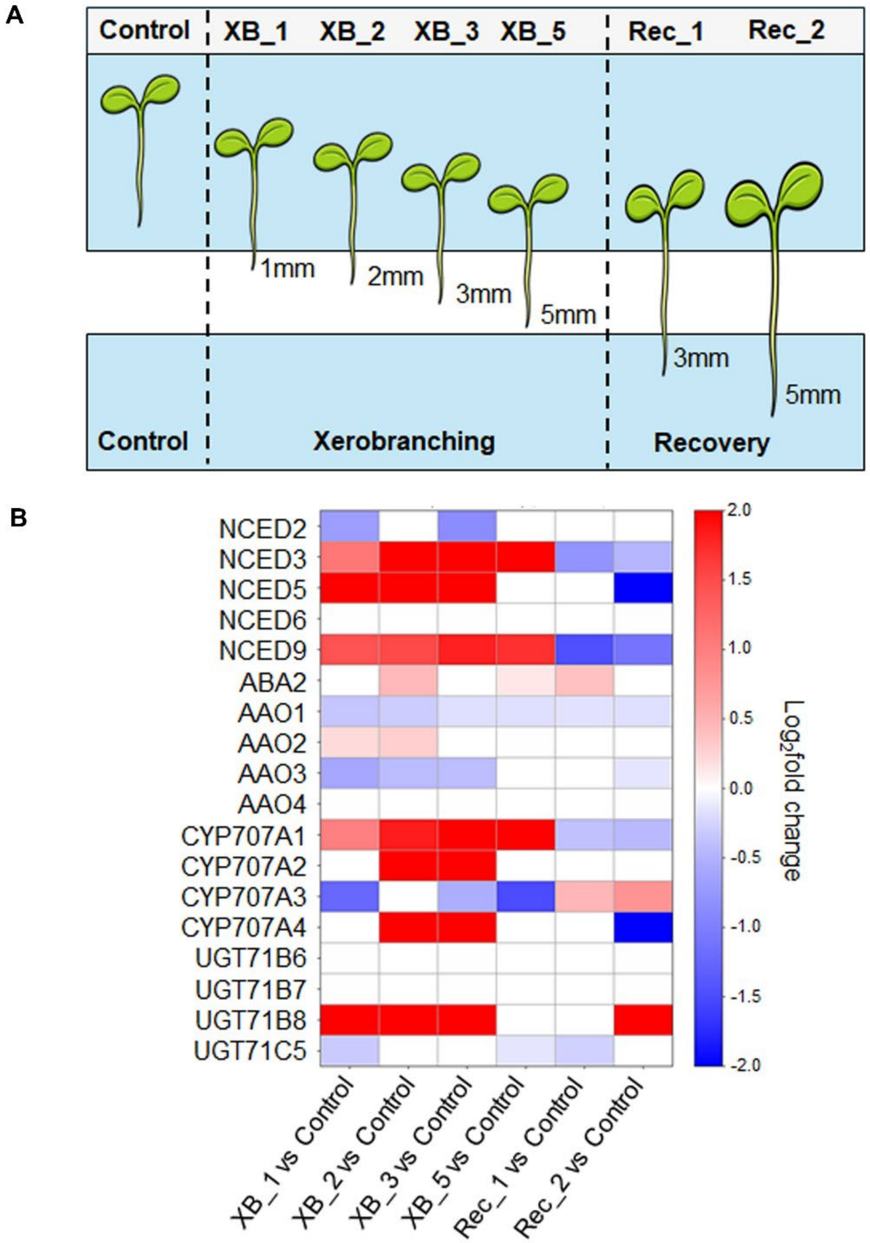


**Sup. Fig. S20. Dynamics of ABA metabolism genes under xerobranching conditions. (A)** Schematic representation of the sampling strategy of Arabidopsis root tips for gene expression analysis. Five-day-old seedlings were subjected to air-gap conditions (xerobranching), followed by recovery (reconnection with agar). XB_1, XB_2, XB_3, and XB_5 indicate root tips that grew1, 2, 3, or 5 mm, respectively, within the air gap. Rec_1 and Rec_2 correspond to roots that grew 3 mm and 5 mm, respectively, after reconnection to agar (recovery conditions). For transcriptomic analysis, the apical 2 mm of the root tips were harvested. **(B)** Heat map showing relative expression levels ($\log_2$ fold change) of ABA metabolism genes during xerobranching and subsequent recovery conditions. Fold changes were calculated relative to control conditions. Only statistically significant changes ($p \leq 0.05$) have been displayed. Empty cells indicate either undetectable expression or non-significant changes.

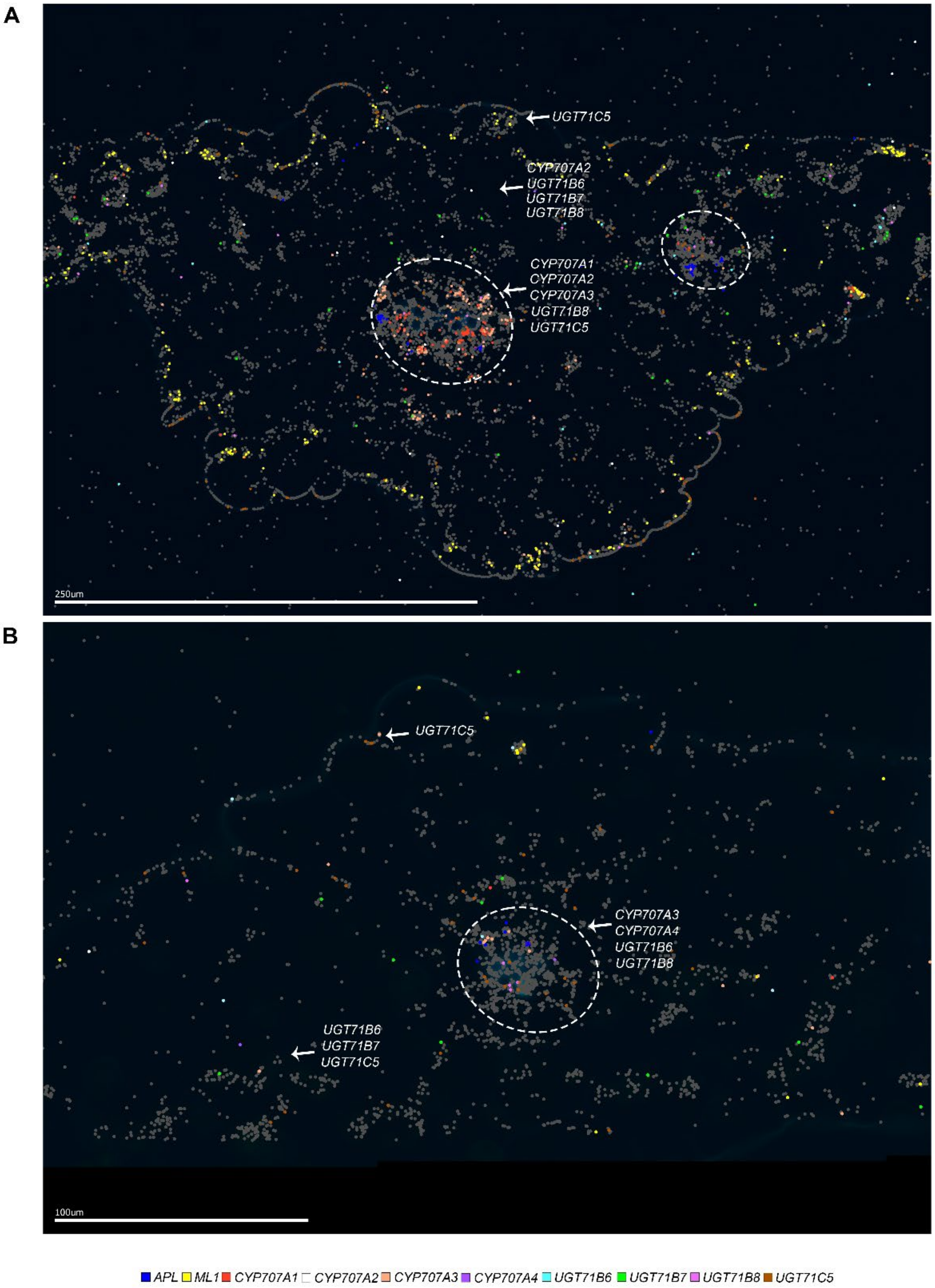


**Sup. Fig. S21. *CYP707As* and *UGTs* expressions are similar before and after drought stress. (A, B)** Leaf spatial transcriptomics for *CYP707A* and *UGT* genes under watered conditions **(A)** and drought stress conditions **(B)**. Scale bars = 250, 100 µm, respectively. Marker genes: *ML1* epidermis, *APL* phloem companion cells. White dashed circles represent the vasculature. Genes written in white indicate the tissue in which they are expressed**.**

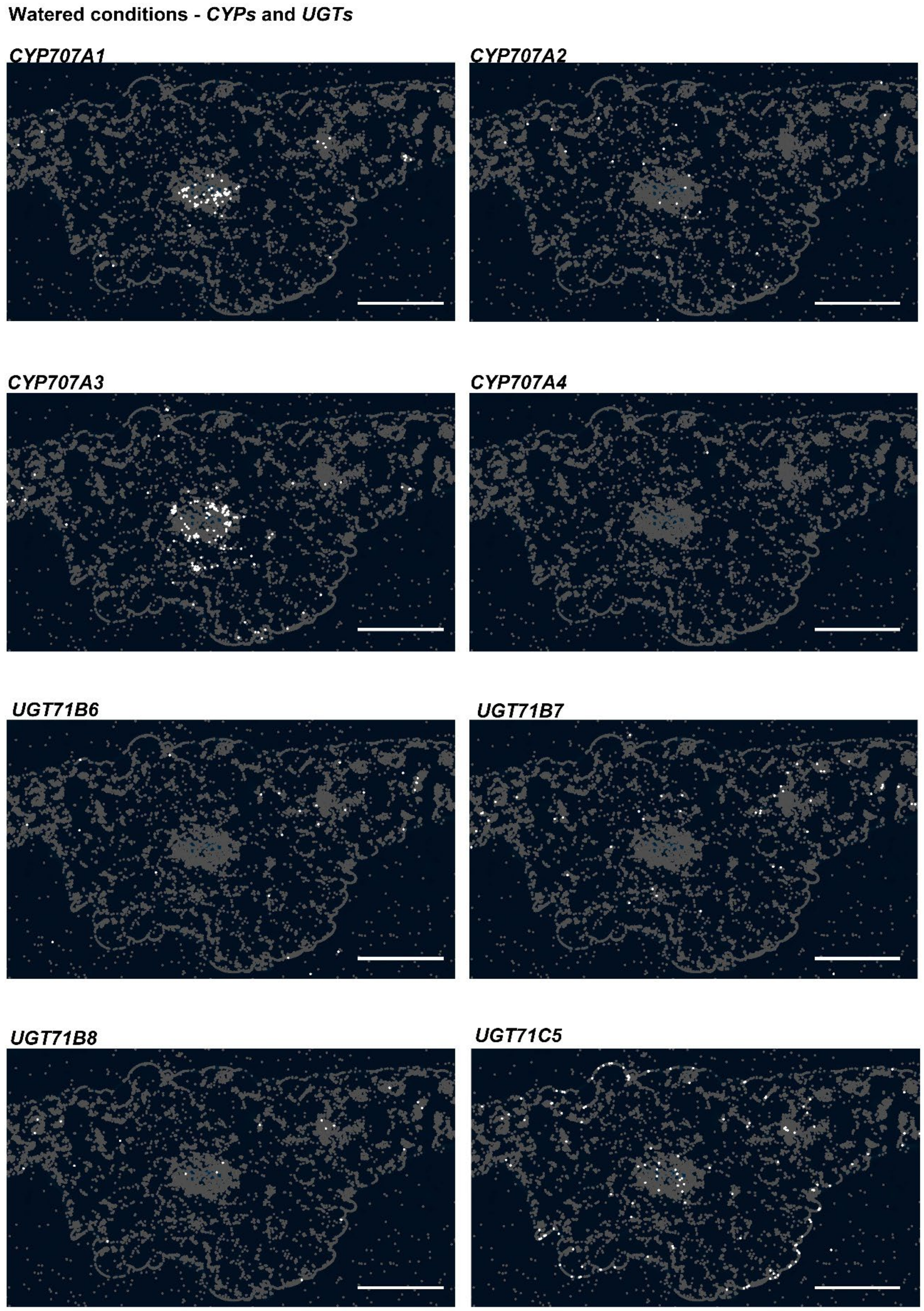


**Sup. Fig. S22. *CYP707As* and *UGTs* are mainly expressed in the leaf vasculature, mesophyll and epidermis.** Leaf spatial transcriptomics for *CYP707A* and *UGT* genes. Scale bar = 250 µm.

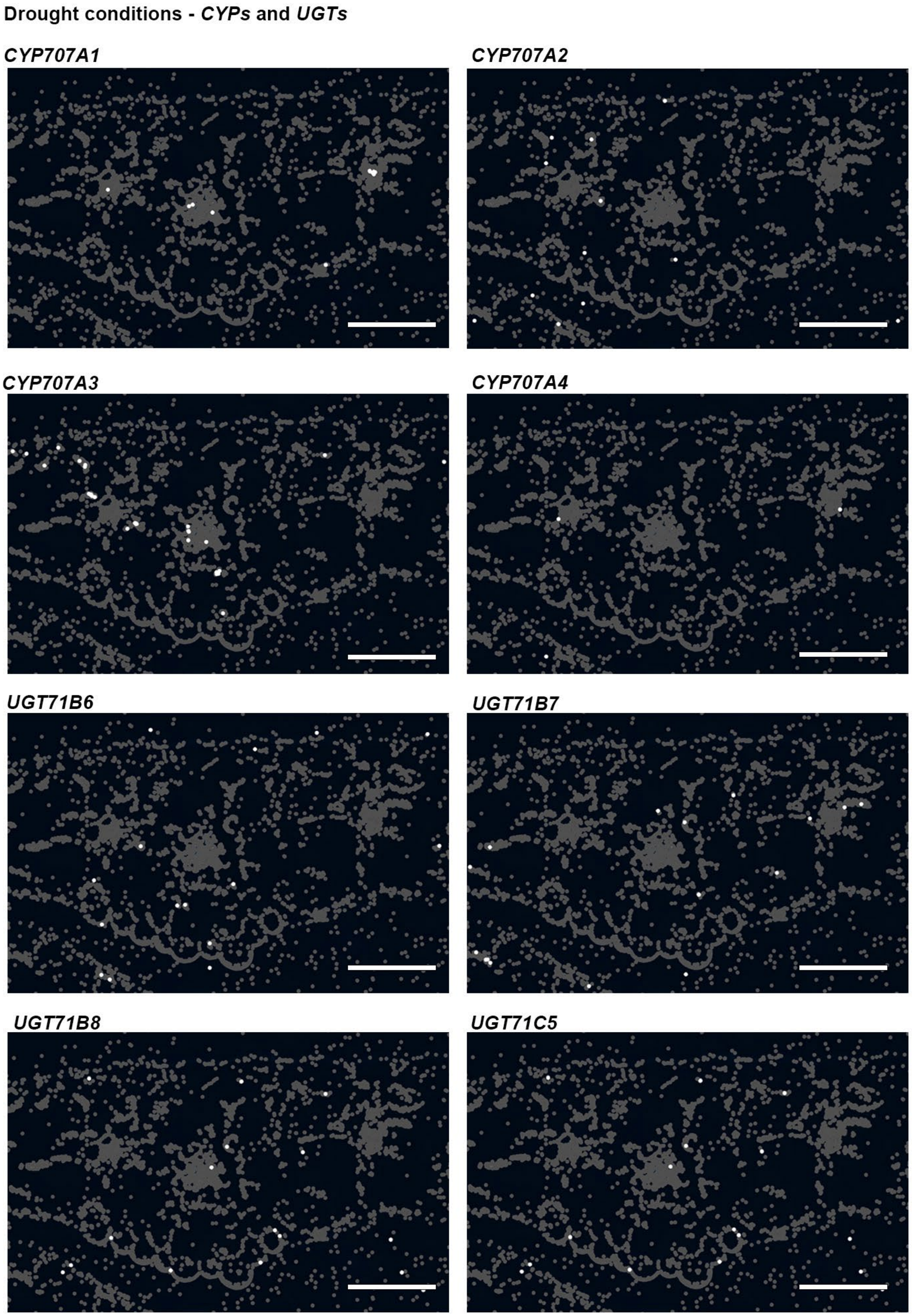


**Sup. Fig. S23.** ***CYP707As*** **and** ***UGTs*** **are mainly expressed in the leaf vasculature, mesophyll and epidermis after drought stress.** Leaf spatial transcriptomics for *CYP707A* and *UGT* genes after 10 days without irrigation. Marker genes: *ML1* epidermis, *APL* phloem companion cells. Scale bar = 100 µm.

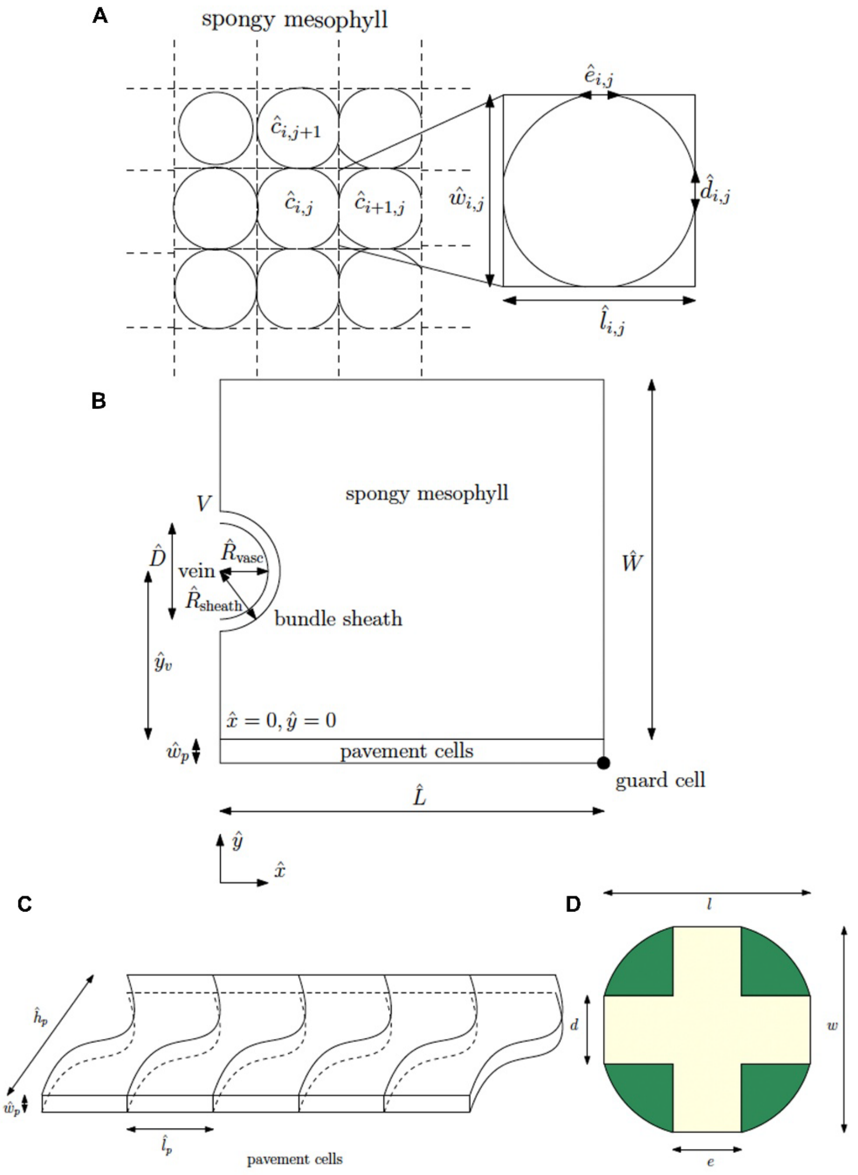


**Sup. Fig. S24. Schematics to aid the mathematical model description and illustrate the specified model parameters. (A)** Schematic of the cellular geometries within the spongy mesophyll. **(B)** Schematic of the leaf cross-section, here showing the model parameter values. **(C)** Schematic of the epidermal layer showing the pavement cell geometries. **(D)** Schematic of a cell geometry in the spongy mesophyll showing prescribed lengths l, w, d and e, and colored regions used in deriving the relationship between the contact lengths and porosity in Supp Appendix Text section 6.

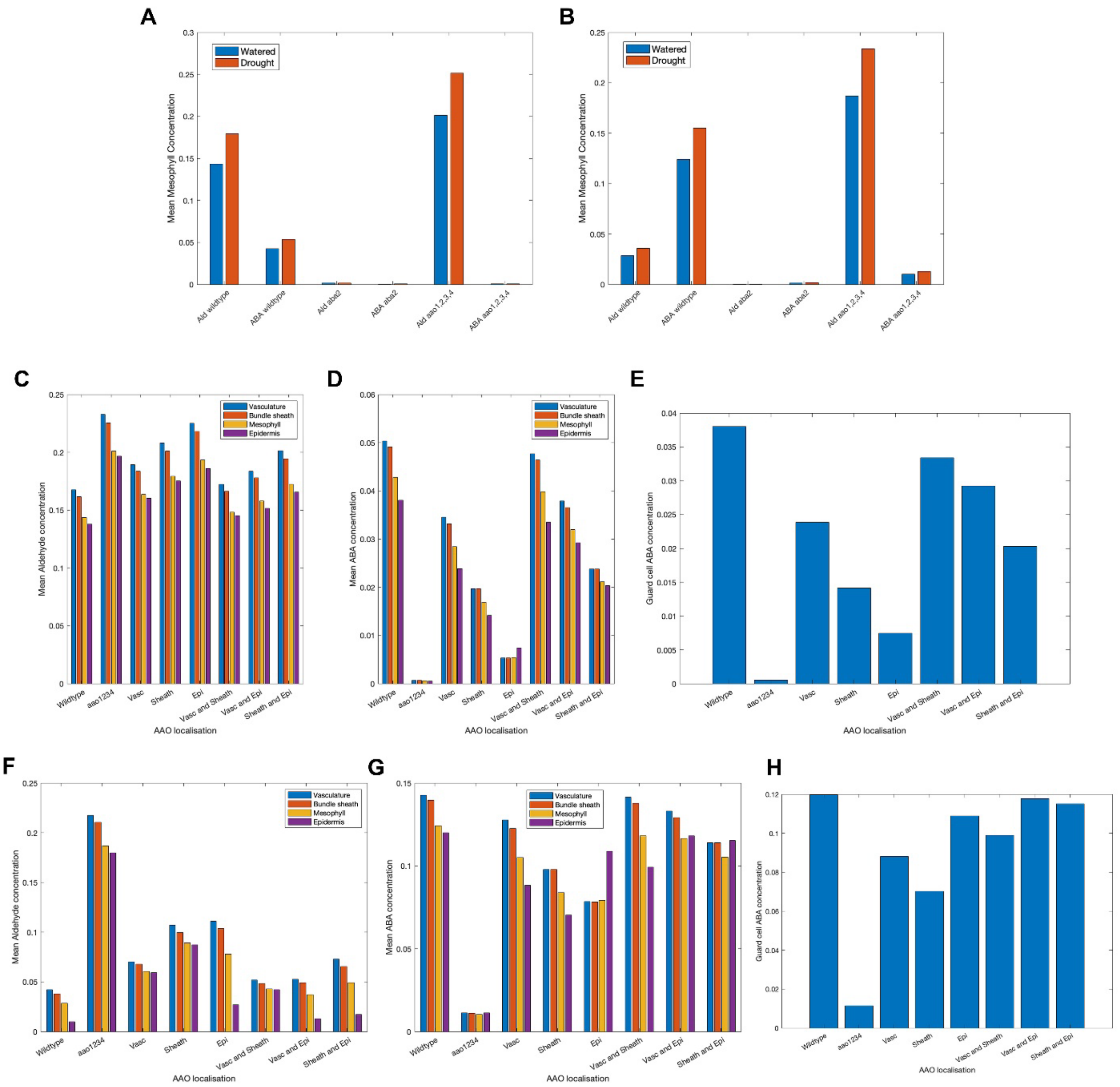

**Sup. Fig. S25. Model predictions are robust to chosen AAO-mediated conversion rates. (A-B)** Model predictions of the mean mesophyll ABA concentrations across different genotypes under well-watered (blue) and drought (orange) conditions. **(C-E)** Predicted AB-ald **(C)**, ABA **(D)** and guard cell-ABA **(E)** concentrations when AAOs are localized in specific tissues. Vas. = Vasculature, B.sheath = Bundle sheath, Ep. = Epidermis. Panels **(A,C-E)** use kAAO=1 and panels **(B,F-H)** use [AAO-epi]=1.

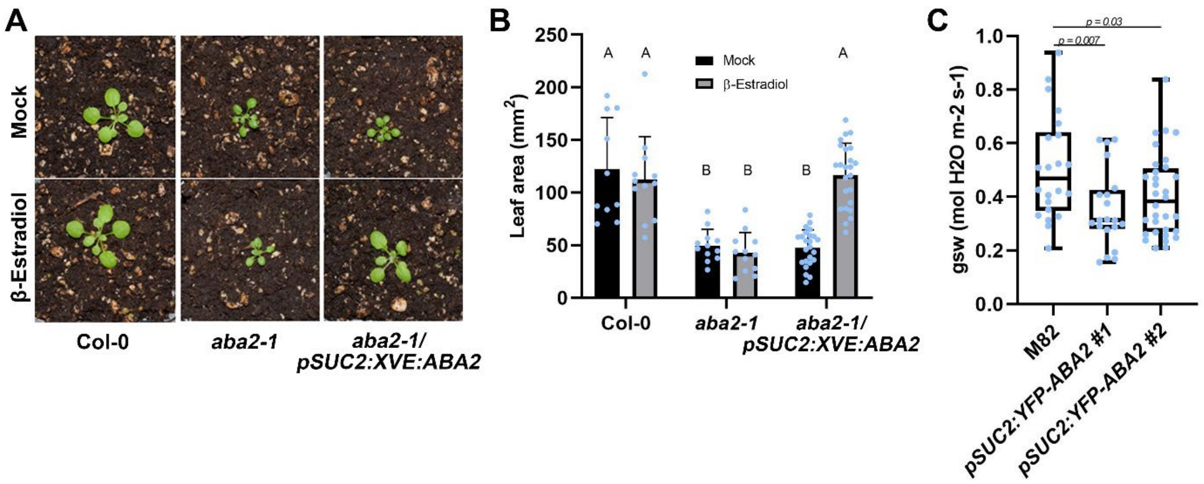


**Sup. Fig. S26. Tissue-specific ABA synthesis is sufficient to rescue ABA deficiency with respect to stomatal aperture. (A, B)** Representative images **(A)** and quantification **(B)** of 21-day-old *aba2-1/pSUC2:XVE:ABA2* plants and respective controls, with and without estradiol treatment. Shown are averages (±SD), n ≥ 8; Significance was determined using Tukey's ad-hoc statistical test (treatments marked with different letters are significantly different). Scale bar = 1 cm. **(C)** Stomatal transpiration measurements of 40-day-old *pSUC2:YFP-ABA2* tomato plants and respective controls gsw = stomatal conductance to water vapor. Shown are averages (±SD), n ≥ 20. Significance was determined using Student's t-test.

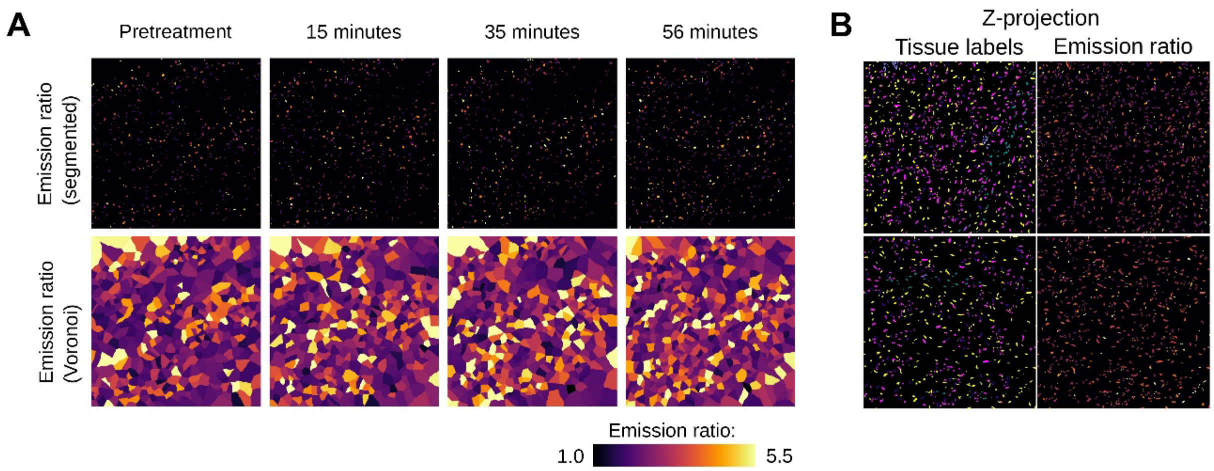


**Sup. Fig. S27. ABACUS2-400n emission ratios across leaf tissues. (A)** Maximum Z and Voronoi project of ABACUS2-400n emission ratios in Fig. 3G. **(B)** Emission ratio of nlsABACUS2–400n in different leaf tissues after a 35-minute petiole feed of 10 µM ABA or mock solution, illuminated in air. Leaves were mounted in perfluorodecalin after treatment to allow deep imaging.

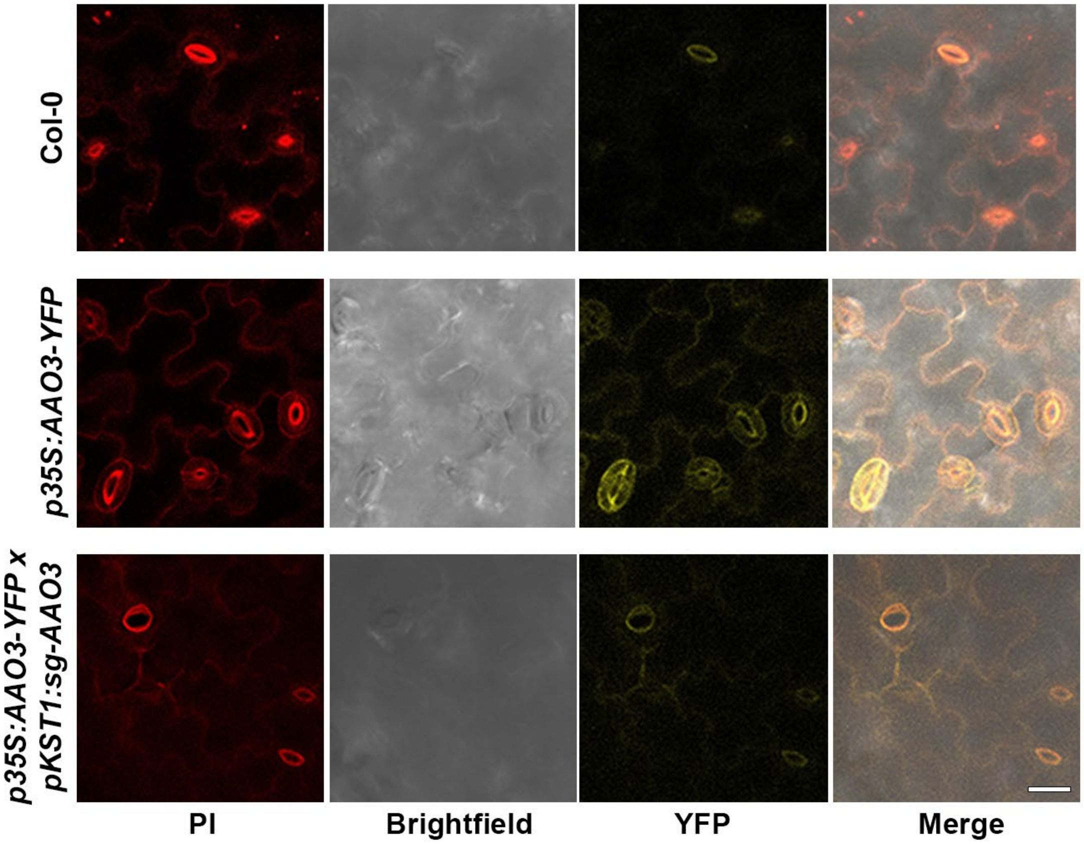


**Sup. Fig. S28. Tissue-specific knockout of AAO3 using guard cell-specific promoter, *KST1*.** Shown are images of 7-day-old *p35S:AAO3-YFP x pKST1:sg-GFP* expression and respective controls in the guard cells and pavement cells. Scale bar = 20 uM.

| | sgRNA sequence | Targeted gene/s |
|---|---|---|
| ***sg-ABA2*** | GTTCGTCTGTTCCACAAGCA | *ABA2* |
| ***sg-AAO3*** | CAGTATGCTTACTTTCCCCG | *AAO3* |
| ***2sg-4AAOs*** | GTCATCACTAGAAGAGTTGG | *AAO1,3,4* |
| | GGGACAAGGGTTATGGACAA | *AAO2,4* |

**Sup. Table S1. sgRNA sequences targeting ABA2, AAO3 and 4AAOs.** List of sgRNA sequences and respective targeted genes in this study.

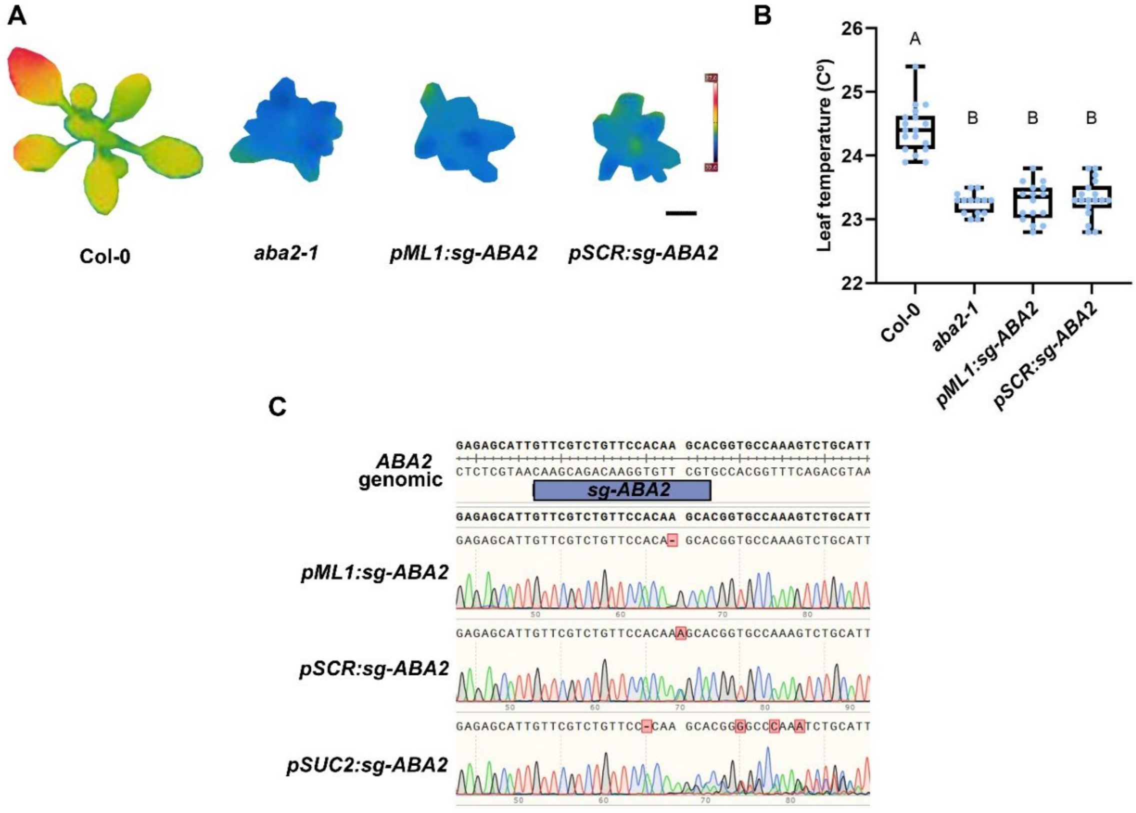


**Sup. Fig. S29*. pML1* and *SCR:Cas9* causes stable meristematic mutations. (A, B)** Shown are images **(A)** and quantification **(B)** of 24-day-old *pML1, pSCR:sg-ABA2* plants with respective controls. n ≥ 14. Scale bar = 1 cm. Significance was determined using Tukey's ad-hoc statistical test (treatments marked with different letters are significantly different). **(C)** Shown are genotyping results of the indicated lines. *ML1* and *SCR* show stable mutations, while *SUC2* shows mixed chromatogram peaks, indicating a possible cell type-specific mutation.

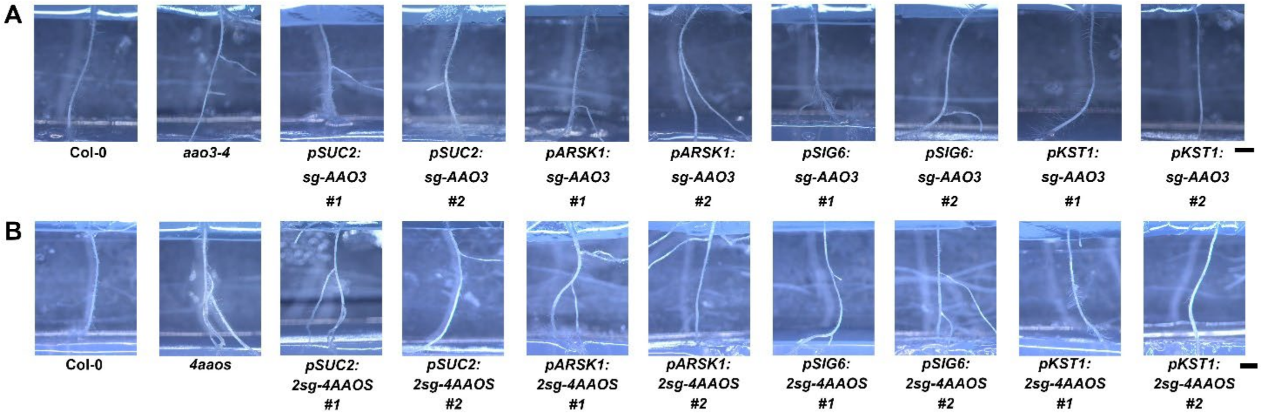


**Sup. Fig. S30. ABA2 and AAOs act from the phloem to regulate xerobranching. (A, B)** Shown are images of 13-day-old roots of cell type-CRISPR lines for *AAO3* **(A),** and *4AAOs* **(B)** with respective controls in an agar-based air-gap assay system (Air gap = ~5 mm). Scale bar = 1 mm.

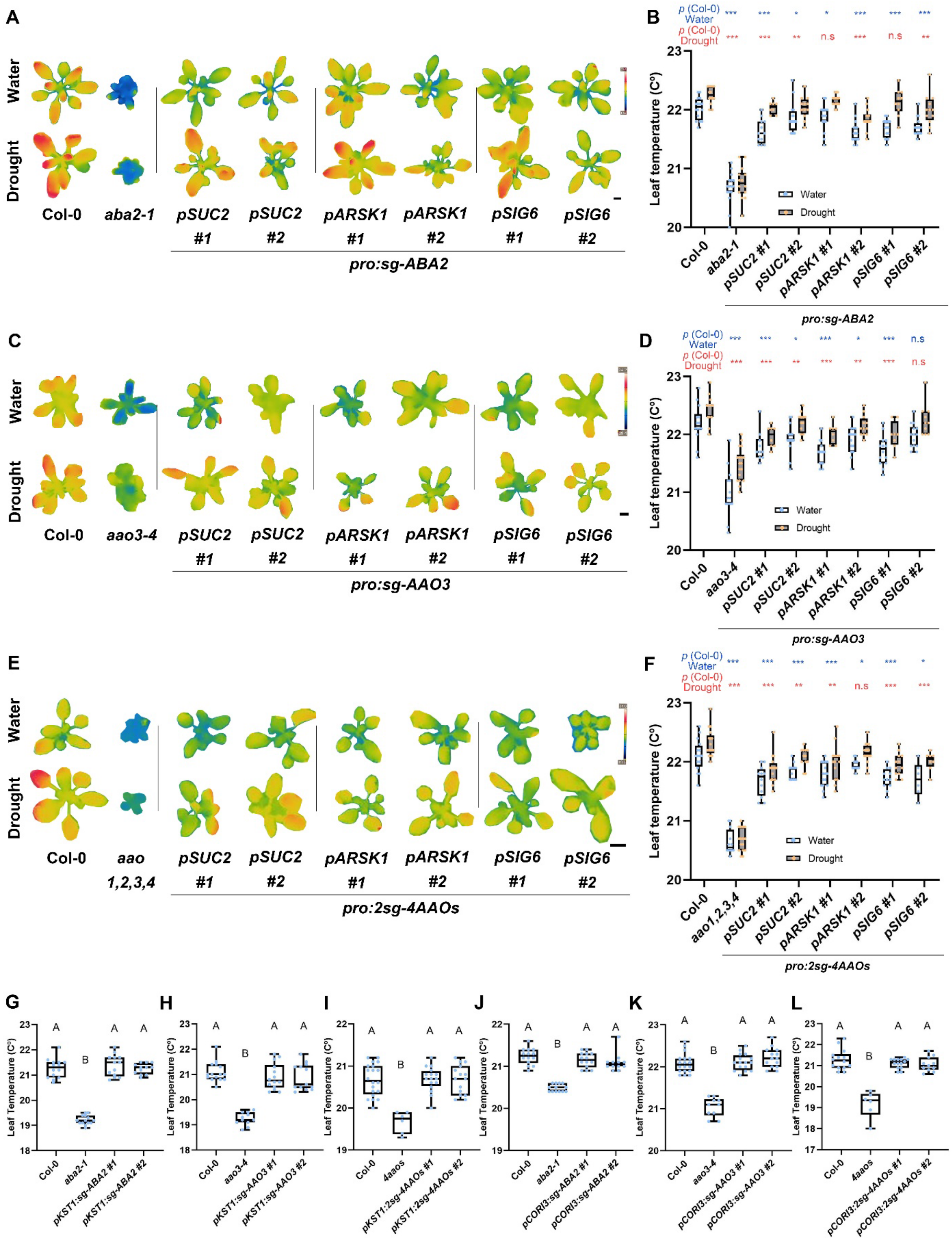
A
Water
Drought
Col-0
aba2-1
pSUC2 #1
pSUC2 #2
pARSK1 #1
pARSK1 #2
pSIG6 #1
pSIG6 #2
pro:sg-ABA2
B
p (Col-0) Water
p (Col-0) Drought
Leaf temperature (C°)
Water
Drought
pro:sg-ABA2
C
Col-0
aao3-4
pro:sg-AAO3
D
pro:sg-AAO3
E
aao 1,2,3,4
pro:2sg-4AAOs
F
aao1,2,3,4
pro:2sg-4AAOs
G
H
I
J
K
L
Leaf Temperature (C°)
pKST1:sg-ABA2 #1
pKST1:sg-ABA2 #2
pKST1:sg-AAO3 #1
pKST1:sg-AAO3 #2
4aaos
pKST1:2sg-4AAOs #1
pKST1:2sg-4AAOs #2
pCORI3:sg-ABA2 #1
pCORI3:sg-ABA2 #2
pCORI3:sg-AAO3 #1
pCORI3:sg-AAO3 #2
pCORI3:2sg-4AAOs #1
pCORI3:2sg-4AAOs #2

**Sup. Fig. S31. ABA2 and AAOs act from the phloem to regulate stomatal closure. (A-F)** Shown are thermal images **(A, C, E)** and quantification **(B, D, F)** of 28-day-old leaves of cell type-CRISPR lines targeting *ABA2* **(A, B)**, *AAO3* **(C, D)** and *4AAOs* **(E, F)** with respective controls with and without drought stress. Scale bar = 1 cm, n ≥ 6. Significance was determined using Student's t-test. * = *p value < 0.05,* ** = *0.005 < p value < 0.01,* and *** = *p value < 0.005*. In blue, *p values* of water treatment, in red, *p values* of drought stress. **(G-I)** Leaf temperature quantification of 28-day-old plants knocked out specifically in the guard cells (using promoter *KST1*) in *ABA2* **(G)**, *AAO3* **(H)**, and *4AAOs* **(I)**. **(J-L)** Leaf temperature quantification of 28-day-old plants knocked out specifically in the spongy mesophyll (using promoter *CORI3*) in *ABA2* **(J)**, *AAO3* **(K)**, and *4AAOs* **(L)**. Significance was determined using Tukey's ad-hoc statistical test (treatments marked with different letters are significantly different), n ≥ 6.

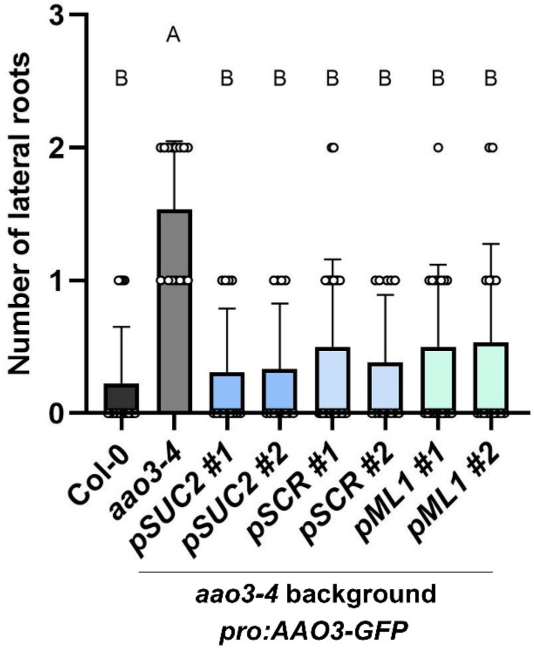


**Sup. Fig. S32. Expression of *AAO3* in different cell types rescues *aao3-4* mutant phenotype.** Quantification of 13-day-old *pSUC2, pSCR, and pML1:AAO3-GFP* on the background of *aao3-4* roots with respective controls in an agar-based air-gap assay system (Air gap = ~5 mm). n ≥ 12. Significance was determined using Tukey's ad-hoc statistical test (treatments marked with different letters are significantly different).

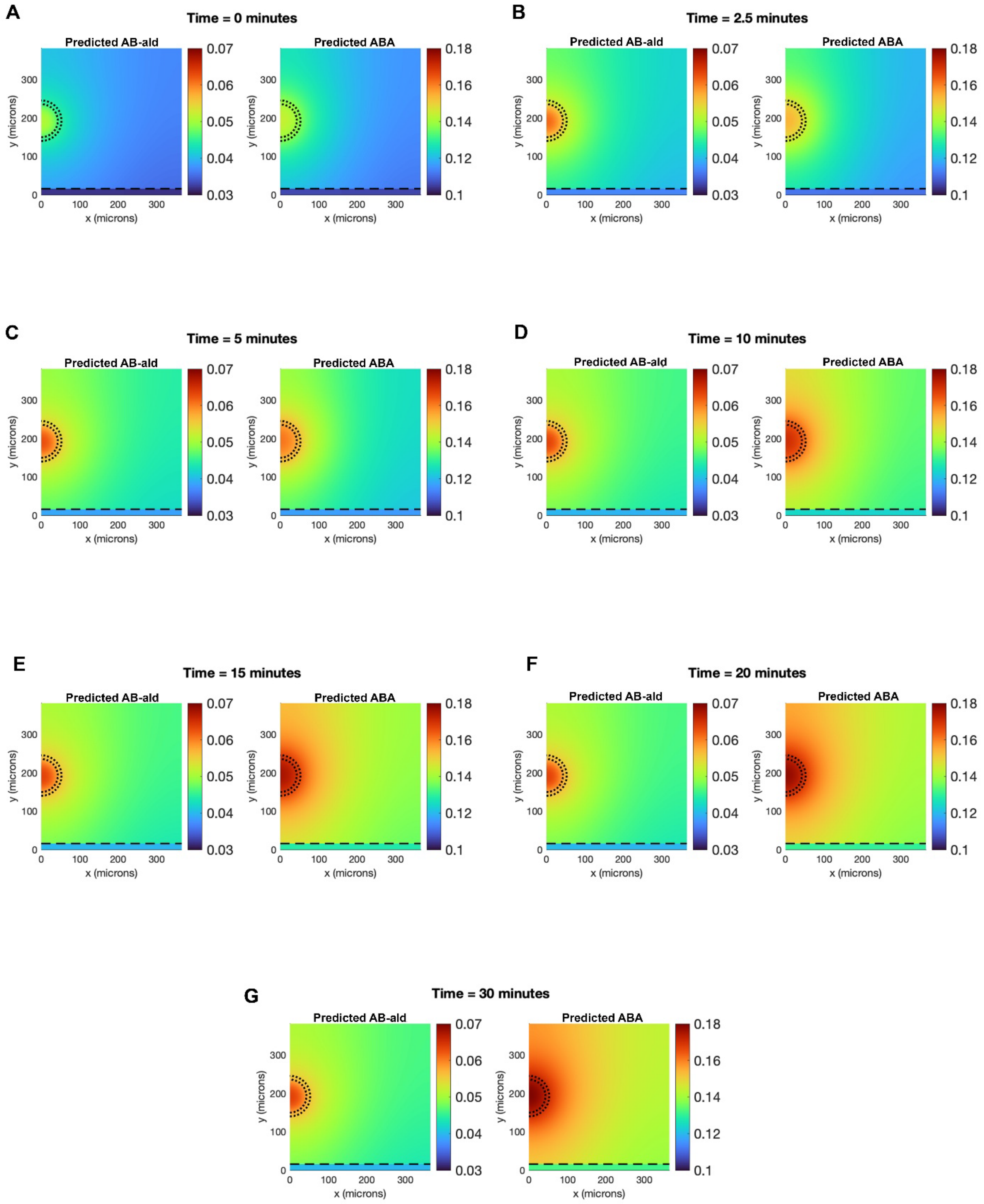


**Sup. Fig. S33. Predicted evolution of AB-ald and ABA distribution in response to water stress.** The model assumes that water stress induces a 25% increase in AB-Ald production in the vasculature. **(A)** t = 0, **(B)** t = 2.5 minutes, **(C)** t = 5 minutes, **(D)** t = 10 minutes **(E)** t = 15 minutes, **(F)** t = 20 minutes, **(G)** t = 30minutes.

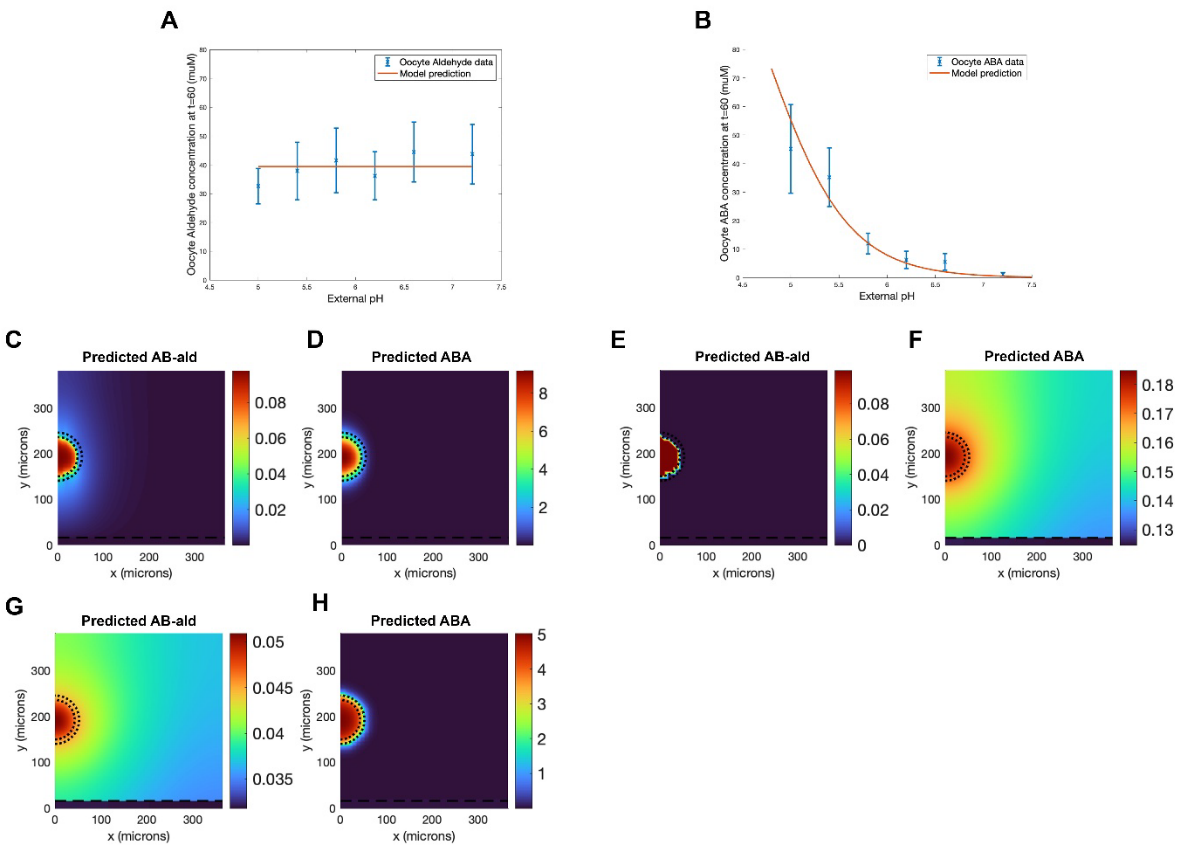


**Sup. Fig. S34. Model predictions on the effect of the permeabilities on the hormone concentrations. (A)** Predicted aldehyde concentration from the oocyte model is in good agreement with the experimental measurements, provided we use the estimated Aldehyde membrane permeability value of 0.017 microns/sec. (see section 7 of the Supplementary Appendix text for further details) **(B)** Predicted ABA concentrations from the oocyte model are in good agreement with the experimental measurements, provided we use the estimated protonated ABA membrane permeability value of 0.10 microns/sec. **(C)** Predicted aldehyde distribution in the leaf cross-section using the passive membrane permeability values estimated from the oocyte data. **(D)** Predicted ABA distribution in the leaf cross-section using the passive membrane permeability values estimated from the oocyte data. **(E)** Predicted Aldehyde distribution in the leaf cross-section, assuming that only ABA moves between adjacent cells. **(F)** Predicted ABA distribution in the leaf cross-section, assuming that only ABA moves between adjacent cells. **(G)** Predicted aldehyde distribution in the leaf cross-section, assuming that only aldehyde moves between adjacent cells. **(H)** Predicted ABA distribution in the leaf cross-section, assuming that only aldehyde moves between adjacent cells.

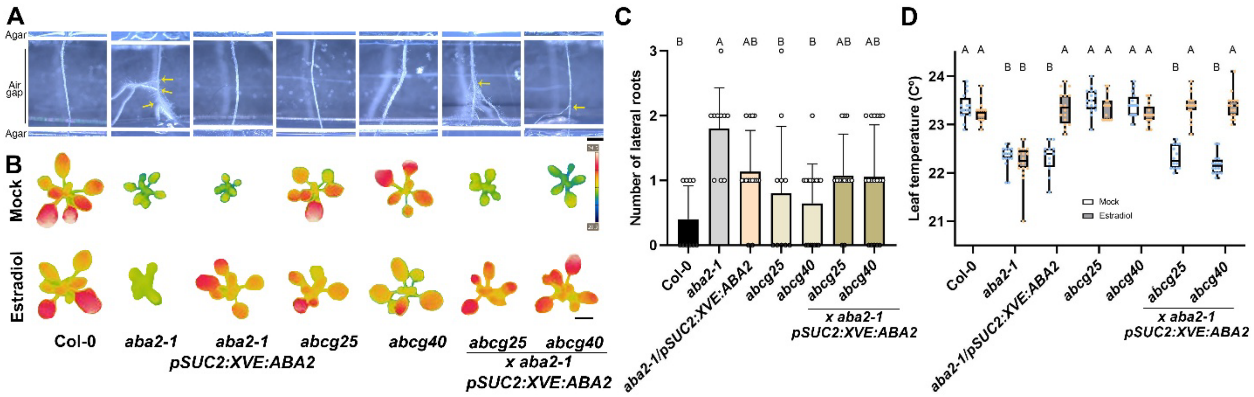


**Sup. Fig. S359.** ***ABCG25*** **and** ***ABCG40*** **are not showing significant xerobranching or stomatal aperture phenotypes. (A, C)** Representative root images **(A)** and quantification **(C)** of 13-day-old *abcg25* and *abcg40* with respective controls in an agar-based air-gap assay system (Air gap = ~5 mm). Yellow arrows indicate lateral roots. Scale bar = 1 mm. n ≥ 10, Significance was determined using Tukey's ad-hoc statistical test (treatments marked with different letters are significantly different). **(B, D)** Representative thermal images **(B)** and quantification **(D)** of 21-day-old *abcg25, abcg40* and respective controls. Plants were either under mock treatment or under 5 µM estradiol treatment. Scale bar = 1 cm. n ≥ 12, Significance was determined using Tukey's ad-hoc statistical test (treatments marked with different letters are significantly different).

| | Primer Fw | Primer Rev | Restriction enzyme | Final vector |
|---|---|---|---|---|
| ***YFP-ABA2*** | acGGTCTCaATTGatggtgagcaagggc | caGGTCTCtAAACtcatctgaagactttaaaggagt | BsaI | *pro:YFP-ABA2* |
| ***pABA2*** | GATGGCGCGCCcacatatcatcaattcatcatgt | CGATGGATCCaatagctttagctccttagatct | AscI, BamHI | *pABA2:YFP-ABA2* |
| ***pAAO3*** | tgcctgcaggaggtgttatgaatgaattgatg | CGGATCCttttttcccagagcagtga | SbfI, BamHI | *pAAO3:AAO3-GFP* |
| ***NLS-YFP*** | acGGTCTCaATTGCTCGACCCGACATGGAG | caGGTCTCtAAACtcagtacagctcgtccatgc | BsaI | *pAAO3:NLS-YFP* |
| ***GUS*** | acGGTCTCaATTGatgttacgtcctgtagaaaccc | aaGGTCTCtAAACtcattgtttgcctccctgc | BsaI | *pAAO3:GUS* |
| ***ABA2* CDS** | CACCATGTCAACGAACACTGAATCTTCT | tcatctgaagactttaaaggagtg | | *pSUC2:XVE:ABA2* |
| ***pSUC2*** | cattcctgcaggggcgcgccttcaagg | CTCGcctcgaggggtaccatttgacaaaccaagaaagtaaga | KpnI, PspXI | *pSUC2:AAO3-GFP* |
| ***pML1*** | CAGggtaccagtttctaaaatgtgctaaaattc | CCGctcgagctaaccggtggattcagggag | SbfI, PspXI | *pML1:AAO3-GFP* |
| ***pSCR*** | catCCTGCAGGgattgtgatcctc | catCTCGAGggagattgaagggttg | SbfI, XhoI | *pSCR:AAO3-GFP* |

**Sup. Table S2. Primers used in this study.**

| Gene | T-DNA line | Primer Fw | Primer Rev | Primer mid |
|---|---|---|---|---|
| *aao1-2* | SALK_069221 | AGCAGCTCGAGTCAAGAACAG | TGCAATATCTGCATGCTTTTG | ATTTTGCCGATTTCGGAAC |
| *aao2-1* | SALK_104895 | ACTGCATGGGAGTGTCTTTTG | GAGGTTTTGGAGGGAAATCTG | |
| *aao3-4* | SALK_072361 | TTCTATTGGAAATGCATTGCC | TAAAACATCGGATGAACCTCG | |
| *aao4-2* | SALK_057531 | ATGCTCCATGTAGACAATGGG | AATTAGTTGTTGGCAACACGG | |

**Sup. Table S3. T-DNA lines and primers used in this study.**

| **Analyte** | **Retention Time [min]** | **Q1 [*m/z*]** | **Q3 [*m/z*]** | **Fragmentor [V]** | **CE [V]** |
|---|---|---|---|---|---|
| 2-propenyl GLS (sinigrin) | 1.25 | 358.0 | 97.0$^{Qt}$ | 98 | 20 |
| $[M-H]^-$ | | 358.0 | 259.0 | 98 | 16 |
| Internal standard (IS) | | 358.0 | 75.0 | 98 | 36 |
| Caffeine (IS) | 1.95 | 195.1 | 123.0$^{Qt}$ | 86 | 36 |
| $[M+H]^+$ | | 195.1 | 83.1 | 86 | 32 |
| Internal standard (IS) | | 195.1 | 69.0 | 86 | 28 |
| ABA | 2.69 | 263.1 | 153.0$^{Qt}$ | 76 | 8 |
| $[M-H]^-$ | | 263.1 | 219.1 | 76 | 12 |
| | | 263.1 | 203.1 | 76 | 32 |
| ABA-GE | 2.42 | 425.2 | 263.1$^{Qt}$ | 91 | 8 |
| $[M-H]^-$ | | 425.2 | 305.1 | 91 | 8 |
| | | 425.2 | 219.1 | 91 | 20 |
| AB-Aldehyde | 3.11 | 249.1 | 97.1$^{Qt}$ | 68 | 12 |
| $[M+H]^+$ | | 249.1 | 231.1 | 68 | 4 |
| | | 249.1 | 121.1 | 68 | 20 |

Qt = quantifier ion, additional transitions were used for identification. Q = quadrupole. CE = collision energy.

**Table S4: MRM transitions for abscisic acid analyzed by LC-MS/MS.**

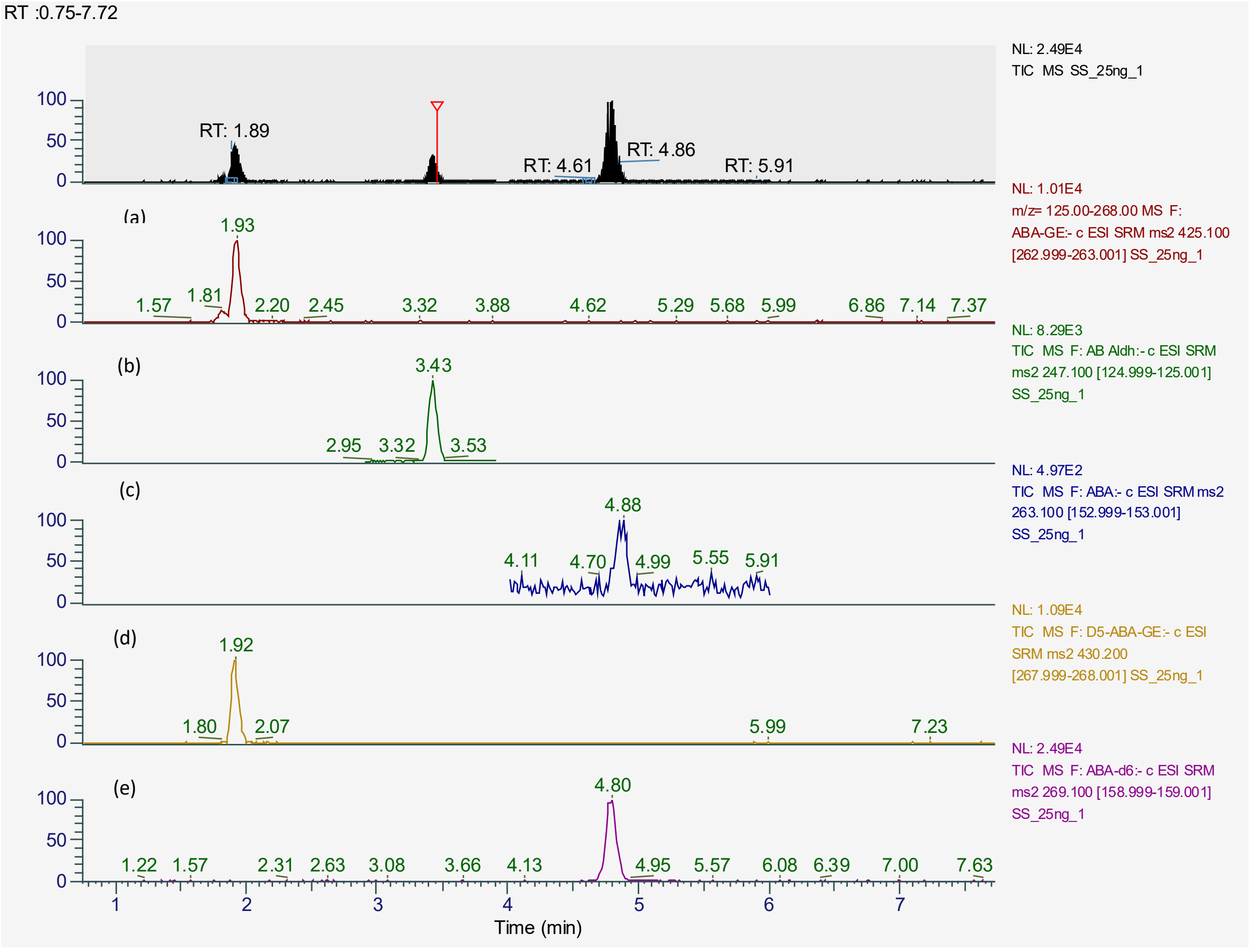


**Sup. Fig S36. Chromatogram of all standard spikes for molecule measurements.** Chromatogram of all standards spiked in solvent (20 ng) with optimized SRMs: ABA-GE (a), AB-Ald (b), ABA (c), d5-ABA-GE (d), AB-Ald (e).

| Compound | Retention Time (min) | RT Window (min) | Polarity | Precursor (m/z) | Product (m/z) | Collision Energy (V) | RF Lens (V) | Dwell Time (ms) | Use Ion |
|---|---|---|---|---|---|---|---|---|---|
| **ABA-d6** | 5 | 10 | Negative | 269.1 | 159 | 13 | 68 | 120 | Quantifier |
| **ABA-GE** | 5 | 10 | Negative | 425.1 | 153.1 | 22 | 74 | 120 | Qualifier |
| **ABA-GE** | 5 | 10 | Negative | 425.1 | 263 | 11 | 74 | 120 | Quantifier |
| **D5-ABA-GE** | 5 | 10 | Negative | 430.2 | 268 | 13 | 64 | 120 | Quantifier |
| **AB-Ald** | 3.4 | 1 | Negative | 247.1 | 125 | 17 | 85 | 120 | Quantifier |
| **ABA** | 5 | 2 | Negative | 263.1 | 153 | 11 | 78 | 120 | Quantifier |
| **ABA** | 5 | 2 | Negative | 263.1 | 219 | 14 | 78 | 120 | Qualifier |

**Sup. Table S5. Optimized SRM transitions for quantifying phytohormones.**

| | Linearity range (ng/ml) | Correlation coefficient ($R^2$) | RT | LOQ, ng/ml | RSD RT,% | RSD area,% |
|---|---|---|---|---|---|---|
| AB-Ald | 1 - 2000 | 0.9996 | 3.43 | 1 | 0.219254 | 0.9-9.0 |
| ABA | 1 - 2000 | 0.9976 | 4.87 | 2 | 0.194773 | 1.9-6.1 |
| ABA-GE | 1 - 2000 | 0.9971 | 1.92 | 0.5 | 0.333919 | 4.2-6.0 |

**Sup. Table S6.  Calibration curve and linearity data for phytohormones.**
